# Potential energy landscapes reveal the information-theoretic nature of the epigenome

**DOI:** 10.1101/059766

**Authors:** Garrett Jenkinson, Elisabet Pujadas, John Goutsias, Andrew P. Feinberg

## Abstract

Epigenetics is defined as genomic modifications carrying information independent of DNA sequence heritable through cell division. In 1940, Waddington coined the term “epigenetic landscape” as a metaphor for pluripotency and differentiation, but epigenetic potential energy landscapes have not yet been rigorously defined. Using well-grounded biological assumptions and principles of statistical physics and information theory, we derive potential energy landscapes from whole genome bisulfite sequencing data that allow us to quantify methylation stochasticity genome-wide and discern epigenetic differences using Shannon’s entropy and the Jensen-Shannon distance. We discover a “developmental wheel” of germ cell lineages and an association between entropy and chromatin structure. Viewing methylation maintenance as a communications system, we introduce methylation channels and show that higher-order chromatin organization can be predicted from their informational properties. Our results provide a fundamental understanding of the information-theoretic nature of the epigenome and a powerful methodology for studying its role in disease and aging.

## INTRODUCTION

The classical definition of epigenetics by Waddington is the emergence of a phenotype that can be perturbed by the environment but whose endpoints are predetermined by genes^1^. Waddington used the language of ordinary differential equations, including the notion of an “attractor”, to describe the robustness of deterministic phenotypic endpoints to environmental perturbations, which he believed to be entirely governed by DNA sequence and genes. However, a growing appreciation for the role that stochasticity and uncertainty play in development and epigenetics^2–4^ has led to relatively simple probabilistic models that take into account epigenetic uncertainty by adding a “noise” term to deterministic models^5,6^. Although some authors have recognized the importance of entropy in DNA methylation, it has so far been defined in an *ad hoc* manner with limited resolution and requiring extensive cell culture expansion and even molecular tagging for its measurement^3,4,7^. Here, we have taken a foundational approach to understanding the nature of epigenetic information by using principles of statistical physics and information theory to organically incorporate stochasticity into the mathematical framework and applying it on primary whole genome bisulfite sequencing (WGBS) datasets. The results allow us to conceptually combine “hard-wired” mechanistic principles of epigenetic biology with the Ising model of statistical physics and, in contrast to metaphorical “Waddingtonian” landscapes, rigorously derive epigenetic potential energy landscapes that can be computed genome-wide. These landscapes encapsulate the higher-order statistical behavior of methylation in a biologically relevant manner, and not just its mean as it has been customary. We quantify methylation uncertainty genome-wide using Shannon’s entropy and provide a powerful information-theoretic methodology for distinguishing epigenomes using the Jensen-Shannon distance between sample-specific potential energy landscapes associated with stem cells, tissue lineages and cancer. Moreover, we establish a relationship between entropy and topologically associating domains (TADs), which allows us to efficiently predict their boundaries from individual WGBS samples. In addition, we demonstrate that methylation can be subject to (non-critical) phase transition that may be associated with important biological functions, such as genomic imprinting. We also introduce methylation channels as models of DNA methylation maintenance and show that their informational properties can be effectively used to predict higher-order chromatin organization using machine learning. Lastly, we introduce a sensitivity index that quantifies the rate by which environmental or external perturbations influence methylation uncertainty along the genome, suggesting that genomic loci associated with high sensitivity are those most affected by such perturbations. This merger of epigenetic biology, statistical physics and information theory yields many fundamental insights into the relationship between information-theoretic properties of the epigenome and nuclear organization in normal development and disease, and demonstrate that we can precisely identify informational properties of individual WGBS samples and their chromatin structure, as well as their differences among tissue lineages, aging, and cancer.

## RESULTS

### Stochastic epigenetic variation and potential energy landscapes

Despite it being well known that stochastic variation is a fundamental property of the DNA methylome^2,3^, genome-wide modeling and analysis of the methylation state continues to focus on individual CpG dinucleotides and ignores statistical dependence among these sites. However, DNA methylation is correlated, at least over small distances, due to the processivity of the DNMT enzymes. Therefore, one cannot adequately analyze methylation with methods that do not take into account such correlation. To this end, and to better understand the relationship between stochastic epigenetic fluctuation and phenotypic variability, we took a different and more general path to methylation modeling and analysis by developing an information-theoretic approach based on the Ising model of statistical physics. This approach leads to a rigorous definition of a potential energy landscape which associates each methylation state with a potential that quantifies the information content of that state. Notably, the Ising model provides a natural way of modeling statistically dependent binary methylation data that is consistent with observed means and pairwise correlations.

Here, DNA methylation is viewed as a process that reliably transmits linear strings of binary (0-1) data from a cell to its progeny in a manner that is robust to intrinsic and extrinsic stochastic biochemical fluctuations. First, the methylation state within a given genomic region containing *N* CpG sites is modeled by an *N*-dimensional binary-valued random vector **X** whose n-th element *X_n_* takes value 0 or 1 depending on whether or not the *n*-th CpG site is unmethylated or methylated, respectively. Then, the potential energy landscape (PEL) of methylation is defined by

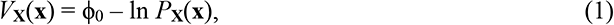

for some constant ϕ_0_, where ***P*_x_**(**x**) is the *joint* probability of a methylation state **x**. As a consequence, *P_**x**_*(**x**) is the Boltzmann-Gibbs distribution of statistical physics^8^, given by

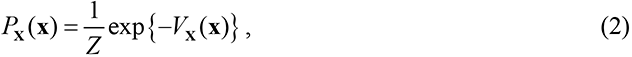

with state energy *V***_x_**(**x**) and partition function

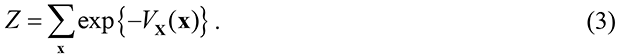

Notably, the potential *V_**x**_*(**x**) - ϕ_0_ quantifies the amount of information associated with the methylation state **x**, which is known to be given by − ln *P*_**x**_(**x**)^9^.

By using the well-known maximum-entropy principle, we determined that the PEL which maximizes our uncertainty about the particular choice of the Boltzmann-Gibbs distribution that is consistent with the methylation means and pairwise correlations is given by

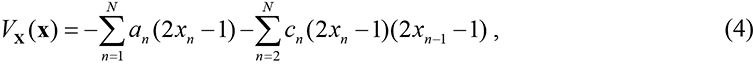

for some parameters *a_n_* and *c_n_*. This leads to a methylation probability *P*_**X**_(**x**) that is modeled by the 1D nearest-neighbor Ising model^8^. Notably, parameter *a_n_* influences the propensity of the *n*-th CpG site to be methylated due to non-cooperative factors, with positive *a_n_* promoting methylation and negative *a_n_* inhibiting methylation, whereas parameter *c_n_* influences the correlation between the methylation states of two consecutive CpG sites *n* and *n* − 1 due to cooperative factors, with positive *c_n_* promoting positive correlation and negative *c_n_* promoting negative correlation (anti-correlation).

We estimated the methylation PEL *V*_**x**_(**X**) from WGBS data corresponding to 35 samples, including stem cells, normal cells from colon, liver, lung, and brain tissues, *matched* cancers from three of these tissues, cultured fibroblasts at 5 passage numbers, CD4^+^ lymphocytes and skin keratinocytes from younger and older individuals, and EBV-immortalized lymphoblasts (see Methods & Supplementary Table 1). To this end, we partitioned the genome into consecutive non-overlapping regions and developed a method for estimating the PEL parameters within each region using a maximum-likelihood approach (see Methods). Our strategy capitalizes on appropriately combining the full information available in multiple methylation reads, especially the correlation between methylation at CpG sites, as opposed to the customary approach of estimating marginal probabilities at each individual CpG site (Fig. 1a).

**Figure 1.**
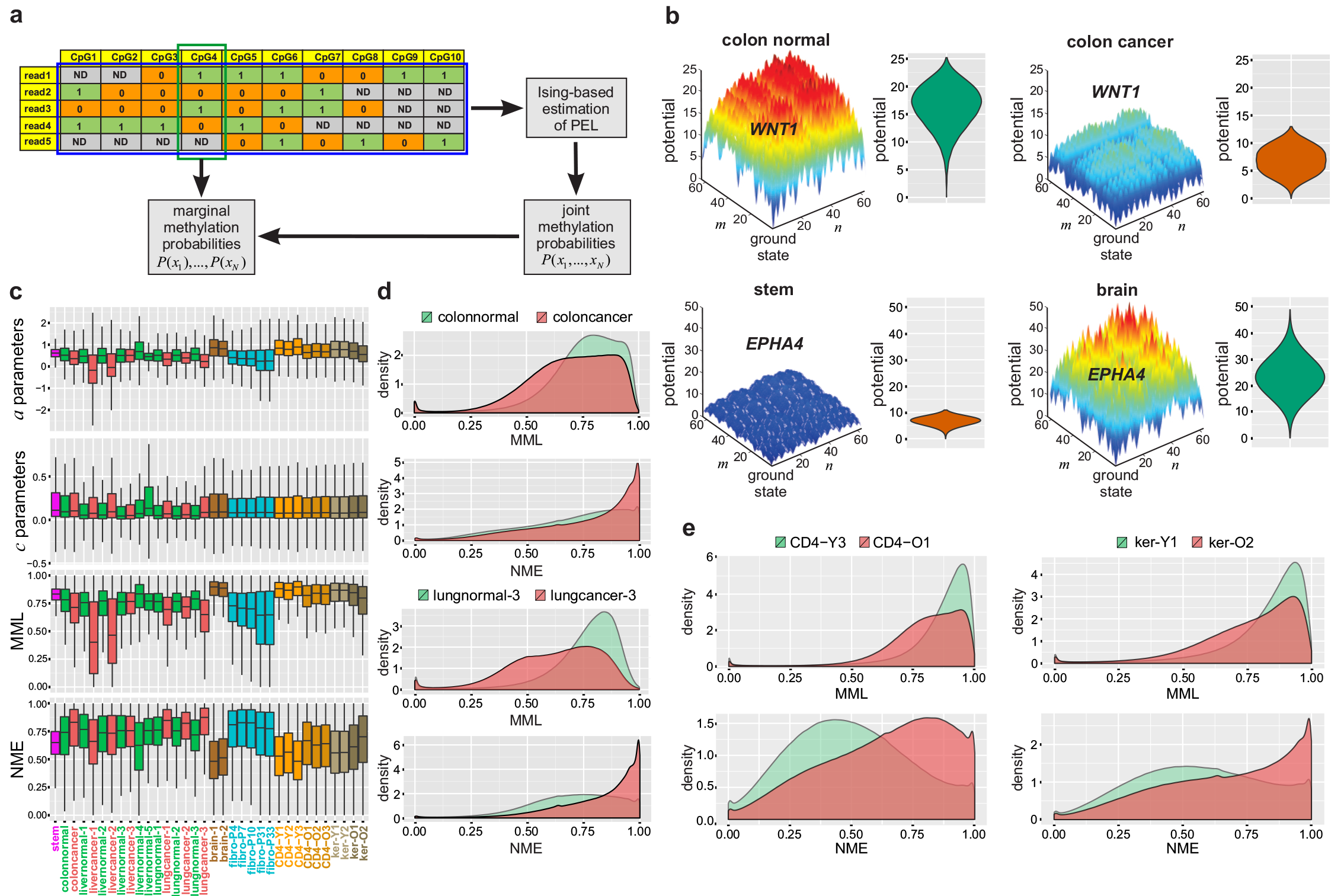
Potential energy landscapes, mean methylation level, and Shannon entropy. **a**,Multiple WGBS reads of the methylation state within a genomic locus are used to form a methylation matrix whose entries represent the methylation status of each CpG site (1: methylated, 0: unmethylated, ND: no data). Most methods for methylation analysis estimate marginal methylation probabilities and means at individual CpG sites by using the methylation information only within each column associated with a CpG site. Our statistical physics approach computes the most likely PEL by determining the likelihood of each row of the methylation matrix, combining this information across rows into an average likelihood, and maximizing this likelihood with respect to the PEL parameters. **b**, PELs associated with the CGIs of *WNT1* in colon normal and colon cancer and *EPHA4* in stem and brain. Point *(m,n)* marks a methylation state, with (0,0) indicating the fully unmethylated state, which turns-out to be the ground state in both examples. **c**, Boxplots of the Ising PEL parameter distributions, as well as the mean methylation level (MML) and normalized methylation entropy (NME) distributions for all samples used in this study. The boxes show the 25% quantile, the median, and the 75% quantile, whereas each whisker has a length of 1.5x the interquartile range. **d**, Genome-wide MML and NME densities associated with two normal/cancer samples show global MML loss in colon and lung cancer, accompanied by a gain in entropy. **e**, Genome-wide MML and NME densities associated with young/old CD4+ lymphocytes and skin keratinocytes show global MML in old individuals, accompanied by a gain in entropy.

For reliable estimation, we reduced the 2*N* − 1 PEL parameters within a genomic region that contains *N* CpG sites to three parameters, *α, β*, and *γ*, characteristic to that region. We did so by setting *a_n_* = *α* + *βρ_n_* and *c_n_* = *γ*/*d_n_*, where *ρ_n_* is the CpG density associated with the *n*-th CpG site and *d_n_* is the CpG distance of the *n*-th CpG site from its “nearest-neighbor” site *n* − 1 (Supplementary Method 1). We used parameter *α* to account for intrinsic factors that uniformly affect CpG methylation and parameter *β* to modulate the influence of CpG density on methylation. Moreover, we set *c_n_* = *γ*/*d_n_* to reflect our expectation that correlation between the methylation states of two consecutive CpG sites decays as the distance between the sites increases.

The resulting PEL encapsulates our view that methylation within a genomic region depends on two distinct factors: the underlying CpG architecture of the genome at that location, quantified by the CpG density *ρ_n_* and distance *d_n_* whose values can be readily determined from the DNA sequence itself, as well as by the current biochemical environment in the nucleus provided by the methylation machinery, quantified by parameters *α, β*, and *γ* whose values must be estimated from available methylation data. Due to its dependence on a small number of parameters, this model allows us to estimate the joint probability distribution of methylation from low coverage WGBS data (as low as 7x in the data used in this study). In turn, this allows one to reliably calculate marginal probabilities at individual CpG sites, compute PELs, and produce a number of novel methylation measures that have not been considered before. We calculated PELs on all 35 samples comprehensively across the entire genome, which can be visualized locally by a 3D representation using Gray’s code (see Methods). For example, computed PELs demonstrate that most methylation states associated with the CpG island (CGI) of *WNT1* in colon normal exhibit high potential (Fig. 1b, fig 3D and violin plots), implying that significant energy is required to leave the fully unmethylated state, which, in this case, is the state of lowest potential (ground state). Any deviation from this state will be rapidly “funneled” back, leading to low uncertainty in methylation. Notably, the methylation states of *WNT1* in colon cancer demonstrate low potential (Fig. 1b, 3D and violin plots), implying that relatively little energy is required to leave the fully unmethylated ground state. In this case, deviations from this state will be frequent and long lasting, leading to uncertainty in methylation. Similarly, the methylation states associated with the CGI of *EPHA4*, a key developmental gene, exhibit low potential in stem cells (Fig. 1b, 3D and violin plots), suggesting that low energy is needed to leave the fully unmethylated state, which is again the ground state, thus leading to uncertainty in methylation. In contrast, *EPHA4* shows high potential in the brain (Fig. 1b, 3D and violin plots), implying that substantial energy is required to leave the fully unmethylated ground state thus leading to low uncertainty in methylation. Lastly, global distributions of the PEL parameters *a* and *c* (Fig. 1c) show that our motivation for using the Ising model is well founded. Specifically, more than 75% of the *c* parameters along the genome are positive, showing extensive cooperativity in methylation (Fig. 1c). Interestingly, a global increase in the values of the *c* parameters is consistently observed in cancer, implying an overall increase in methylation cooperativity in tumors. In addition, most samples demonstrate positive median *a* values, indicating that methylation is more common than nonmethylation, except in two liver cancer samples which were subject to extended extreme hypomethylation. Even in those cases, however, *c* is increased in the tumors.

### Epigenetic entropy quantifies methylation uncertainty in biological states

Due to their first-order marginal nature, means and variances produce a narrow view of methylation and its uncertainty. Previous methods of methylation analysis have attempted to provide a more comprehensive approach by using the notions of epipolymorphism and combinatorial (Boltzmann) entropy^3,4,7^, which are based on empirically estimating the probabilities of specific methylation patterns (epialleles). We have demonstrated that, in contrast to the model-based estimation of joint probabilities and Shannon entropy employed here, empirical estimation of epiallelic probabilities, epipolymorphisms and combinatorial entropies, requires much higher coverage than routinely available from WGBS data (Supplementary Note 1). With regards to a previous study^4^, we often found that the 95% confidence intervals of empirically estimated epipolymorphisms will not include the true values resulting in potentially large errors.

In order to account for variation of the statistical properties of methylation within the estimation regions and perform methylation analysis at a higher resolution, we further partitioned the genome into genomic units (GUs) of 150bp each and characterized methylation within each GU with the probability distribution *P_L_*(*l*) of the methylation level *L* (see Methods). We then quantified methylation uncertainty within a GU containing *N* CpG sites by defining the normalized methylation entropy (NME)

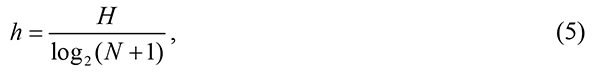

where

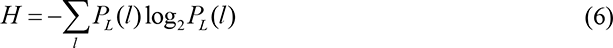

is the informational (Shannon) entropy^9^ of the methylation level within the GU that provides an average assessment of the amount of epigenetic information conveyed by any given GU. When all methylation levels are equally likely (fully disordered state), the NME takes its maximum value of 1 regardless of the number of CpG sites, whereas it achieves its minimum value of 0 only when a single methylation level is observed (perfectly ordered state).

The NME is an effective measure of methylation uncertainty that we can reliably compute genome-wide from low coverage WGBS data using our Ising model, together with the mean methylation level (MML) E[*L*], which is the average of the methylation means at individual CpG sites within a GU. We therefore compared the genome-wide distributions of MML and NME values among samples. Consistent with previous reports, the MML was globally higher in stem cells and brain tissues than in normal colon, liver, and lung and that the same was true for CD4^+^ lymphocytes and skin keratinocytes (Fig. 1c). In addition, the MML was reduced in all seven cancers studied compared to their matched normal tissues, and was also progressively lost in cell culture (Fig. 1c,Fig. 1d). We also observed low NME in stem and brain cells, as well as in CD4^+^ lymphocytes and skin keratinocytes associated with young subjects, and a global increase of NME in most cancers except for liver cancer, which exhibited profound hypomethylation leading to a less entropic methylation state (Fig. 1c, Fig. 1d & Extended Data Fig. 1). While changes of NME in cancer were often associated with changes in MML (Extended Data Fig. 1a), this was often not the case (Extended Data Fig. 1b,c,d), indicating that changes in stochasticity are not necessarily related to changes in mean methylation and demanding that both be assessed when interrogating biological samples. Lastly, we computed MML and NME distributions over selected genomic features and provided a genome-wide breakdown showing lower and more variable methylation levels and entropy values within CGIs and TSSs compared to other genomic features, such as shores, exons, introns, etc. (Extended Data Fig. 2a,b).

**Figure 2.**
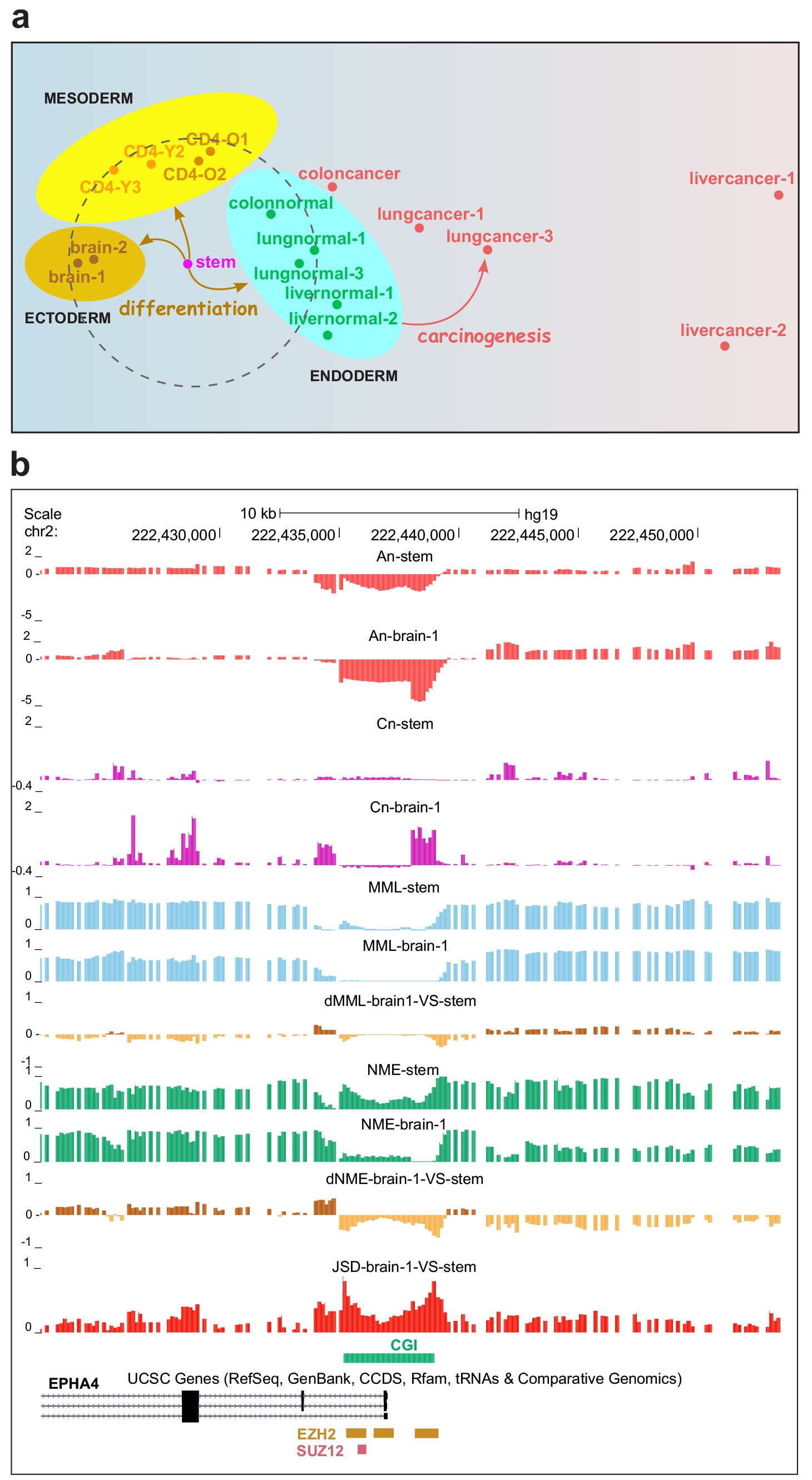
Epigenetic distances delineate lineages and reveal locations of differential PEL regulation. **a** MDS visualization of genomic dissimilarity between 17 diverse cell and tissue samples, evaluated using the Jensen-Shannon distance (JSD), reveals grouping of samples into clear categories based on lineage. **b**, The promoter of *EPHA4* shows binding of EZH2 and SUZ12 and demonstrates negligible differential methylation between stem cells and brain but high JSD that is driven by the PEL parameters, which leads to gain of entropy in brain.

Global hypomethylation and gain in entropy was found in all three CD4CD4^+^ lymphocyte samples from older people compared to three from younger individuals and in both skin keratinocyte samples compared to younger samples (Fig. 1c,e), with the percentage change in entropy being more pronounced. For example, we found an average 23% increase (11 – 38% range) in median NME genome-wide between young and old CD4 samples but only an average 5.6% decrease (3.2 – 8.5% range) in median MML. Note that, for genome-wide comparisons, the 95% confidence intervals for the median are too small to visualize and direct statistical comparisons of the medians are contraindicated. However, to account for biological and statistical variability, we constructed an empirical null distribution amongst the young samples (see Methods) and statistically estimated that up to 34% of the GUs were differentially entropic, demonstrating that profound changes in entropy can result in old individuals. Notably, striking differences were observed between true aging and cell culture. Although passage number in fibroblasts was also associated with progressive global hypomethylation, the entropy distribution was relatively stable (Fig. 1c & Extended Data Fig. 3a). For example, the promoters of *CYP2E1* and *FLNB*, two genes with are known to be downregulated with age^10^, exhibited noticeable gain in methylation level and entropy in old CD4^+^ lymphocytes, which was in stark contrast to the lack of changes with passage in *CYP2E1* and the noticeable loss of entropy in *FLNB* (Extended Data Fig. 3b,c) in cultured fibroblasts. Therefore, age-related PELs in multiple tissues do not seem to be well characterized by increasing fibroblast passage number, and aging appears to be associated with a gain in entropy.

### Informational distances delineate lineages and identify developmentally critical genes

In order to understand the relationship between epigenetic information and phenotypic variation, we sought to precisely quantify epigenetic discordance between pairs of samples using the Jensen-Shannon distance (JSD), which measures the dissimilarity between the probability distributions of the methylation level within a GU across the two samples (see Methods). We then asked if we could use this distance to distinguish colon, lung, and liver from each other and from matched cancers, as well as from stem, brain, and CD4^+^ lymphocytes. For computational feasibility, we limited our study to 17 representative cell and tissue samples and computed all 136 pairwise epigenetic distances genome-wide. We then visualized the results by performing multidimensional scaling (see Methods). The samples fell into clear categories based on developmental germ layers (Fig. 2a), with clusters of ectoderm (brain), mesoderm (CD4), and endoderm (normal colon, lung, and liver) derived tissues located roughly equidistant from stem cells (Fig. 2a, dashed circle). On the other hand, cancerous tissues were far removed from their normal matched tissues as well as from the stem cells (Fig. 2a).

Given the interesting relationship between the stem cell sample and the three germ layers, we examined genes that exhibited noticeable differential methylation level (dMML) and/or JSD in stem cells compared to differential tissues. To this end, we ranked genes based on the magnitude of dMML as well as on the JSD within their promoters (see Methods & Supplementary Data 1) and were surprised to find that many genes known to be involved in development and differentiation showed relatively small changes in dMML yet very high JSD, indicating that the probability distributions of methylation level within their promoters were different, despite little difference in mean methylation level. To explore this further, we asked whether *non-mean* related methylation differences could identify genes between sample groups that would have previously been occult to mean-based analyses by employing a relative JSD-based ranking scheme (RJSD) that assigned a higher score to genes with higher JSD but smaller dMML (see Methods). For example, in the stem cell to brain comparison, we found many key genes at the top of the RJSD list, such as *IGF2BP1, FOXD3, NKX6-2, SALL1, EPHA4*, and *OTX1*, with RJSD-based GO annotation ranking analysis^11^, revealing key categories associated with stem cell maintenance and brain cell development (Supplementary Data 1 & 2). Similarly, 30 GO categories showed 10-fold or greater enrichment in the RJSD list, compared to 5 categories in the MML list (FDR q values ≤ 0.05). We obtained similar results when we compared stem cells to normal lung, with RJSD-based GO annotation analysis revealing key developmental categories and genes in both mesodermal and stem cell categories (Supplementary Data 1 & 2). Comparison of stem cells to CD4^+^ lymphocytes showed enrichment for immune-related functions driven by dMML and many developmental and morphogenesis categories driven by RJSD (Supplementary Data 2).

In contrast, when we compared differentiated tissues, we noticed that dMML-based GO annotation analysis resulted in a higher number of significant categories than RJSD-based analysis, and these were closely related to differentiated functions, such as immune regulation and neuronal signaling in the case of brain and CD4, 16 GO categories with 10-fold or greater enrichment for MML compared to 3 categories for RJSD (Supplementary Data 2; FDR q values ≤ 0.05). Interestingly, when we compared lung normal to cancer, we noticed that RJSD-based GO annotation analysis produced a higher number of significant categories than dMML-based analysis, and these were again related to developmental morphogenesis categories. There were 40 GO categories with 10-fold or greater enrichment for RJSD compared to 7 categories for MML (Supplementary Data 2; FDR q values ≤ 0.05). Taken together, these results show that PEL computation reveals major changes in the probability distributions of DNA methylation associated with developmentally critical genes, and that the shape of these distributions, rather than their means per se, may often be closely related to pluripotency and fate lineage determination in development and cancer.

We next explored the link between changes in the probability state, as reflected by the JSD and the values of the PEL parameters *a_n_* and *c_n_*. For example, a CGI near the promoter of *EPH4A* shows high JSD when comparing stem cells with brain (Fig. 2b). Although this region exhibits comparable mean methylation levels, it displays high JSD over the entire CGI and especially over its shores. Notably, the JSD is not driven by methylation propensity, since the PEL parameters *a_n_* are strongly negative in both stem and brain, in which case the fully unmethylated state is the PEL’s ground state (Fig. 1b, lower panel), resulting in low methylation level within the CGI. However, it is driven by methylation cooperativity at the CGI shores in brain, since the PEL parameters *c_n_* are strongly positive, compared to low methylation cooperativity in stem (almost zero *c_n_*’s) that flattens the PEL (Fig. 1b, lower panel) and results in higher entropy than in brain (Fig. 2b). Intriguingly, the region shows binding of EZH2 and SUZ12, functional enzymatic components of the polycomb repressive complex 2 (PRC2) which regulates heterochromatin formation^12^. Likewise, *SIM2*, a master regulator of neurogenesis, is associated with high JSD regions with similar EZH2/SUZ12 binding, which span several CGIs located near its promoter (Extended Data Fig. 4a). In this case, a gain of entropy is observed in brain, corresponding to a simultaneous loss in methylation propensity (through reduced *a_n_*’s) and a gain in methylation cooperativity (through increased *c_n_*’s). Similar remarks hold for other developmental genes, such as *ASCL2, SALL1*, and *FOXD3* (Extended Data Fig. 4b,c,d; see figure legend for details).

We repeatedly observed the presence of EZH2 and SUZ12 binding sites in areas of high JSD, suggesting that they may play a critical role in generating increased entropy with minimal change in mean methylation. In order to determine whether this association was significant, we used Fisher’s exact test and compared promoters and enhancers with high dMML to those with low dMML as well as promoters and enhancers with high JSD to those with low JSD. We observed several-fold greater enrichments for both EZH2 and SUZ12 binding sites at promoters and enhancers with high JSD vs. low JSD, which provided further evidence of the importance of the JSD (Supplementary Table 2). We then performed binomial logistic regression of EZH2/SUZ12 binding data on JSD scores at promoters and enhancers and found significant positive association (EZH2: score = 5.6 for promoters & 18.1 for enhancers, P value < 2.2 × 10^−16^; SUZ12: score = 6.2 for promoters & 23 for enhancers, P value < 2.2 × 10^−16^; see Supplementary Table 2). Taken together, these results show a significant association of EZH2 and SUZ12 with promoters and enhancers at high JSD regions of the genome, suggesting the intriguing possibility that the PRC2 complex controls stochastic variability in DNA methylation at selected genomic loci by regulating the methylation PEL.

### Methylation energy landscapes uncover bistable behavior associated to genomic imprinting

As a direct consequence of known results of statistical physics that relate the magnetization and covariance of the 1D Ising model with its underlying parameters^8^, we postulated that methylation can be subject to (non-critical) phase transition and, in particular, to a bistable behavior that manifests itself as a coexistence of two distinct epigenetic phases: a fully methylated and a fully unmethylated phase (Extended Data Fig. 5a,b & Supplementary Method 2).

To investigate whether bistability in methylation might be associated with important biological functions, we examined its possible enrichment in several genomic features (see Methods). We found (Supplementary Table 3) that bistable GUs are in general enriched in CpG island shores (ORs > 1 in 29/34 phenotypes, P values < 2.2 × 10^−16^) and promoters (ORs > 1 in 26/34 phenotypes, P values < 1.68 × 10^−9^), but depleted in CGIs (ORs < 1 in 26/34 phenotypes, P values < 2.2 × 10^−16^) and gene bodies (ORs < 1 in 29/34 phenotypes, P values ≤ 3.06 × 10^−14^). Moreover, we noticed that bistable GUs were associated with higher NME than the rest of the genome (Fig. 3a; comparing the bistable regions (yellow) to the rest of the genome (purple)).

**Figure 3.**
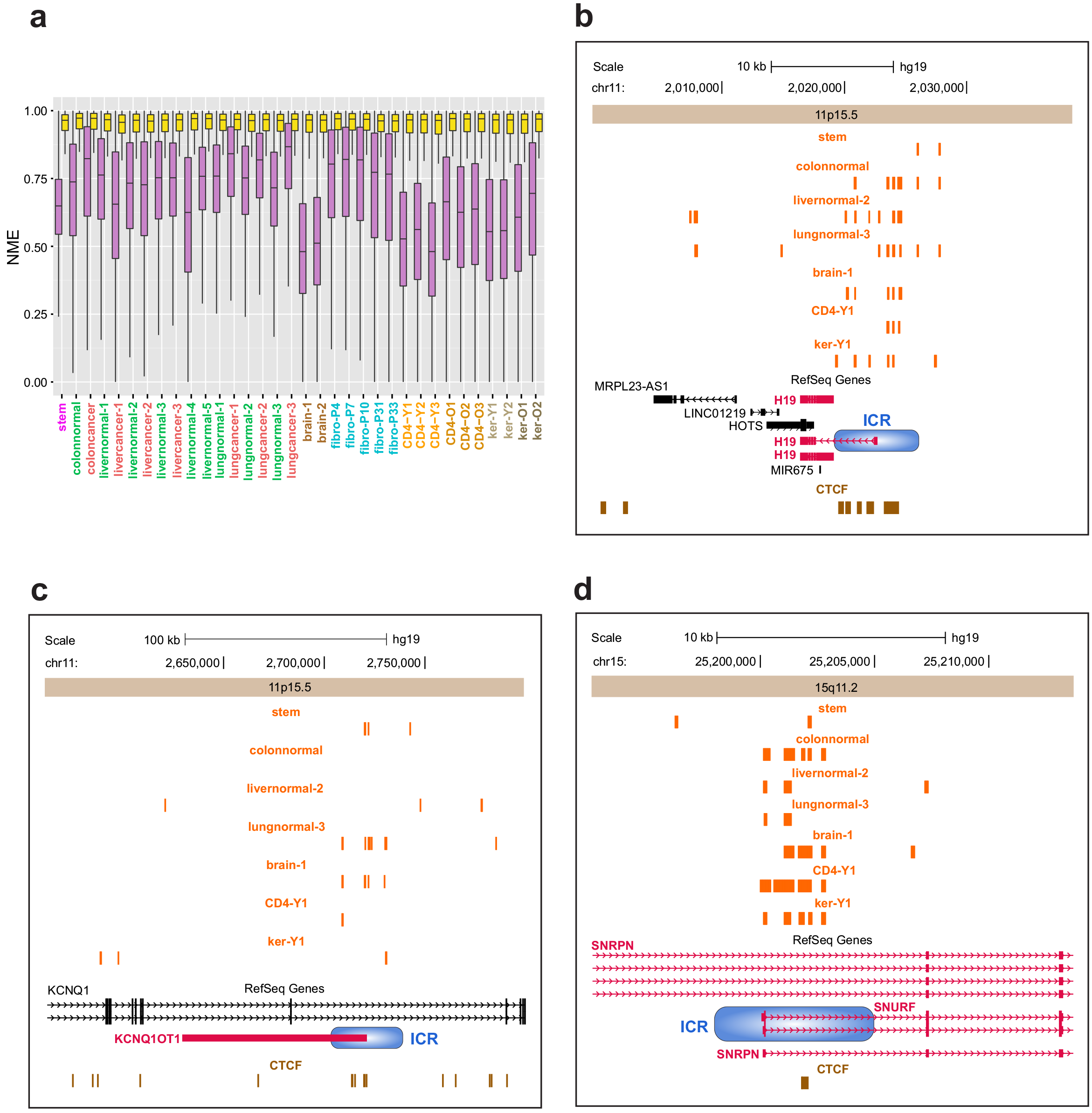
Methylation bistability and imprinting. **a** Boxplots of NME distributions within bistable GUs (yellow) as compared to the rest of the genome (purple). The boxes show the 25% quantile, the median, and the 75% quantile, whereas each whisker has a length of 1.5x the interquartile range. **b,** UCSC genome browser image displaying part of the 11p15.5 chromosomal region associated with *H19*. **c,** A portion of the 11p15.5 chromosomal region associated with *KCNQ1OT1*. **d**, The 15q11.2 chromosomal region near the *SNURF* promoter.

In order to investigate whether methylation bistability is associated with specific genes, we rank-ordered each gene in the genome using a bistability score, which we calculated as the average frequency of methylation bistability within the gene’s promoter in 17 normal samples (see Methods). We found a substantial number of highly ranked genes to be imprinted (Supplementary Data 3). This is attributed to the fact that imprinting has been associated with allele-specific methylation comprising full methylation on one chromosome and complete unmethylation on the other giving rise to bistable methylation^13^. In fact, 82 curated imprinted genes from the *Catalogue of Parent of Origin Effects* (CPOE) were much more highly ranked in our list than would be expected by chance (P value 2.89 × 10^−16^), with notable overrepresentation of imprinted genes near the top of the list. Interestingly, more than 8% of imprinted genes in CPOE appeared in the top 25 bistable genes (*SNRPN, SNURF, MEST, MESTIT1, ZIM2, PEG3, MIMT1*), raising the possibility that imprinting of these genes may be associated with allele-specific methylation of selective loci near their promoters.

We also investigated the possibility that genes subject to monoallelic expression (MAE) are associated with bistability. By using a recently created data set of 4,227 MAE genes, we detected only a slight enrichment of bistability in these genes, likely because MAE is not a result of silenced expression from one of the two alleles^14^. We noticed, however, that 10 MAE genes, not classified in CPOE as being imprinted, exhibited methylation bistability (score > 0.1), raising the possibility that these genes might be imprinted, and one of these, *C11ORF21*, lies within the BWS domain but is not known to be imprinted. Additionally, some of the genes highly ranked in the bistability list that are not imprinted/MAE may be methylated in some cells and not in others.

Considerable effort has been previously expended to identify imprinted genes in the 11p15.5 chromosomal region related to Beckwith-Wiedemann syndrome and loss of imprinting in cancer^15–18^. We therefore assessed the position of bistable marks in this well-studied imprinted locus and revealed a correspondence with known imprinting control regions (ICRs) and CTCF binding sites just upstream of *H19*, as well as near the promoter of *KCNQ1OT1* (Fig. 3b,c). Bistable marks were also found near the *SNURF/SNRPN* promoter, which matched the location of a known ICR (Fig. 3d), as well as near the *PEG3/ZIM2* and*MEST/MESTIT1* promoter regions (Extended Data Fig. 6).

### Entropy blocks predict TAD boundaries

Topologically associating domains (TADs) are structural features of the genome that are highly conserved across tissue types and species.^19–21^. Their importance stems from the fact that loci within these domains tend to frequently interact with each other, with much less frequent interactions being observed between loci within adjacent domains. Genome-wide detection of TAD boundaries is an essential but experimentally challenging task. However, a recent method has demonstrated that TAD boundaries can be reasonably predicted from histone mark ChIP-seq data (CTCF, H3k4me1) using a computational approach^22^. We therefore examined the possibility of using the NME to computationally locate TAD boundaries using WGBS data.

We observed that, in many samples, known TAD boundary annotations were visually proximal to boundaries of entropy blocks (EBs), i.e., genomic blocks of consistently low or high NME values (Fig. 4a & Extended Data Fig. 7, see Methods), which suggests that TAD boundaries may be located within genomic regions that separate successive EBs. To determine whether this is true, we computed EBs in the WGBS stem data and identified 404 regions predictive of TAD boundaries (see Methods). We then found that 5,862 annotated TAD boundaries in H1 stem cells^20^ were located within these predictive regions or were close in a statistically significant manner and correctly identified 6% of the annotated TAD boundaries (362 out of 5,862) derived from 90% of computed predictive regions (see Methods & Supplementary Note 2 for details and P values). We then extended our analysis by combining the TAD boundary annotations for H1 stem cells with available annotations for IMR90 lung fibroblasts^20^ (a total of 10,276 annotations). Since TADs are largely thought to be cell-type invariant^20,21^, we realized that we can predict the location of more TAD boundaries by combining information from EBs derived from additional phenotypes (Fig. 4b). We therefore employed WGBS data from 17 different cell types, computed the corresponding EBs, determined predictive regions for each cell type, and appropriately combined these regions to form a single list that encompasses information (6,632 predictive regions) from all cell types (see Methods). Our analysis produced results similar to those obtained in the case of stem cells and demonstrated that TAD boundaries that fell within identified predictive regions did so significantly more often than expected by chance, resulting in 62% correct identification of the annotated TAD boundaries (6,408 out of 10,276) derived from 97% of computed predictive regions (see Methods & Supplementary Note 2 for details and P values), a performance that can be further improved by including additional phenotypes in our analysis.

**Figure 4.**
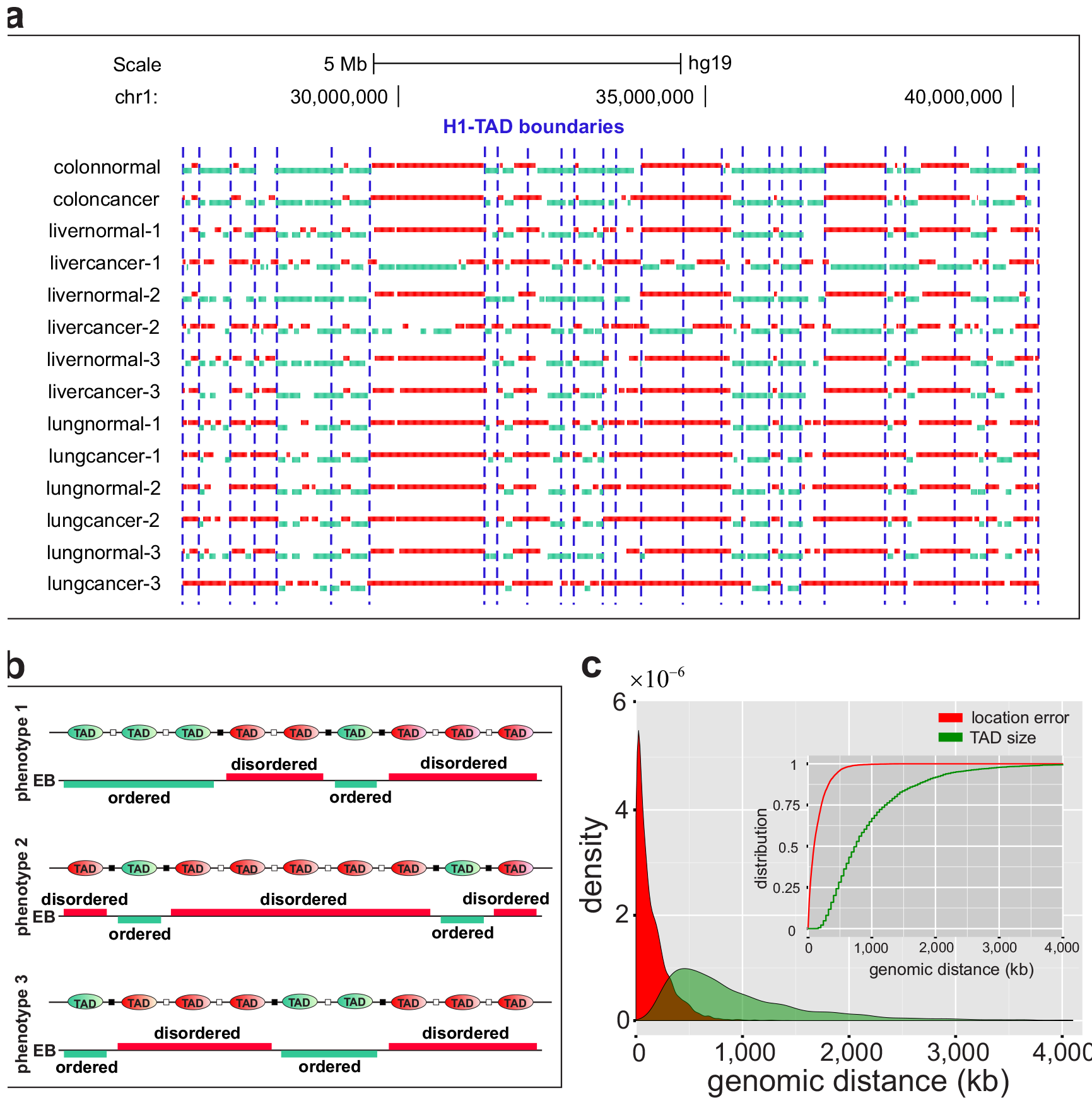
Entropy blocks and TAD boundaries. **a** In the normal/cancer panel, a subset of known TAD boundary annotations in H1 stem cells appeared to be correlated with boundaries of entropic blocks (green: ordered, red: disordered), suggesting that TADs may maintain a consistent level of methylation entropy within themselves. **b**, Regions of entopic transitions can be effectively used to identify the location of some TAD boundaries (black squares). Since TADs are cell-type invariant, the location of more TAD boundaries can be identified using additional WGBS data corresponding to distinct phenotypes. **c**, Probability densities and cumulative probability distributions (insert) of the TAD boundary location error and TAD sizes.

To further assess our predictions, we noted that a TAD boundary can be naturally located at the center of the associated predictive region in the absence of prior information. We then found that errors of locating TAD boundaries in this manner were small when compared to the TAD sizes, as demonstrated by estimating the probability density and the corresponding cumulative probability distribution of the location errors as well as of the TAD sizes using a kernel density estimator (Fig. 4c). Computed cumulative probability distributions implied that the probability that the location error is smaller than *N* bp’s was larger than the probability that the TAD size is smaller than *N*, for every *N*. We therefore concluded that the location error was smaller than the TAD size in a well-defined statistical sense (stochastic ordering). We also observed that the median location error was an order of magnitude smaller than the median TAD size (94-kb vs. 760-kb). Taken together, these observations provide strong statistical evidence that there is an underlying relationship between EBs and TADs, and that this relationship can be easily harnessed to effectively predict TAD boundaries from WGBS data.

### Methylation channels explain epigenetic memory maintenance

Stable conservation of the DNA methylation state is essential for epigenetic memory maintenance. In order to quantify this process, we employed a noisy binary communication channel^9^, which dynamically updates the methylation state and leads to an information-theoretic perspective that enables a fundamental understanding of the relationship between reliability of methylation maintenance, energy availability, and methylation uncertainty. We reasoned that stable maintenance of the methylation state at a CpG site can be approximately modeled by a stochastic methylation channel (MC) near equilibrium, quantified by the transmission probabilities μ and ν of demethylation and *de novo* methylation, respectively (Fig. 5a & Supplementary Method 3). These probabilities are thought to be regulated by the maintenance and *de novo* methyltransferases (DNMT1, DNMT3A, and DNMT3B), by active (TET) and passive demethylation processes, as well as by other potential mechanisms, which are anticipated to be constrained by the free energy available for methylation maintenance.

**Figure 5.**
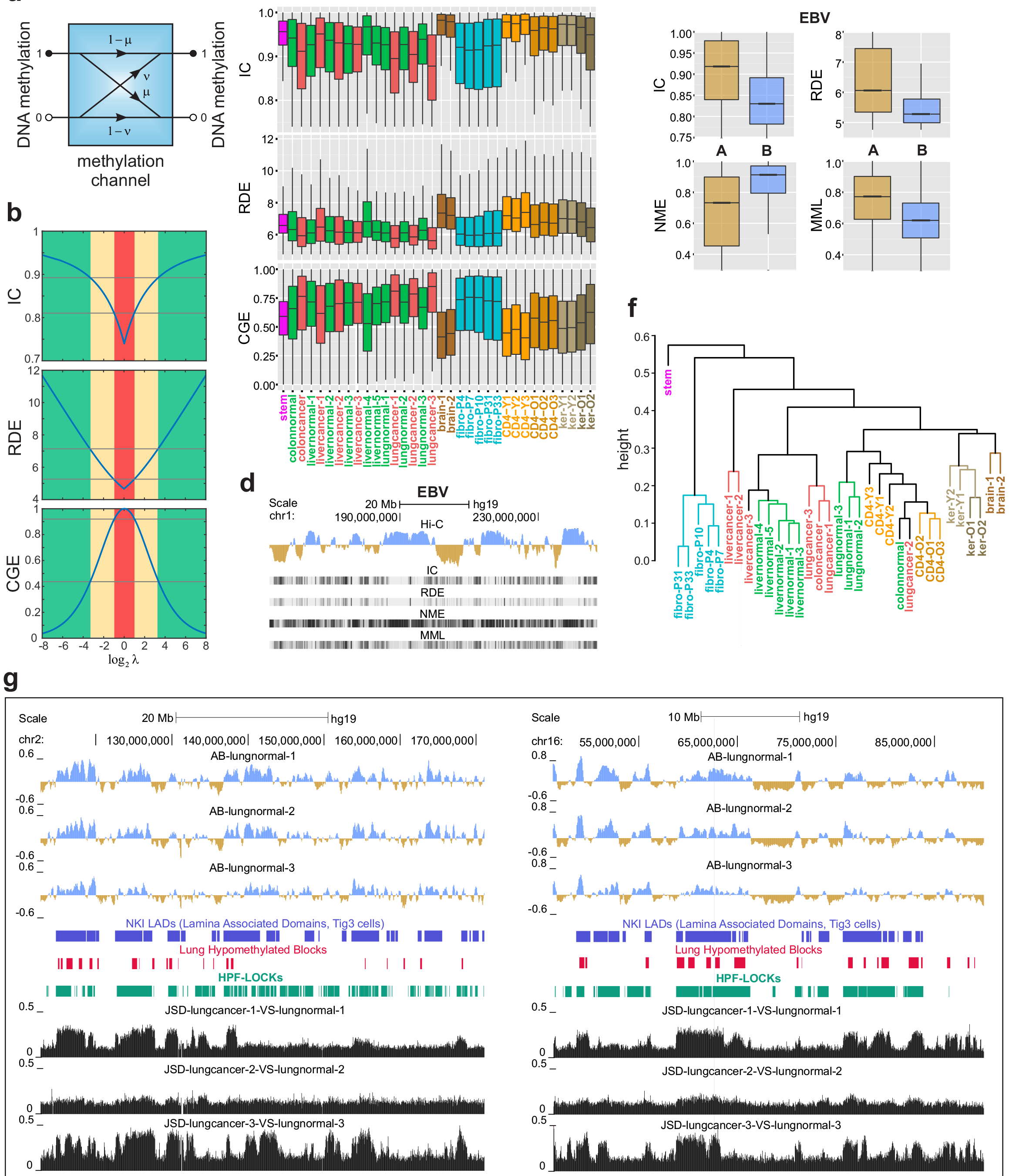
Information theoretic properties of methylation channels predict large scale chromatin organization. **a** A methylation channel (MC) transmits the methylation state at a CpG site of the genome (1: methylated; 0: unmethylated) using four conditional probabilities (p: demethylation probability; v: *de novo* methylation probability). **b**, Derived formulas predict that methylation maintenance by a high capacity MC (IC = 0.89) will dissipate significant energy (RDE = 7.125), achieving a low probability of error (= 0.0073) and resulting in an ordered methylation state (CGE = 0.44), whereas methylation maintenance by a low capacity MC (IC = 0.81) will dissipate a smaller amount of energy (RDE = 5.25), reaching a higher probability of error (= 0.026) and resulting in a disordered methylation state (CGE = 0.92). The thresholds correspond to entropy levels of 0.44 and 0.92 used to identify ordered and disordered genomic units and compute entropic blocks (see Methods). **c**, Boxplots of genome-wide ICs, RDEs and CGEs at individual CpG sites show global differences among cell types. The boxes show the 259 quantile, the median, and the 75% quantile, whereas each whisker has a length of 1.5x the interquartile range. **d,** Analysis of Hi-C and WGBS data reveals that maintenance of the methylation state within compartment B (blue) in EBV cells is mainly performed by MCs with low information capacity (IC) that dissipate low amounts of energy (RDE) resulting in a relatively disordered (NME) and less methylated (MML) state than in compartment A (brown). **e**, Notched-boxplots of genome-wide distributions of IC, RDE, NME and MML demonstrate their attractiveness as features for predicting compartments A/B using WGBS data from single samples, where the notches represent 95% confidence intervals (too small to be visible) around the median. **f**, Net percentage of A/B compartment switching was used as a dissimilarity measure in hierarchical agglomerative clustering. At a given height, a cluster is characterized by lower overall compartment switching than an alternative grouping of samples. **g**, UCSC genome browser images of two chromosomal regions show significant overlap of compartment B in normal lung (blue) with hypomethylated blocks, LADs, and LOCKs. Gain in JSD is observed within compartment B (blue) in normal lung during carcinogenesis.

The amount of methylation uncertainty associated with a MC at a particular CpG site is given by the CG entropy (CGE) *S* = − (1 − *p*) log_2_(1 − *p*)−*p* log_2_*p*, where*p* is the probability of methylation at that site. Notably, only a certain amount of methylation information can be transmitted by a MC, with the maximum possible amount given, on the average, by the information capacity^9^ (IC) (see Supplementary Method 3). Moreover, a MC may drift towards imperfect transmission, which reduces reliability of methylation maintenance by increasing the probability of error. We therefore sought to study the relationship between CGE, IC and reliability of transmission in order to better understand how MCs influence epigenetic memory.

We first reasoned that increased reliability in transmission can be achieved only by a MC that consumes an appreciable amount of free energy, which must be dissipated to the surroundings in the form of heat, in agreement with the first law of thermodynamics. We therefore chose to measure reliability of transmission by computing the amount of dissipated heat. Consistently with general engineering principles^23^, we postulated that the (minimum) energy *E* dissipated during maintenance of the methylation state is approximately related to the probability of transmission error *π* by *E* ~ −*k_B_T*ln*π*, where *k_B_* is Boltzmann’s constant and *T* is the absolute temperature. Since the proportionality factor is not known in this relationship, we used the relative dissipated energy (RDE)

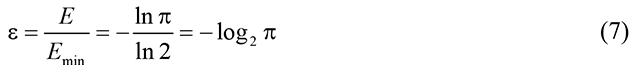

as a measure of reliability in methylation transmission, where *E*_min_ ~ −*k_B_T*ln2 is the least possible energy dissipation (Supplementary Method 3). This implies that higher reliability (lower probability of error) can only be achieved by increasing the amount of free energy available for methylation maintenance, whereas reduction in free energy can lead to lower reliability (higher probability of error). Notably, it is not physically possible for a MC to achieve exact transmission of the methylation state (zero probability of error) since this would require an unlimited amount of available free energy.

We then recognized that calculating ICs and RDEs genome-wide requires computation of the probabilities μ and ν of demethylation and *de novo* methylation at each CpG site of the genome, which is not currently possible. To remedy this, we estimated the turnover ratio *λ* = *ν*/*μ* at each CpG site from WGBS data, derived approximate formulas for the average IC and RDE within a cell population in terms of *λ*, and showed that the CGE can be directly computed from *λ* (Supplementary Method 3). We found that a high capacity MC leads to a less entropic methylation state at the expense of higher energy consumption than a low capacity MC (Fig. 5b, green), which consumes less energy but leads to a highly entropic methylation state (Fig. 5b, red), thus establishing an important relationship between information-theoretic properties of MCs and the nature of epigenetic memory maintenance.

Lastly, we approximately computed ICs, RDEs, and CGEs in individual samples and comparative studies genome-wide (Fig. 5c & Extended Data Fig. 2c,d).We observed a global pattern of IC and RDE loss in colon and lung cancer, accompanied by a global gain in CGE (Fig. 5c), although this was not true in liver cancer (Fig. 5c). Moreover, stem cells demonstrated a narrow range of relatively high IC and RDE values, whereas brain cells, CD4^+^ lymphocytes, and skin keratinocytes exhibited high levels of IC and RDE, with noticeable loss in old individuals (Fig. 5c). Notably, the methylation state within CGIs and TSSs is maintained by MCs whose capacities are overall higher than within shores, shelves, open seas, exons, introns and intergenic regions, and this is accomplished by significantly higher energy consumption (Extended Data Fig. 2c,d). These results reveal an information-theoretic view of genome organization, according to which methylation within certain regions of the genome is reliably transmitted by high capacity MCs leading to low uncertainty in the methylation state at the expense of high energy consumption, while methylation within other regions of the genome is transmitted by low capacity MCs that consume less energy but leading to high uncertainty in the methylation state.

### Information-theoretic prediction of chromatin changes in development and cancer

The 3D spatial organization of the genome allows for regions that are linearly located far from each other to come into proximity and reside in the same regulatory environment. Recent work seeking to understand this organization has demonstrated the existence of cell-type specific compartments A and B^20,21,24^, which are known to be associated with gene-rich transcriptionally active open chromatin and gene-poor transcriptionally inactive closed chromatin, respectively.

Despite the fact that identifying compartments A/B is becoming an increasingly important aspect of fully characterizing the epigenome of a given sample, the availability of such data is limited by cost, technical difficulties, and the need for sizable amounts of input material with intact nuclei required by conformation capture technologies such as Hi-C^24^ Computational prediction methods using data obtained by more routine experimental methods, such as a large number of replicated Illumina 450k DNA methylation microarrays or measures of DNA accessibility, show promise in addressing this problem^25^. In this regard, we sought to predict compartments A and B from local information-theoretic properties of the methylome in individual WGBS samples.

Comparing known Hi-C data from EBV cells to calculated MCs from WGBS data, we observed enrichment of low IC, high NME, and low RDE within compartment B, and the opposite was globally true for compartment A (Fig. 5d,e). These observations led us to hypothesize that information-theoretic properties of methylation maintenance can be effectively used to predict the locations of compartments A and B. To test this prediction, we employed a random forest regression model to learn the informational structure of A/B compartments from available “ground-truth” data. To build this model, we used a feature vector that included the information capacity (IC) and relative dissipated energy (RDE) of a MC, as well as the normalized methylation entropy (NME) and mean methylation level (MML). Random forest regression was capable of reliably predicting A/B compartments from single WGBS samples (Extended Data Fig. 8a), resulting in cross-validated average correlation of 0.74 and an average agreement of 81% between predicted and true A/B signals when using a calling margin of zero, which increased to 0.82 and 91% when the calling margin was set equal to 0.2 (see Methods for details). These results suggest that a small number of local information-theoretic properties of methylation maintenance can be highly predictive of large scale chromatin organization, such as compartments A and B. Once properly trained, the random forest A/B predictor can be applied robustly on any WGBS sample.

Consistent with the fact that compartments A and B are cell-type specific, and in agreement with results of a previous study that demonstrated extensive A/B compartment reorganization during early stages of development^14^, we observed many differences between predicted compartments A/B (Extended Data Fig. 8b,c,d,e). We also found from methylation data alone that the predicted compartment transitions often corresponded to TAD boundaries identified from Hi-C data by Dixon *et. al*.^14^ (Extended Data Fig. 8b). In order to comprehensively quantify observed differences in compartments A and B, we computed percentages of A to B and B to A switching in all sample pairs (Supplementary Data 4). We observed high levels (≥ 20%) of A to B and B to A switching between stem and most of the remaining samples, > 10% switching between brain and most of the remaining samples, and low levels (< 10%) of switching between most normal colon, liver and lung samples. We also noticed > 10% B to A switching between colon, liver and lung normal and most cancer samples.

We subsequently noticed that the net percentage of A/B compartment switching can be employed as a dissimilarity measure between two samples, and used this measure to cluster our samples (Fig. 5f & Methods). The clusters reflect a notable distinction of A/B switching among samples, with 31/34 samples being clustered in a biologically meaningful manner, despite the fact that the random forest model was trained using limited data, and provide evidence that a substantial portion of the observed levels of A/B switching can be attributed to epigenetic differences between the samples. Notably, stem cell differentiation is associated with high levels of chromatin reorganization (Fig. 5f). In particular, differentiated lineages and cancer are cluster together but they are distinguished from each other, while the brain is clustered closest to stem cells, as it also suggested by biochemical studies^26^. Moreover, young CD4 samples form one cluster, whereas old CD4 samples form another, and the same is true for skin.

Intriguingly, normal lung showed strikingly different chromatin organization from lung cancer, as did colon normal from colon cancer (Fig. 5f), so we attempted to relate these changes to known chromatin or methylation structures. Previous studies have demonstrated the presence of large hypomethylated blocks in cancer that are remarkably consistent across tumor types.^27^ These blocks have been shown to correspond closely to large-scale regions of chromatin organization, such as lamin-associated domains (LADs) and large organized chromatin K9-modifications (LOCKs)^2,28^. Consistent with our observations on the information-theoretic properties of compartment B and of carcinogenesis (Fig. 5c,d,e), we asked whether hypomethylated blocks are associated mainly with compartment B (see Methods). We found (Extended Data Fig. 8f) significant overlap with compartment B in normal lung (OR ≈ 3.3, P value < 2.2 × 10^−16^), and the same was true for LADs (OR ≈ 4, P value < 2.2 × 10^−16^) and LOCKs (OR ≈ 5.3, P value < 2.2 × 10^−16^). Interestingly, compartment B in normal tissue may exhibit regions of large JSD values (Fig. 5g), suggesting that considerable epigenetic changes may occur within this compartment during carcinogenesis. We further supported this observation by computing the genome-wide distributions of JSD values between normal/cancer within compartments A and B in normal (Extended Data Fig. 8f). For example, B to A switching in colon cancer included the *HOXA* and *HOXD* gene clusters, whereas B to A switching in lung cancer included the *HOXD* gene cluster but not *HOXA* (Extended Data Fig. 8g,h). Moreover, it included *SOX9* in colon cancer and the tyrosine kinase *SYK* in both colon and lung cancer (Extended Data Fig. 8i). Fewer regions showed A to B switching in cancer, consistent with the directionality of LAD and LOCKs changes in cancer. Interestingly, this included *MGMT* in colon but not lung, a gene implicated in the repair of alkylation DNA damage that is known to be methylated and silenced in colorectal cancer, and the mismatch repair gene *MSH4* (Extended Data Fig. 8j). Together with our previous observation of significant B to A switching between normal/cancer samples, these results suggest that compartment B demarcates genomic regions in which it is more likely for methylation information to be degraded during carcinogenesis.

### Entropic sensitivity quantifies environmental influences on epigenetic stochasticity

Epigenetic changes, such as altered DNA methylation and post-translational modifications of chromatin, integrate external and internal environmental signals with genetic variation to modulate phenotype. In this regard, we sought to investigate the influence of environmental exposure on methylation stochasticity by following a sensitivity analysis approach, which enabled us to quantify the effect of environmental variability on methylation entropy. To this end, we viewed environmental variability as a process that directly influences the methylation PEL parameters and built a stochastic approach that allowed us to approximately relate the amount σ_*h*_ of NME variation with the amount σ of parameter variation by σ_*h*_ ≈ ησ_*h*_, where η measures the absolute rate of NME change due to this variation (see Supplementary Method 4). This suggests using η to quantify entropic sensitivity to environmental conditions, since larger values of η imply larger variation in methylation entropy. We named η the *entropic sensitivity index* (ESI) and developed a method to estimate its values genome-wide from single WGBS data, which quantifies the influence of environmental fluctuations on epigenetic uncertainty in individual samples and comparative studies (Fig. 6, Extended Data Figs. 2e & 9). For example, in colon normal, entropic sensitivity was observed within the CGI associated with *WNT1*, with part of it exhibiting gain in entropy and loss of sensitivity in colon cancer (Fig. 6a).

**Figure 6.**
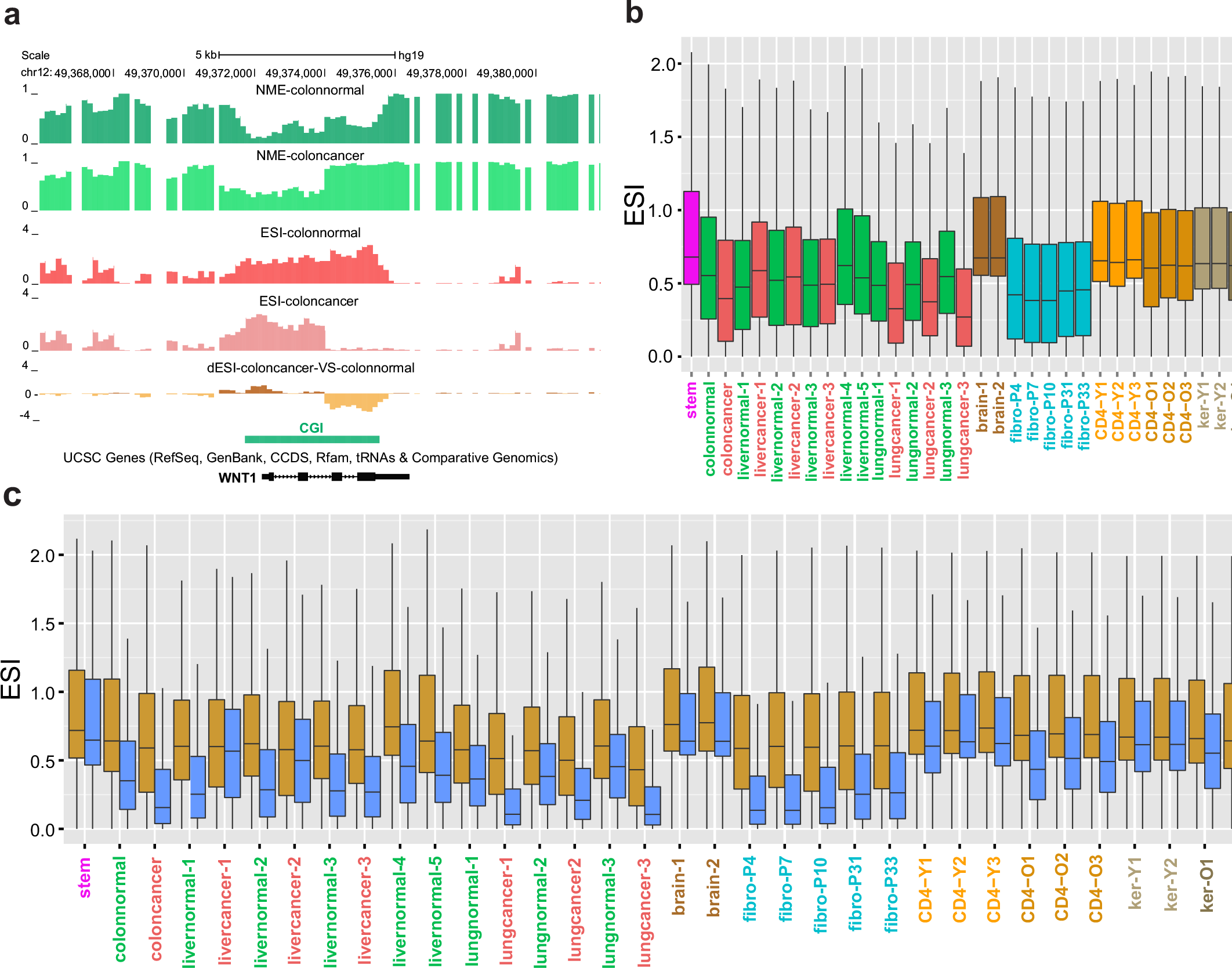
Entropic sensitivity distributions in single samples and comparative studies. **a** Gain of entropy and loss in ESI is observed within a portion of the CGI associated with *WNT1*. **b**, Boxplots of genome-wide ESI distributions corresponding to the samples used in this study reveal global differences in entropic sensitivity across cell types. The boxes show the 25% quantile, the median, and the 75% quantile, whereas each whisker has a length of 1.5x the interquartile range. **c**, Boxplots of genome-wide ESI distributions within compartment A (brown) and compartment B (blue) show that entropic sensitivity is appreciably higher within compartment A than within compartment B.

Globally, we observed differences in ESI among tissues (Fig. 6b & Extended Data Fig. 9a), with stem and brain cells exhibiting higher levels of entropic sensitivity than the rest of the samples. Together with the fact that brain cells are highly methylated (Fig. 1c), high levels of entropic sensitivity would predict that brain can show high rates of demethylation in response to environmental stimuli, consistent with recent data showing that the DNA demethylase *Tet3* acts as a synaptic activity sensor that epigenetically regulates neural plasticity by means of active demethylation^29^, and a similar observation could be true for stem cells and CD4^+^ lymphocytes. Colon and lung cancer exhibited global loss of entropic sensitivity, whereas gain was noted in liver cancer. Moreover, CD4^+^ lymphocytes and skin keratinocytes exhibited global loss of entropic sensitivity in older individuals (Fig. 6b), while cultured fibroblasts showed noticeably lower ESI. Higher and more variable ESI values were observed within CGIs and at TSSs compared to other genomic features, such as shores, exons, and introns (Extended Data Fig. 2e). However, some unmethylated CGIs exhibited low entropic sensitivity (Extended Data Fig. 9b), whereas gain or loss of entropic sensitivity within CGIs was observed between normal and cancer (Extended Data Fig. 9c,d) as well as in older individuals (Extended Data Fig. 9e,f). Notably, differences in ESI were not simply due to entropy itself, as many regions of low entropy showed small ESI values (Extended Data Fig. 9b,c,d), while other such regions exhibited noticeable ESI values (Extended Data Fig. 9c,e,f), indicating sensitivity to environmental perturbations.

We also examined the relationship of entropic sensitivity to higher-order chromatin structure. We found that entropic sensitivity within compartment A was noticeably higher than in compartment B in all samples except stem cells (Fig. 6c), consistent with the notion that transcriptionally active compartment A would be more responsive to stimuli. Moreover, observed differences among normal tissues and between normal and cancer were largely confined to compartment B (Fig. 6c). One could notice substantial loss of entropic sensitivity in compartment B in older CD4^+^ lymphocytes and skin keratinocytes, but not in compartment A. This is in contrast to cell culture that showed a sensitivity gain in compartment B (Fig. 6c).

To further investigate entropic sensitivity changes between tissues, we ranked genes according to their differential ESI (dESI) within their promoters between colon normal and colon cancer (Supplementary Data 5). Colon cancer showed several LIM-domain proteins, including *LIMD2* (ranked 4^th^), which transduce environmental signals regulating cell motility and tumor progression^30^, as well as genes implicated in colon and other types of cancer, such as *QKI* (ranked 1^st^), a critical regulator of colon epithelial differentiation and suppressor of colon cancer^31^ that was recently discovered to be a fusion partner with *MYB* in glioma leading to an autoregulatory feedback loop^32^, *HOXA9* (ranked 8^th^), a canonical rearranged homeobox gene that is dysregulated in cancer, and *FOXQ1* (ranked 9^th^), which is overexpressed and enhances tumorigenicity of colorectal cancer^34^. Together, these results suggest that environmental exposure may influence epigenetic uncertainty in cells with a level of sensitivity that varies along the genome and between compartments in a cell-type specific manner, and present the intriguing possibility that disease, environmental exposure, and aging are associated with substantial loss or gain of entropic sensitivity that could compromise the integration of environmental cues regulating cell growth and function.

## DISCUSSION

In this study, we employed the Ising model of statistical physics to derive, from whole genome bisulfite sequencing, epigenetic potential energy landscapes (PELs) representing intrinsic epigenetic stochasticity. Rather than epigenetic landscapes with external “noise” terms^5,6^, we employed biologically sound principles of methylation processivity, distance-dependent cooperativity, and CpG density to build a rigorous approach to modeling DNA methylation landscapes. This approach was not only capable of modeling stochasticity in DNA methylation from low coverage data, but also allowed genome-wide analysis of Shannon entropy at high resolution and uncovered new properties of the epigenome, such as a relationship between TADs and informational entropy. By incorporating fundamental principles of information theory into a framework of methylation channels, we could also predict in detail high-order chromatin organization from single WGBS samples without performing Hi-C experiments.

Several novel insights ensued from this analysis. We found that Shannon entropy varied markedly among tissues, across the genome and across features of the genome. We consistently observed loss of methylation and entropy gain in cells from older individuals, in contrast to cell culture which exhibited large losses of methylation level and a relatively stable entropy distribution with passage. Genes associated with entropy gain appeared to be highly relevant to aging, although the full implications of this observation require further investigation. In some instances, we observed that high entropy was due to a bistable behavior in methylation level characterized by the coexistence of a fully methylated and a fully unmethylated state. We associated this behavior to many known imprinted regions, in agreement with the fact that imprinting is mostly related to allele-specific methylation.

Rather than identifying differentially methylated regions (DMRs) among compared samples using marginal statistics, we employed the Jensen-Shannon distance (JSD) to compute information-theoretic epigenetic differences genome-wide. This approach allowed us to determine epigenetic differences between individual samples with the potential clinical advantage of identifying specific epigenetic differences which are unique to that sample compared to a matched normal tissue. Analysis of a panel of tissues of diverse origins revealed a “developmental wheel” of the three germ cell lineages around a stem cell hub. Consistently, cancers were extremely divergent and most importantly not intermediate in their methylation properties between stem cells and normal tissue.

We investigated whether the JSD simply embodies mean differences that have been characterized in the past, or if it reveals new insights independent of the mean. To address this question, we identified genomic regions with high JSD but low mean differences between sample pairs, with greater enrichment for many categories of stem cell maintenance or lineage development than found for regions with mean differences per se, suggesting a key role of stochasticity in development. In turn, this type of stochasticity appeared to be driven by localized regions of high cooperativity, which tends to flatten the PEL with little change in mean methylation. We found regions with high JSD and low mean methylation differences to be enriched in Polycomb repressive complex (PRC2) binding sites, suggesting a possible role for PRC2 in stochastic switching during development. Intriguingly, PRC2 components are critical for stochastic epigenetic silencing in an early area of the field of epigenetics, position effect variegation^35,36^, which also involves stochasticity. We suggest that PRC2 is important not only for gene silencing but also for regulating epigenetic stochasticity in general.

A new insight of this work is a relationship between TAD boundaries and entropy blocks. We demonstrated that TAD boundaries can be located within transition domains between high and low entropy in one or more cell types. This suggests a model in which TAD boundaries, which are relatively invariant across cell type and are associated with CTCF binding sites, are potential transition points at which high and low entropy blocks can be demarcated in the genome, and the particular combination of TAD boundaries that transition between high and low entropy define, in large part, the A/B compartments distinguishing tissue types.

We also introduced an information-theoretic approach to epigenetics by means of methylation channels, which allowed us to estimate the information capacity of the methylation machinery to reliably maintain the methylation state. We found a close relationship between information capacity, CG entropy, and relative dissipated energy, as well as between regional localization of high information capacity and attendant high energy consumption (e.g., within shores and compartment A). We realized that informational properties of methylation channels could be used to predict A/B compartments and designed a machine learning algorithm to perform such predictions on widely available WGBS samples from individual tissues and cell culture. This method can be used to predict large scale chromatin organization from DNA methylation data on individual samples. Single paired WGBS data sets of normal and cancer were used to predict A/B compartment transitions. Both colon and lung cancers showed marked compartment switching, most often from B to A, with regions of B to A switching corresponding closely to LADs and LOCKs. Domains of B to A and A to B switching included many genes which are activated or silenced in cancer, suggesting that compartment switching could contribute to cancer.

Lastly, by viewing environmental variability as a process that directly influences the methylation PEL parameters, we introduced the concept of entropic sensitivity, identifying genomic loci where external factors are likely to influence the methylation PEL. While we have only begun to explore the epigenetic implications of entropic sensitivity, it appears that aging and some cancers are associated with global loss of entropic sensitivity and thus to less responsive PELs. If this observation holds true on further study, it could be related to the well-known reduced physiological plasticity of aging, as well as with the autonomous nature of tumor cells.

Although the present study was exploratory and discovery-based in nature, it allowed us to establish a novel modeled-based paradigm for the analysis of epigenetic information that can substantially increase resolution and dramatically reduce the cost of genome-wide epigenetic investigations using small numbers of samples or even individual patient paired-samples, a task that could be crucial in personalized medicine. We however note that biological variability makes broad generalization of results a difficult prospect in human biology. For example, we previously predicted that stochasticity in cancer would increase compared to normal states^5^, but here we observed the unexpected result that certain liver cancer samples have such extreme hypomethylation that they actually experience a reduction in entropy because the methylation state is very likely to be unmethylated. As with all modeling frameworks, the model is only as useful as the data it is built on; contamination, sampling biases and other effects are concerns in this study, just as they are in all other WGBS studies over limited samples. However, the approach presented here should motivate further development of strategies and methods for studying the informational properties of the epigenome and their relationship to disease, and its utility will increase as more WGBS data sets become available for application of this high-resolution methodology.

This study demonstrates a potential relationship between epigenetic information, entropy and energy that may maximize efficiency in information storage in the nucleus. Pluripotent stem cells require a high degree of energy to maintain methylation channels, with certain regions of the genome containing highly deformable PELs corresponding to differentiation branch points, as suggested metaphorically by Waddington, which we can now identify and map the parameters responsible for plasticity. In differentiated cells, large portions of the genome (compartment B, LADs, LOCKs) need not maintain high information capacity and attendant high energy consumption, with their relative sequestration thus providing increased efficiency. However, when domains within compartment B switch to compartment A, previously accumulated epigenetic errors become deleterious and, compounded with reduced entropic sensitivity, may decrease the chance for homeostatic correction.

Finally, the stochastic nature of DNA methylation and the close relationship between methylation entropy, channel capacity, dissipated energy and chromatin structure demonstrated in this paper raises the intriguing possibility that DNA methylation in a given tissue may carry information about both the current state and the possibility of stochastic switching. This information could then be propagated in part through methylation channels over many cycles of DNA replication, even for higher order chromatin organization where the chromatin post-translational modifications themselves may be lost during cell division. This could imply that epigenetic information is carried by a population of cells as a whole, and that this information not only helps to maintain a differentiated state but to also help mediate developmental plasticity throughout the life of an organism.

## METHODS

**Samples for whole genome bisulfite sequencing.** We used previously published WGBS data corresponding to 10 samples, which included H1 human embryonic stem cells^37^, normal and matched cancer cells from colon normal and cancer cells from liver^38^, keratinocytes from skin biopsies of sun protected sites from younger and older individuals^39^, and EBV-immortalized lymphoblasts^40^. We also generated WGBS data corresponding to 25 samples that included normal and matched cancer cells from liver and lung, pre-frontal cortex, cultured HNF fibroblasts at 5 passage numbers, and sorted CD4^+^ T-cells from younger and older individuals, all with IRB approval. Pre-frontal cortex samples were obtained from the University of Maryland Brain and Tissue Bank, which is a Brain and Tissue Repository of the NIH NeuroBioBank. Peripheral blood mononuclear cells (PBMCs) were isolated from peripheral blood collected from healthy subjects and separated by using a Ficoll density gradient separation method (Sigma-Aldrich). CD4+ T-cells were subsequently isolated from PBMCs by positive selection with MACS magnetic bead technology (Miltenyi). Post-separation flow cytometry assessed the purity of CD4^+^ T-cells to be at 97%. Primary neonatal dermal fibroblasts were acquired from Lonza and cultured in Gibco’s DMEM supplemented with 15% FBS (Gemini BioProducts).

**DNA isolation.** Genomic DNA was extracted from samples using the Masterpure DNA Purification Kit (Epicentre). High molecular weight of the extracted DNA was verified by running a 1% agarose gel and by assessing the 260/280 and 260/230 ratios of samples on Nanodrop. Concentration was quantified using Qubit 2.0 Fluorometer (Invitrogen).

**Generation of WGBS libraries.** For every sample, 1% unmethylated Lambda DNA (Promega, cat # D1521) was spiked-in to monitor bisulfite conversion efficiency. Genomic DNA was fragmented to an average size of 350bp using a Covaris S2 sonicator (Woburn, MA). Bisulfite sequencing libraries were constructed using the Illumina TruSeq DNA Library Preparation kit protocol (primers included) or NEBNext Ultra (NEBNext Multiplex Oligos for Illumina module, New England BioLabs, cat # E7535L) according to the manufacturer’s instructions. Both protocols use a Kapa HiFi Uracil+ PCR system (Kapa Biosystems, cat # KK2801).

For Illumina TruSeq DNA libraries, gel-based size selection was performed to enrich for fragments in the 300-400bp range. For NEBNext libraries, size selection was performed using modified AMPure XP bead ratios of 0.4x and 0.2x, aiming also for an insert size of 300-400bp. After size-selection, the samples were bisulfite converted and purified using the EZ DNA Methylation Gold Kit (Zymo Research, cat # D5005). PCR-enriched products were cleaned up using 0.9X AMPure XP beads (Beckman Coulter, cat # A63881).

Final libraries were run on the 2100 Bioanalyzer (Agilent, Santa Clare, CA, USA) using the High-Sensitivity DNA assay for quality control purposes. Libraries were then quantified by qPCR using the Library Quantification Kit for Illumina sequencing platforms (cat # KK4824, KAPA Biosystems, Boston, USA), using 7900HT Real Time PCR System (Applied Biosystems) and sequenced on the Illumina HiSeq2000 (2x100bp read length, v3 chemistry according to the manufacturer’s protocol with 10x PhiX spike-in) and HiSeq2500 (2x125bp read length, v4 chemistry according to the manufacturer’s protocol with 10x PhiX spike-in).

**Quality control and alignment.** FASTQ files were processed using Trim Galore! v0.3.6 (Babraham Institute) to perform single-pass adapter-and quality-trimming of reads, as well as running FastQC v0.11.2 for general quality check of sequencing data. Reads were then aligned to the hg19/GRCh37 genome using Bismark v0.12.3 and Bowtie2 v2.1.0. Separate mbias plots for read 1 and read 2 were generated by running the Bismark methylation extractor using the “mbias_only” flag. These plots were used to determine how many bases to remove from the 5′ end of reads. The number was generally higher for read 2, which is known to have poorer quality. The amount of 5′ trimming ranged from 4bp to 25bp, with most common values being around 10bp. BAM files were subsequently processed with Samtools v0.1.19 for sorting, merging, duplicate removal and indexing.

FASTQ files associated with the EBV sample were processed using the same pipeline described for the in-house samples. BAM files associated with the normal colon and liver samples, obtained from Ziller *et al.*^38^, could not be assessed using the Bismark methylation extractor due to incompatibility of the original alignment tool (MAQ) used on these samples. We therefore followed the advice of the authors and trimmed 4bp from all reads for those files.

**Genomic features and annotations.** Files and tracks bear genomic coordinates for hg19. CGIs were obtained from Wu *et al.*^41^ CGI shores were defined as sequences flanking 2-kb on either side of islands, shelves as sequences flanking 2-kb on either side of shores, and open seas as everything else. The R Bioconductor package “TxDb.Hsapiens.UCSC.hg19.knownGene” was used for defining exons, introns and TSSs. Promoter regions were defined as sequences flanking 2-kb on either side of TSSs. A curated list of enhancers was obtained from the VISTA enhancer browser (http://enhancer.lbl.gov)^42^ by downloading all human (hg19) positive enhancers that show reproducible expression in at least three independent transgenic embryos. Hypomethylated blocks (colon and lung cancer) were obtained from Timp *et al.*^27^. H1 stem cell LOCKs and Human Pulmonary Fibroblast (HPF) LOCKs were obtained from Wen *et al.*^43^ LAD tracks associated with Tig3 cells derived from embryonic lung fibroblasts were obtained from Guelen *et al.*^44^. Gene bodies were obtained from the UCSC genome browser (https://genome.ucsc.edu). H1 and IMR90 TAD boundaries were obtained from http://chromosome.sdsc.edu/mouse/hi-c/download.html. BED files for Hi-C data processed into compartments A and B were provided by Fortin and Hansen (https://github.com/Jfortin1/HiC_AB_Compartments). CTCF and EZH2/SUZ12 binding data were obtained from the UCSC genome browser (Transcription Factor ChIP-seq track (161 factors) from ENCODE).

**Computation and display of potential energy landscapes.** To compute the PEL within a genomic region of interest, we estimated parameters *a_n_* and *c_n_* from WGBS data and used Eq. 4 of the Main Text. Since the size of the methylation state space within a genomic region with *N* CpG sites grows geometrically in terms of *N*, we limited PEL computation within regions of the CGIs near the promoters of *WINT1* and *EPHA4* containing 12 CpG sites. To plot the PEL, we distributed the 2^12^ computed values over a 64 × 64 square grid using a 2D version of Gray’s code^45^, so that methylation states located adjacent to each other in the east/west and north/south directions differ in only one bit.

**Estimation of PEL parameters.** By partitioning the genome into regions of equal size, we estimated the PEL parameters α, β, and γ within a region by maximizing the average log-likelihood 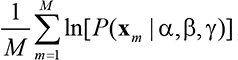, where **x**_1_, **x**_2_,…,**x**_M_ are *M* independent observations of the methylation state within the region. To take into account partially observable methylation states, we replaced *P*(x_m_ | α, β, γ) by the joint probability distribution over only those sites at which methylation information is available, which we calculated by marginalizing *P(x_m_ | α, β, γ)* over these sites. After extensive experimentation, we considered 3-kb estimation regions by striking a balance between estimation and computational performance. To avoid statistical overfitting, we did not model regions with less than 10 CpG sites, as we would had to estimate three parameters from a small number < 10 of variates. We also ignored regions with not enough data for which less than 2/3 of the CpG sites were observed or the average depth of coverage was less than 2.5 observations per CpG site. We finally performed optimization using the multilevel coordinate search (MCS) algorithm^46^.

**Genomic units and methylation level.** Since the Ising model depends on the CpG density and distance, its statistical properties may vary within each 3-kb region used for parameter estimation, suggesting that a smaller genomic region must be employed for high resolution methylation analysis. In this regard, and consistent with the length of DNA within a nucleosome, we partitioned the genome into non-overlapping genomic units (GUs) of 150bp each and performed methylation analysis at a resolution of one GU. We subsequently quantified methylation within a GU that contains *N* CpG sites using the methylation level 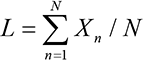 and calculated its probability distribution *P_L_*(*l*) genome-wide directly from the Ising probability distribution of the methylation state.

**Statistical evaluation of differential entropy in aging.** Using the three young CD4 samples, we first computed the absolute NME differences (dNMEs) at each GU associated with all three pairwise comparisons and, by pooling these values, we constructed an empirical null distribution that accounted for biological and statistical variability of differential entropy in the young samples. We then computed the absolute dNME values corresponding to a young-old pair (CD4-Y3,CD4-O1) and performed multiple hypotheses testing to reject the null hypothesis that the observed NME difference is due to biological or statistical variability. By using Bioconductor’s “qvalue” package with default parameters, we performed FDR^47^ and estimated the probability that the null hypothesis is rejected at a randomly chosen GU, thus approximately computing the fraction of GUs which were found to be differentially entropic for reasons other than biological or statistical variability among the young samples.

**Epigenetic distances, multidimensional scaling, and gene ranking.** To quantify methylation differences between two samples within a GU, we employed the Jensen-Shannon distance (JSD)^48^

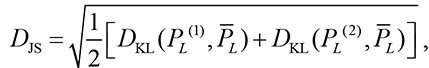

where *P_L_*^(1)^ and *P_L_*^(2)^ are the probability distributions of the methylation level within the GU in the two epigenotypes, 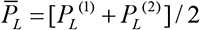 is the average distribution of the methylation level, and

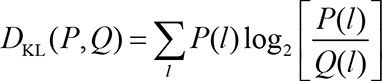

is the relative entropy or Kullback-Leibler divergence^9^. Given a sample taken from one of the two probability distributions *P* and *Q*, the JSD is a normalized distance metric that takes values between 0 and 1, whereas the square JSD is the average information the sample provides about the identity of the distribution: it equals 0 only when the two distributions are identical and reaches its maximum value of 1 if the two distributions do not overlap and can, therefore, be perfectly distinguished from a single sample.

To quantify the epigenetic distance between two samples, we computed the JSDs between all corresponding pairs of GUs genome-wide, sorted these values in increasing order, and determined the smallest value in the list such that 90% of the distances is less than or equal to that value (90-th percentile). To visualize epigenetic similarities or dissimilarities between samples, we computed the epigenetic distances between all pairs of samples, formed the corresponding dissimilarity matrix, and employed a two-dimensional representation using multidimensional scaling (MDS) based on Kruskal’s non-metric method, to find a twodimensional configuration of points whose inter-point distances correspond to the epigenetic dissimilarities among the samples.

To rank genes in terms of the magnitude of dMML, or the JSD, within their promoters, we centered a 4-kb window at the TSS of each gene in the genome, computed the absolute dMML or JSD value within each GU that “touches” this window, and scored the gene by averaging these values. To rank genes using a relative JSD scheme that assigns a higher score to genes with higher JSD but smaller dMML, we scored a gene by the ratio of its ranking in the dMML-ranked list to that in the JSD-ranked list.

**Methylation bistability and imprinting.** To identify bistable GUs in a given WGBS sample, we detected bimodality in the probability distribution *P_L_*(*l*) of the methylation level within a GU. To evaluate enrichment of bistability in a particular genomic feature, we defined two binary (0-1) random variables *R* and *B* for each GU, such that *R* = 1, if the GU overlaps the genomic feature, and *B* = 1, if the GU is bistable. We then tested against the null hypothesis that *R* and *B* are statistically independent by applying the *χ*^2^-test on the 2×2 contingency table for *R* and *B* and calculated the odds ratio (OR) as a measure of enrichment. We evaluated bistability enrichment within CGIs, shores, promoters, and gene bodies. To evaluate possible association between bistability and gene imprinting, we calculated the fraction of bp’s within the promoter region of a gene that overlapped bistable GUs. Consistent with our expectation that genomic imprinting is highly conserved across tissue types, we assigned a bistability score at each gene by averaging the fractions of bistable bp’s calculated from the normal samples. We then used these scores to rank the genes in order of decreasing bistability. To calculate a P value for the CPOE set of 82 imprinted human to be ranked higher in the bistability list than by chance, we computed the P value of each imprinted gene by testing against the null hypothesis that the gene appears at a random location in the bistability list. We then used the gene’s rank as the test statistic and noticed that, under the null hypothesis, its distribution is uniform, which implies that we can calculate the P value by dividing the gene’s ranking in the bistability list by the total number of genes in the list. Since the number of imprinted genes identified in the bistability list (68 genes) is much smaller than the total number of genes (15,820 genes), we assumed statistical independence of the individual hypothesis tests and combined the resulting P values by using Fisher’s method.

**Computation of entropic blocks.** Computation of EBs requires detection of ordered and disordered blocks; i.e., large genomic regions of consistently low or high NME values. To effectively summarize the genome-wide status of NME in a single sample, we computed the NME value *h* within each GU and classified it into one of three classes: ordered (0 ≤ *h* ≤ 0.44), weakly ordered/disordered (0.44 ≤ *h* ≤ 0.92), and disordered (0.92 ≤ *h* ≤ 1). We determined the threshold values by investigating the relationship between the NME within a GU that contains one CpG site and the ratio of the probability *p* of methylation to the probability 1-*p* of unmethylation at that site. To this end, we focused on the odds ratio *r* = *p*/(1 −*p*) and considered the methylation level to be “ordered” if *r* ≥ 10 or *r* ≤ 1/10 (i.e., if the probability of methylation is at least 10x larger than the probability of unmethylation, and likewise for the probability of unmethylation), in which case, *p* ≥ 0.9091 or*p* ≤ 0.0909, which correspond to a maximum NME value of 0.44. Moreover, we considered the methylation level to be “disordered” if 1/2 ≤ *r* ≤ 2 (i.e., if the probability of methylation is no more than 2x the probability of unmethylation, and likewise for the probability of unmethylation), in which case, 0.3333 ≤ *p* ≤ 0.6667, which corresponds to a minimum NME value of 0.92.

To compute EBs, we slid a window of 500 GUs (75-kb) along the genome and labeled the window as being ordered or disordered if at least 75% of its GUs were effectively classified as being ordered or disordered, respectively. We then determined ordered or disordered blocks by taking the union of all ordered or disordered windows and by removing discordant overlappings.

**Prediction of TAD boundaries.** Using EBs computed for a given epigenotype, we identified predictive regions of the genome that might contain TAD boundaries by detecting the space between successive EBs with distinct labels (ordered or disordered). For example, if an ordered EB located at chr1: 1-1000 were followed by a disordered EB at chr1: 1501-2500, then chr1: 1001-1500 was deemed to be a predictive region. To reduce false identification of predictive regions, we did not consider successive EBs of the same type, since the genomic space between two such EBs may be due to missing data or other unpredictable factors. To control the resolution of locating a TAD boundary, we only considered gaps smaller than 50-kb. This resulted in a resolution of an order of magnitude smaller than the mean TAD size (~900-kb). To combine predictive regions obtained from methylation analysis of several distinct epigenotypes, we computed the “predictive coverage” of each bp by counting the number of predictive regions that contained the bp. We then combined predictive regions by grouping consecutive bp’s whose predictive coverage was at least 4. We subsequently applied this method on WGBS data corresponding to 17 distinct cell and tissue types (stem, colonnormal, coloncancer, livernormal-1, livercancer-1, livernormal-2, livercancer-2, livernormal-3, livercancer-3, lungnormal-1, lungcancer-1, lungnormal-2, lungcancer-2, lungnormal-3, lungcancer-3, brain-1, brain-2), and analyzed our results using ‘GenometriCorr’^49^, a statistical package for evaluating the correlation of genome-wide data with given genomic features. Finally, we considered a boundary prediction to be “correct” when the distance of a “true” TAD boundary from the center of a predictive region was less than the first quartile of the “true” TAD width distribution (Fig. 4c insert – green).

**A/B compartment prediction and analysis.** Genome-wide prediction of A/B compartments was performed by a random forest regression model. We trained this model using a small number of available Hi-C data associated with EBV and IMR90 samples^50^, as well as A/B tracks produced by the method of Fortin and Hansen (FH) using long-range correlations computed from pooled 450k array data associated with colon cancer, liver cancer, and lung cancer samples^25^. Due to the paucity of currently available Hi-C data, we included the FH data in order to increase the number of training samples and improve the accuracy of performance evaluation. We first paired the Hi-C and FH data with WGBS EBV, fibro-P10, and colon cancer samples, as well as with samples obtained by pooling WGBS liver cancer (livercancer-1, livercancer-2, livercancer-3) and lung cancer (lungcancer-1, luncancer-2, lungcancer-3) data. We subsequently partitioned the entire genome into 100-kb bins (to match the available Hi-C and FH data), and computed eight information-theoretic features of methylation maintenance within each bin (median values and interquartile ranges of IC, RDE, NME and MML). By using all feature/output pairs, we trained a random forest model using the R package ‘randomForest’ with its default settings, except that we increased the number of trees to 1,000. We then applied the trained random forest model on each WGBS sample and produced A/B tracks that approximately identified A/B compartments associated with the samples. Since regression takes into account only information within a 100-kb bin, we averaged the predicted A/B values using a three-bin smoothing window and removed from the overall A/B signal its genome-wide median value, as suggested by Fortin and Hansen^25^.

To test the accuracy of the resulting predictions, we employed 5-fold cross validation, which involved training using four sample pairs and testing on the remaining pair for all five combinations. We evaluated performance by computing the average correlation as well as the average percentage agreement between the predicted and each of the “ground-truth” A/B signals within 100-kb bins at which the absolute values of the predicted and “ground-truth” signals were both greater than a calling margin, where we used a non-zero calling margin to remove unreliable predictions. We finally calculated agreement by testing whether the predicted and the “ground-truth” A/B values within a 100-kb bin had the same sign.

For each pair of WGBS samples, we computed the percentage of A to B compartment switching by dividing the number of 100-kb bin pairs for which an A prediction is made in the first sample and a B prediction is made in the second sample by the total number of bins for which A/B predictions were available in both samples, and similarly for the case of B to A switching. We summed these percentages and formed a matrix of dissimilarity measures, which we then used as an input to a Ward error sum of squares hierarchical clustering scheme^51^, which we implemented using the R package ‘hclust’ by setting the method variable to ‘ward.D2’.

To test the significance of overlapping of hypomethylated blocks, LADs, and LOCKs with compartment B, we used available hypomethylated blocks, LOCKs, and LADs, and predicted compartment B data for the lungnormal-1, lungnormal-2, and lungnormal-3 samples, which best match the previous tracks. To evaluate enrichment of hypomethylated blocks (and similarly for LADs and LOCKs) within compartment B, we defined two binary (0-1) random variables *R* and *B* for each GU, such that *R* = 1 if the GU overlaps a block, and *B* = 1 if the GU overlaps compartment B. We then tested against the null hypothesis that *R* and *B* are statistically independent by applying the *χ*^2^-test on the 2 × 2 contingency table for *R* and *B* and calculated the odds ratio (OR) as a measure of enrichment.

## Acknowledgements

This work was supported by NIH Grants DP1ES022579, R01AG042187, and R01CA054348 to A.P.F., NSF Grant CCF-1217213 to J.G., and NIH Grant AG021334 to Jeremy Walston. E.P. was supported in part by the Medical Scientist Training Program. The funders had no role in study design, data collection and analysis, decision to publish, or preparation of the manuscript. We thank: Xin Li, Amy Vandiver, and Jeremy Walston for cells and/or FASTQ files; Rakel Trygvadottir, Birna Berndsen, and Adrian Idrizi for sequencing; Jean-Phillippe Fortin and Kasper Hansen for providing A/B compartment data; Andrew Sullivan and Alex Gimelbrant for access to imprinted gene and MAE datasets; and Winston Timp and Kasper Hansen for critical reading of the manuscript.

### Competing Financial Interests

The authors declare no competing financial interests.

## Supplemental Information

**Extended Data Figure 1.**
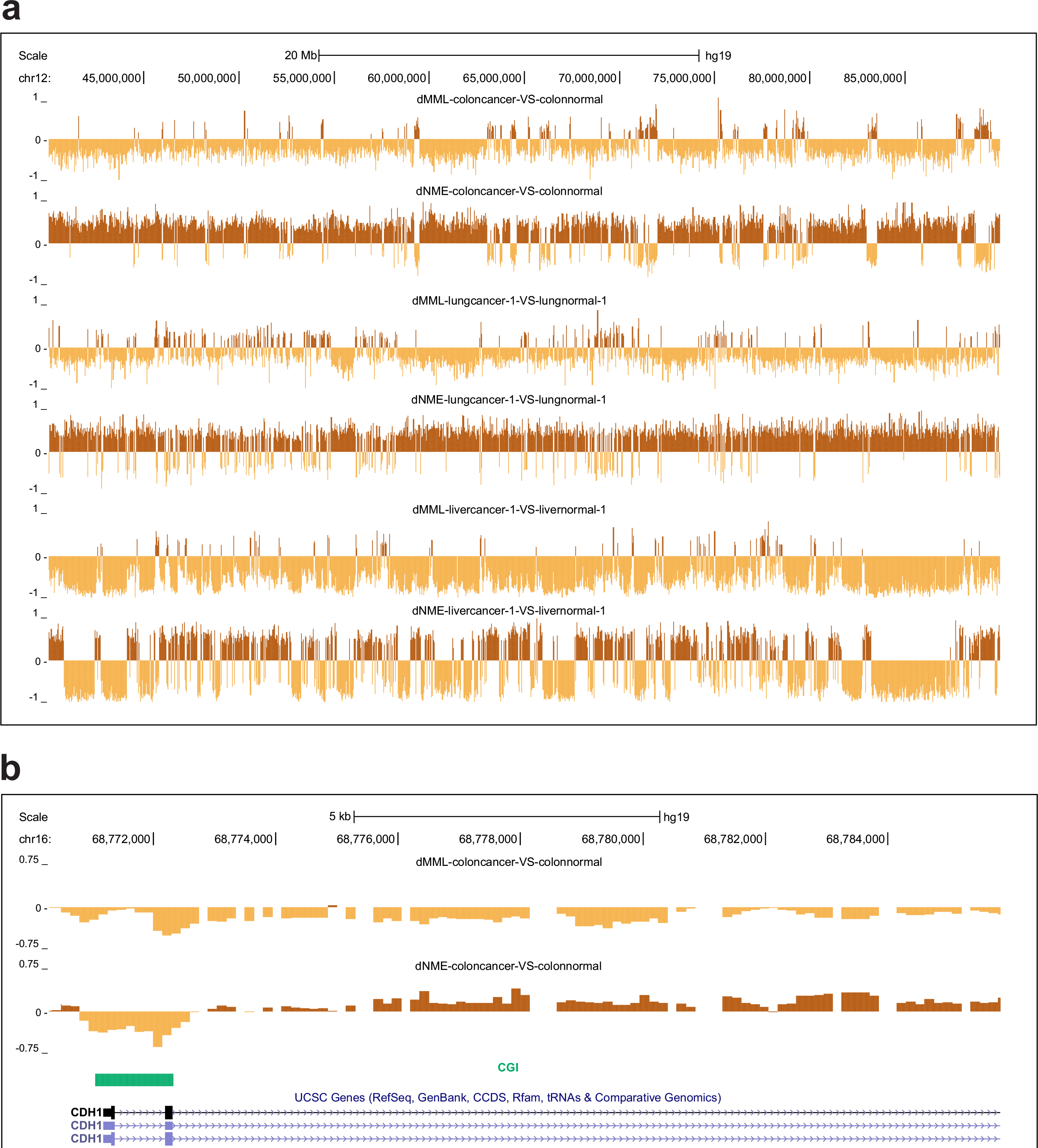
Changes in methylation entropy in cancer. **a**, UCSC genome browser image showing significant loss in mean methylation level (dMML) in colon and lung cancer, accompanied by gain in methylation entropy (dNME). Liver cancer exhibits loss of methylation entropy within large regions of the genome due to profound hypomethylation. **b**, The CGI near the promoter of *CDH1*, a tumor suppressor gene, exhibits entropy loss in colon cancer. **c**, The CGI near the promoter of *NEU1* shows gain of methylation entropy in lung cancer. *NEU1* sialidase is required for normal lung development and function, whereas its expression has been implicated in tumorigenesis and metastatic potential. **d**, Noticeable loss of methylation entropy is observed in liver cancer at the shores of the CGI near the promoter of *ENSA*, a gene that is known to be hypomethylated in liver cancer.

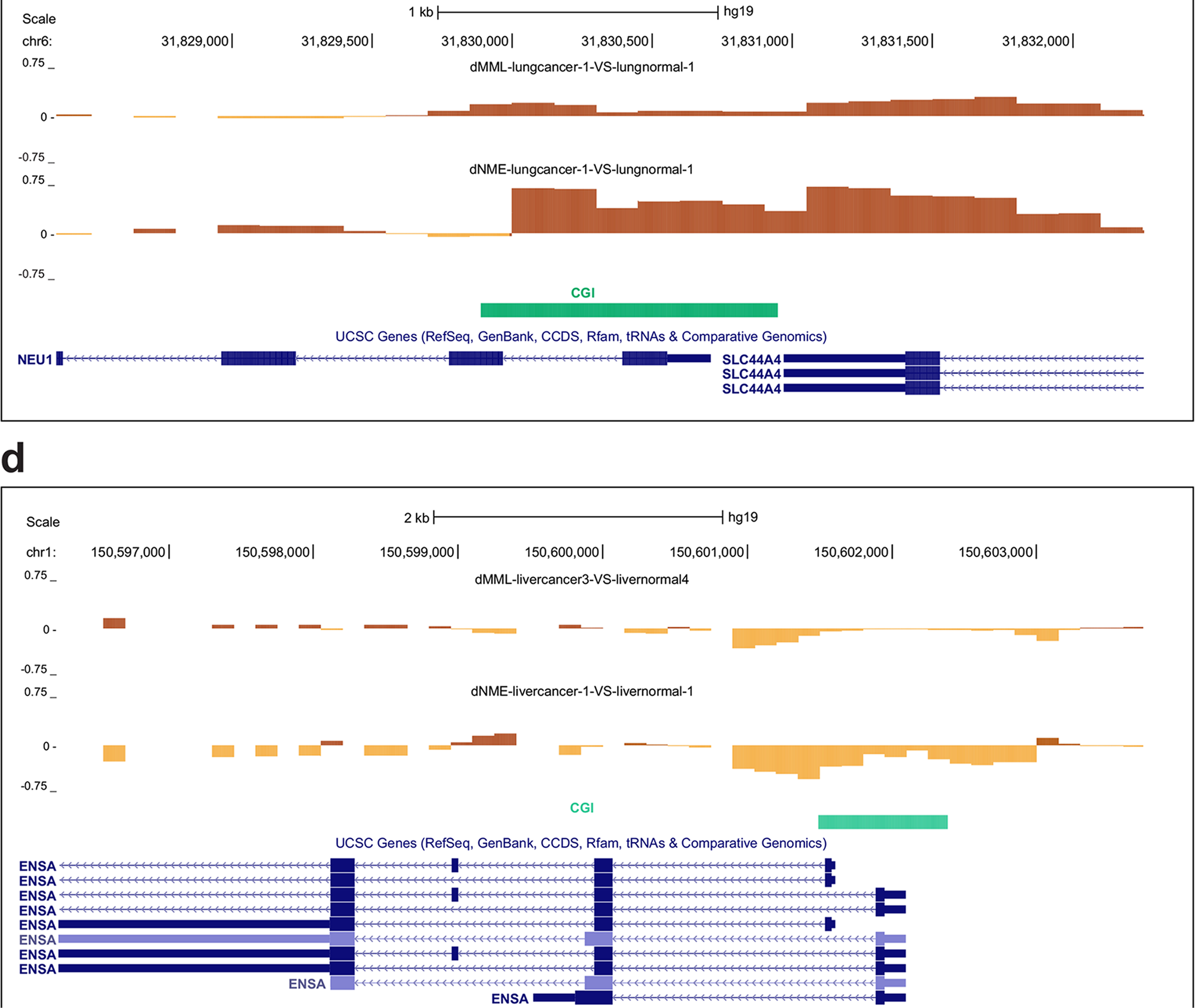

**Extended Data Figure 2.**
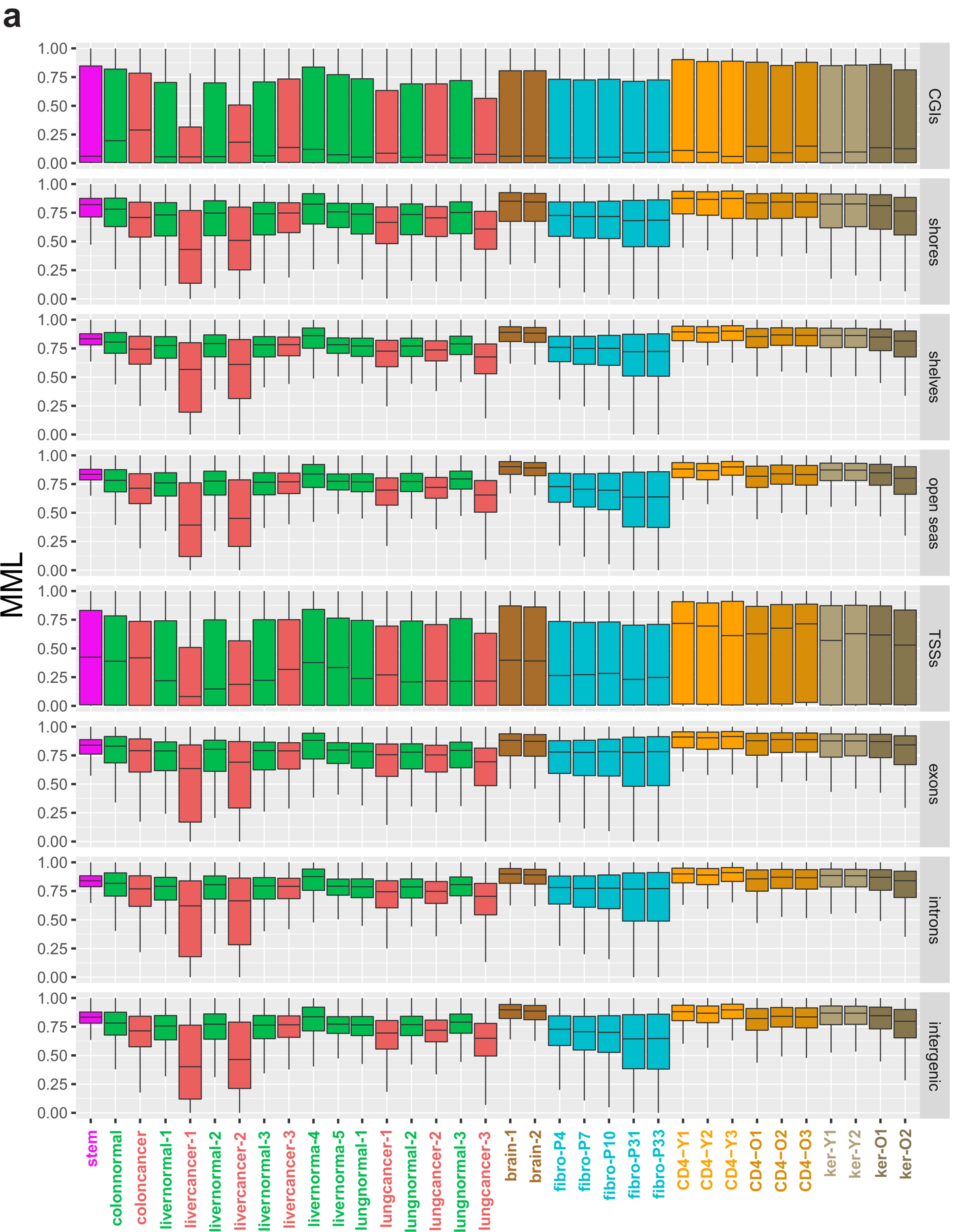
Breakdown of genome-wide distributions of methylation measures. **a**, Boxplots of genome-wide distributions of methylation measures for all samples used in this study within CGIs, shores, shelves, open seas, TSSs, exons, introns, and intergenic regions. The boxes show the 25% quantile, the median, and the 75% quantile, whereas each whisker has a length of 1.5x the interquartile range. **a**, Mean methylation level (MML). **b**, Normalized methylation entropy (NME). **c**, Information capacity (IC). **d**, Relative dissipated energy (RDE). **e,** Entropic sensitivity index (ESI).

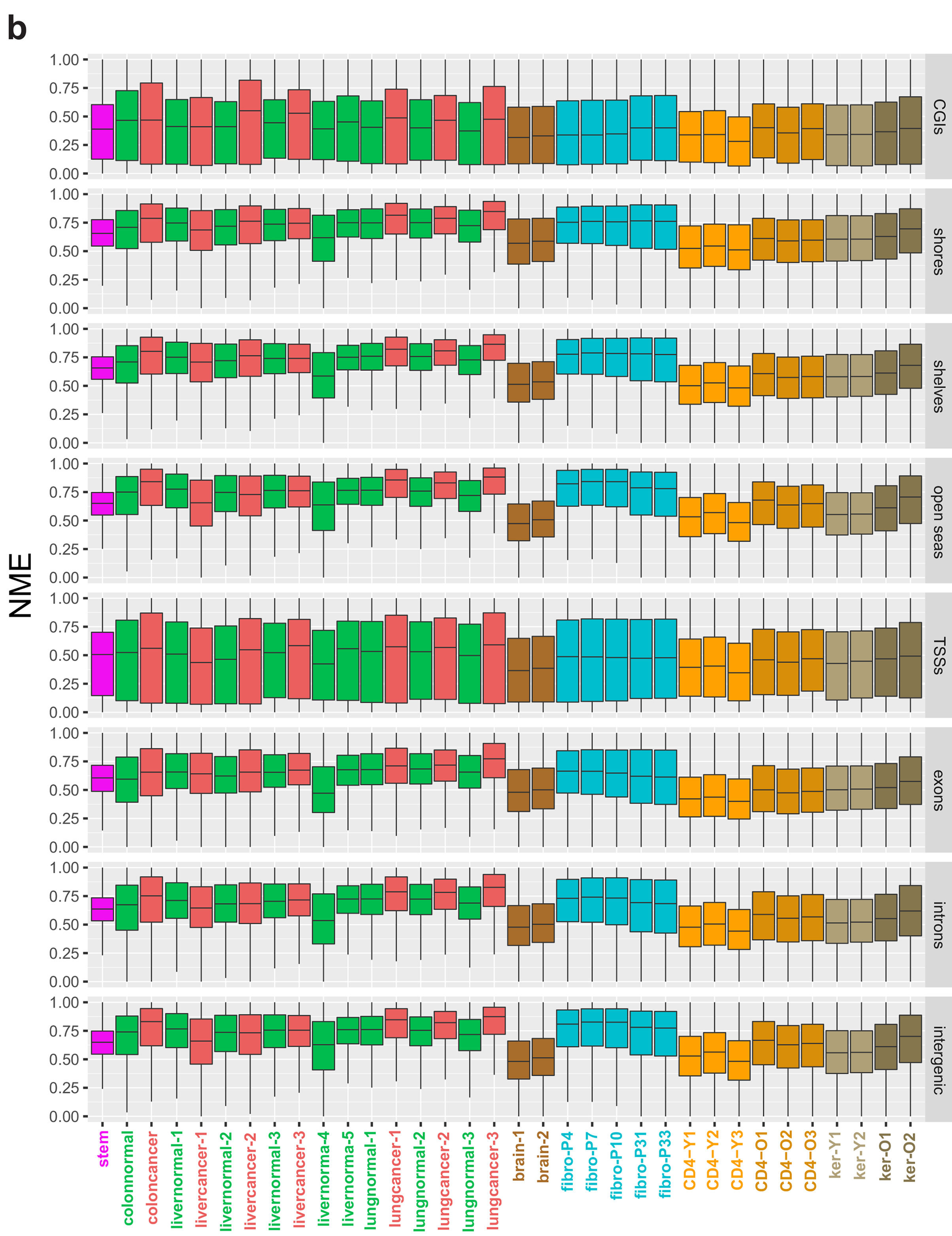

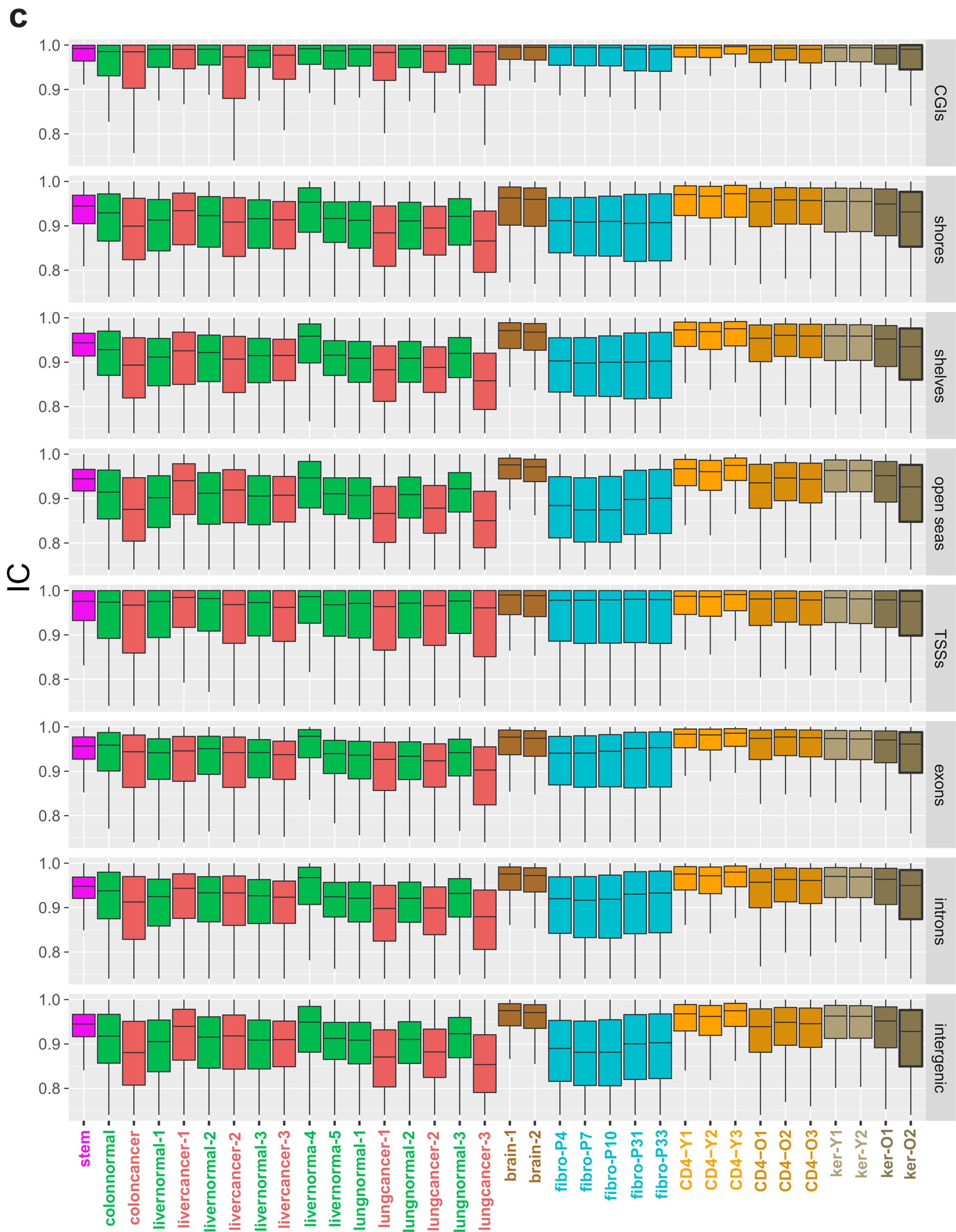

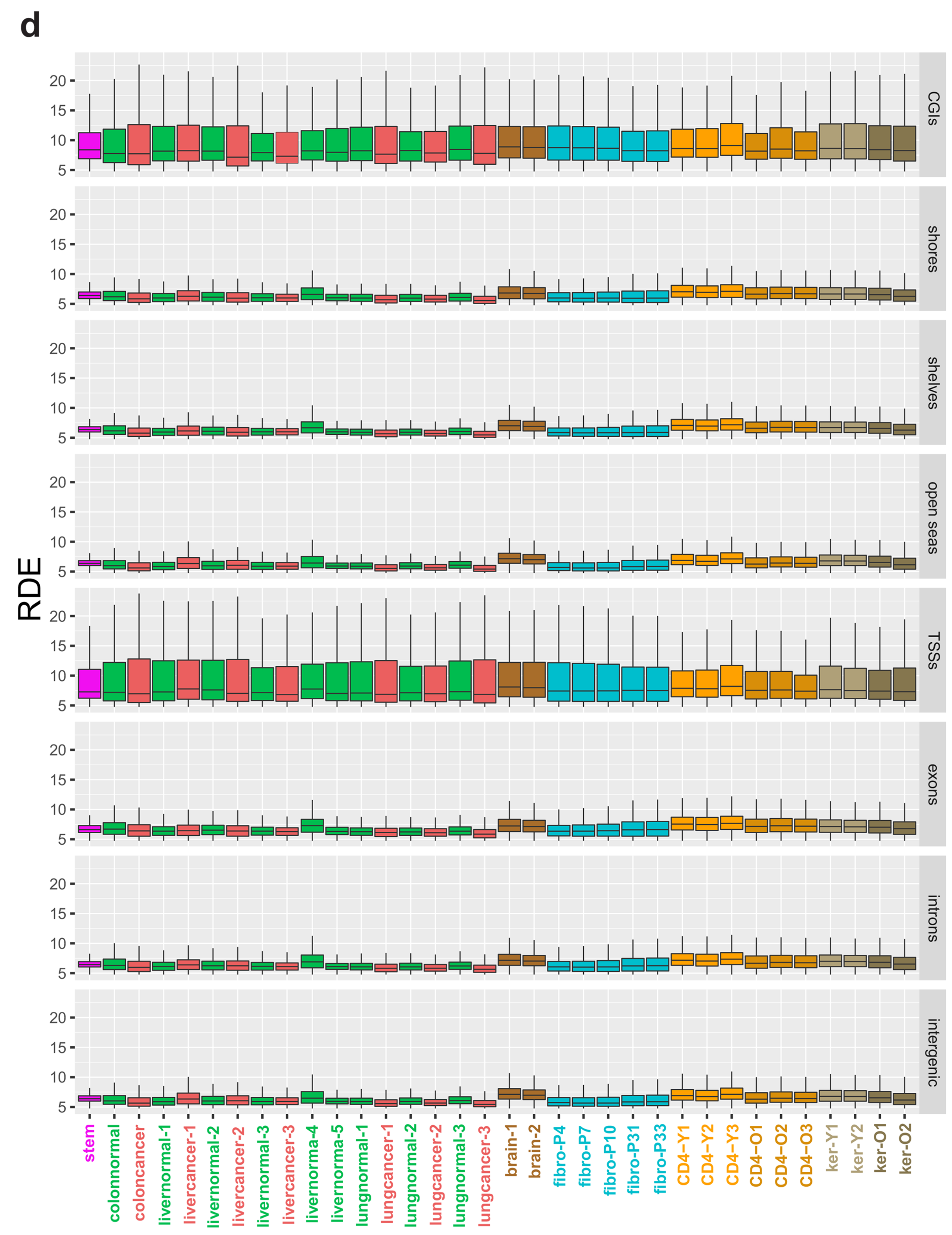

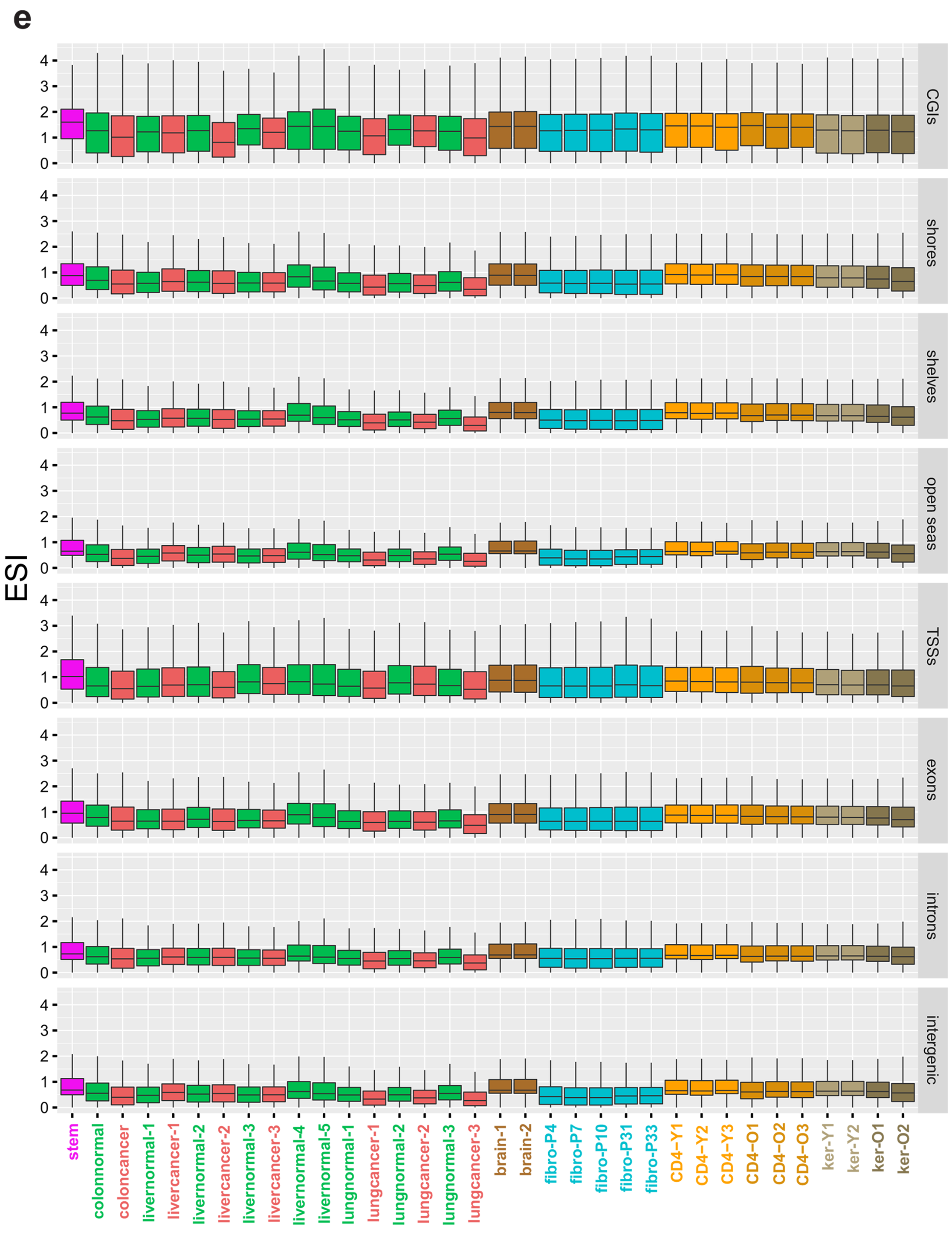

**Extended Data Figure 3.**
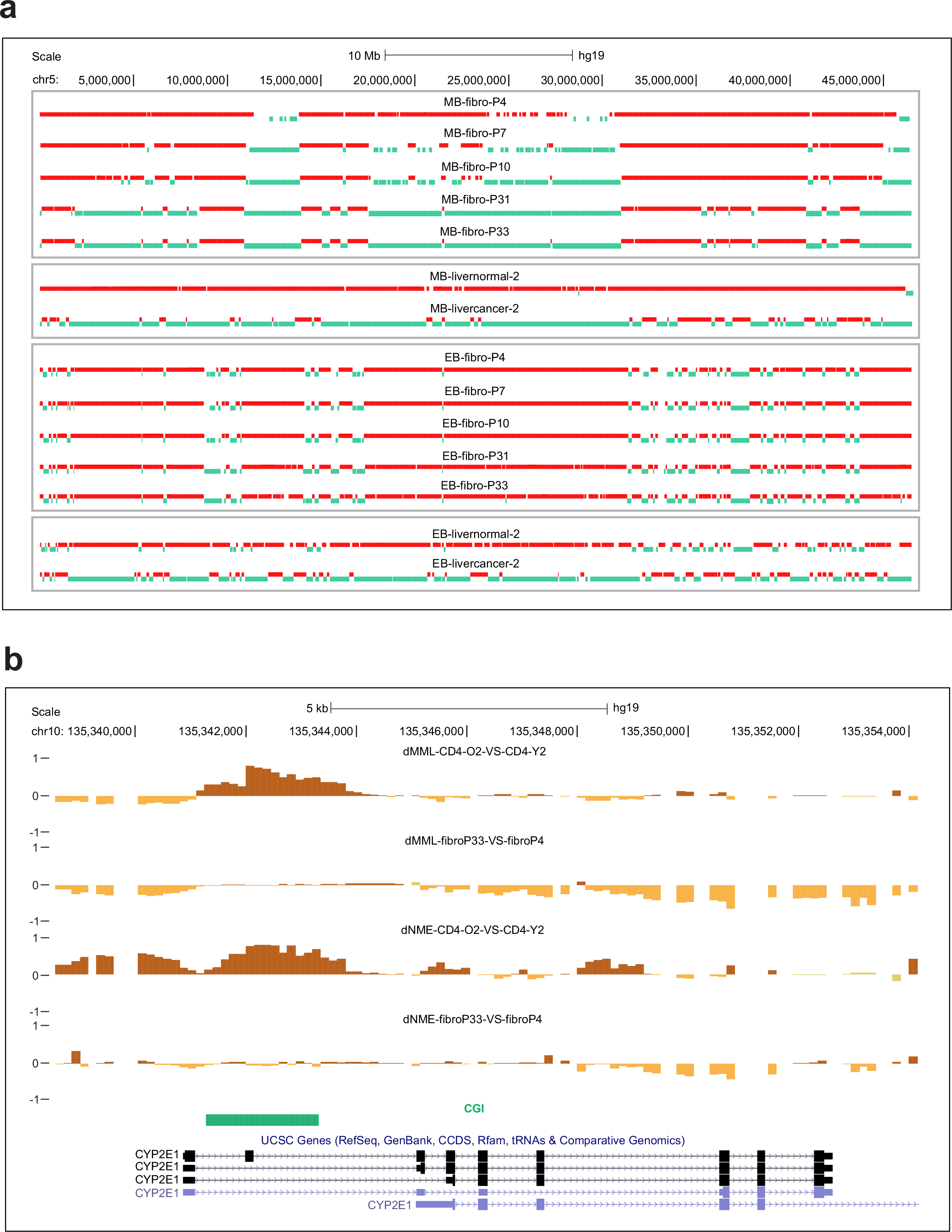
Fibroblasts may not be appropriate for modeling aging. **a**, Unmethylated blocks (MB-green) progressively form with passage in HNF fibroblasts and this process is similar to the one observed during carcinogenesis in liver cells. However, entropic blocks (EB-red) remain relatively stable. **b**, An example of the potentially misleading nature of HNF fibroblasts as a model for aging is *CYP2E1*, a gene that has been found to be downregulated with age. The differential mean methylation level (dMML) track shows methylation gain in old CD4^+^ lymphocytes near the promoter of this gene, whereas no appreciable change in methylation level is observed with passage. Similarly, the *CYP2E1* promoter demonstrates large entropy differential (dNME) in old CD4+ lymphocytes, but virtually no entropy change with passage in HNF fibroblasts. **c**, Noticeable gain in methylation entropy is also observed near the promoter of *FLNB* in old CD4^+^ lymphocytes, a gene found to be downregulated with age. However, the *FLNB* promoter exhibits loss of entropy with passage in fibroblasts.

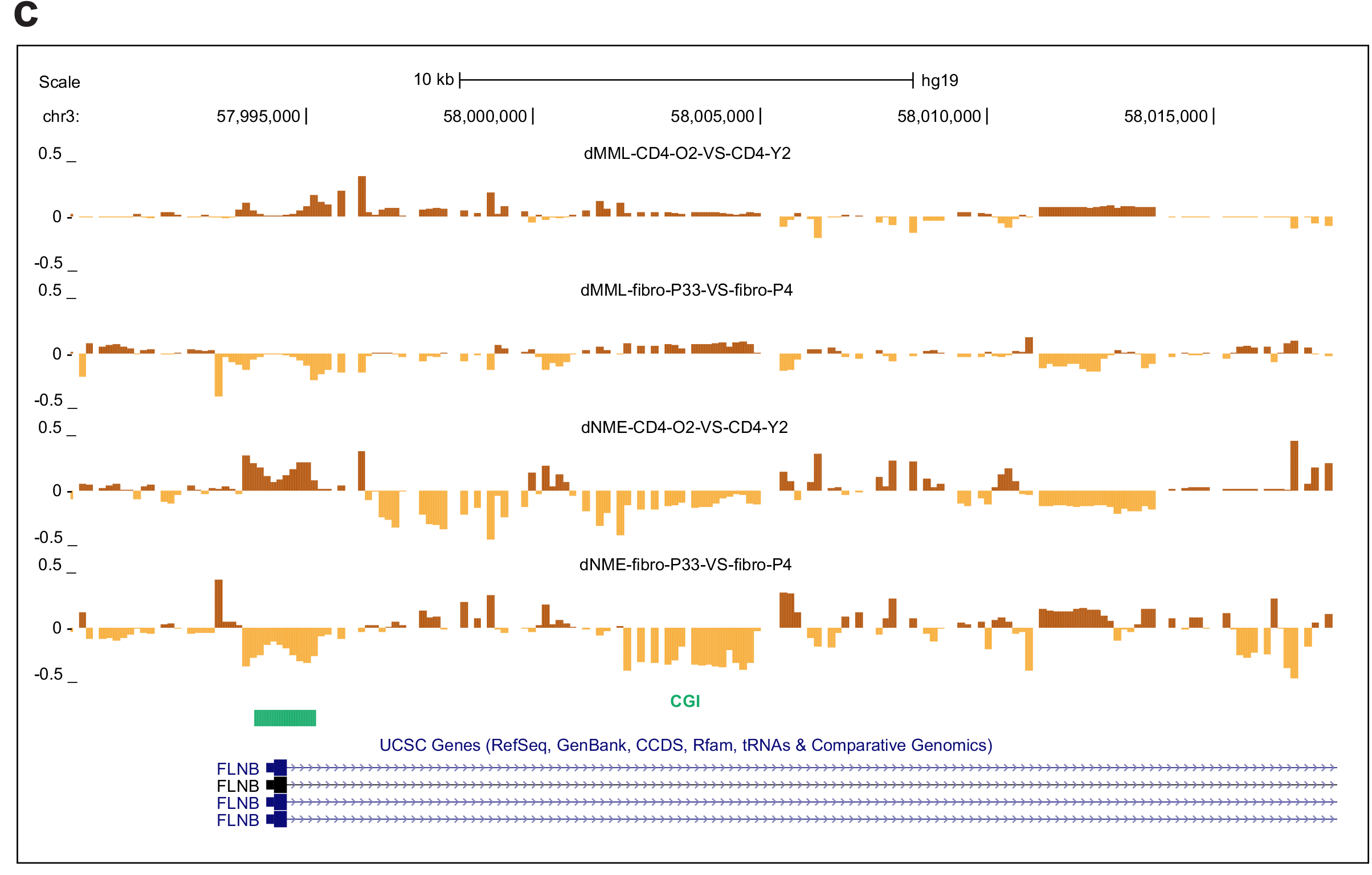

**Extended Data Figure 4.**
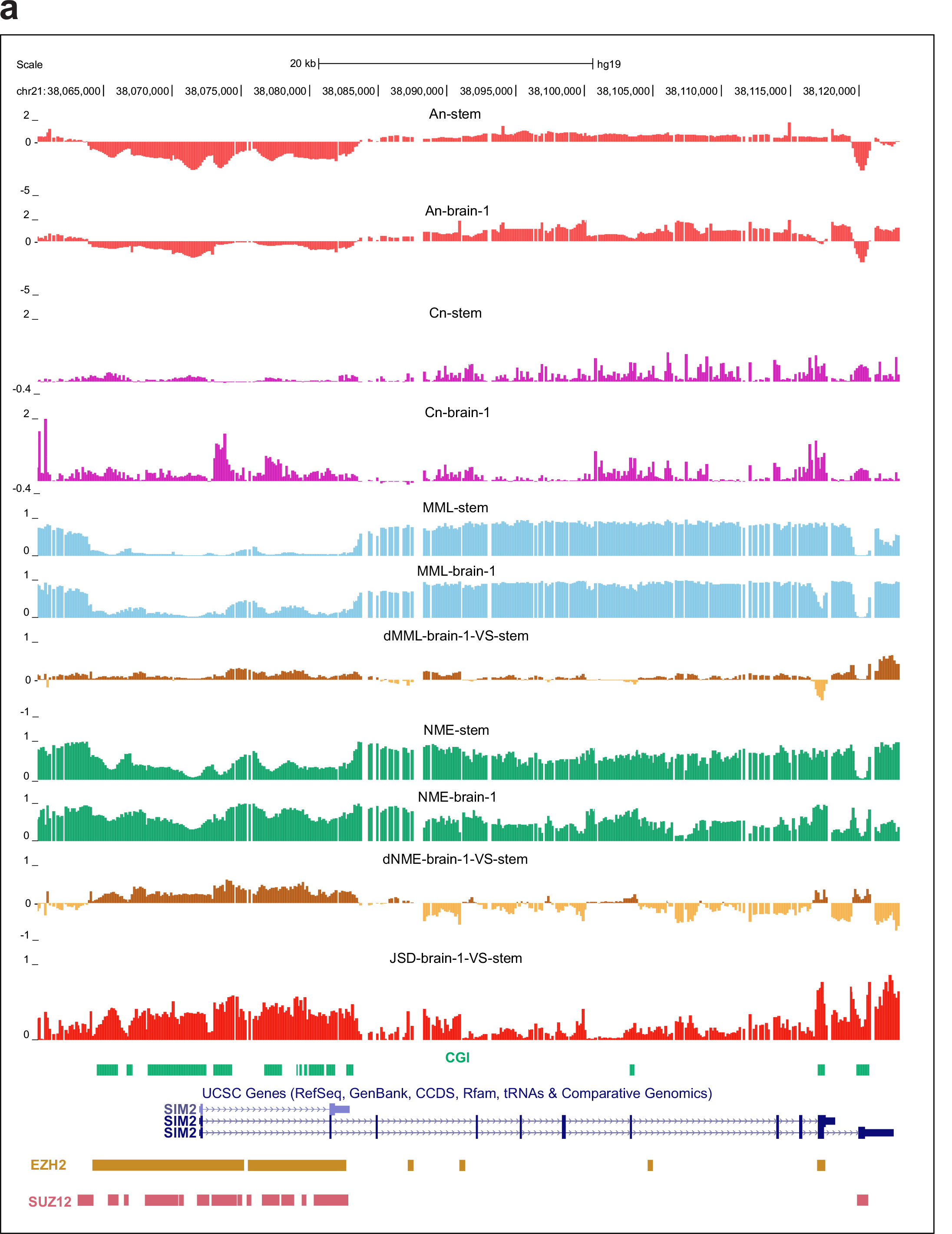
Differential regulation within genomic regions of high JSD but low dMML near promoters. **a**, The promoter of *SIM2*, a master regulation of neurogenesis, exhibits low level of differential methylation (dMML) but high JSD between stem cells and brain, demonstrating large epigenetic distance. Regulation of the PEL parameters results in low methylation level in both stem and brain but in an entropy gain in brain. This region shows binding of EZH2 and SUZ12, key components of the histone methyltransferase PRC2. **b**, A similar behavior is observed within a 14-kb region that contains *FOXD3*, a transcription factor associated with pluripotency. **c**, The promoter of *SALL1*, a key developmental gene, exhibits differential behavior between stem and brain that is similar to the one exhibited by *SIM2*. **d**, The promoter of *ASCL2*, a developmental gene involved in the determination of the neuronal precursors in the peripheral and central nervous systems, exhibits a similar behavior as the promoters of *SIM2* and *SALL1* but shows entropy loss in brain.

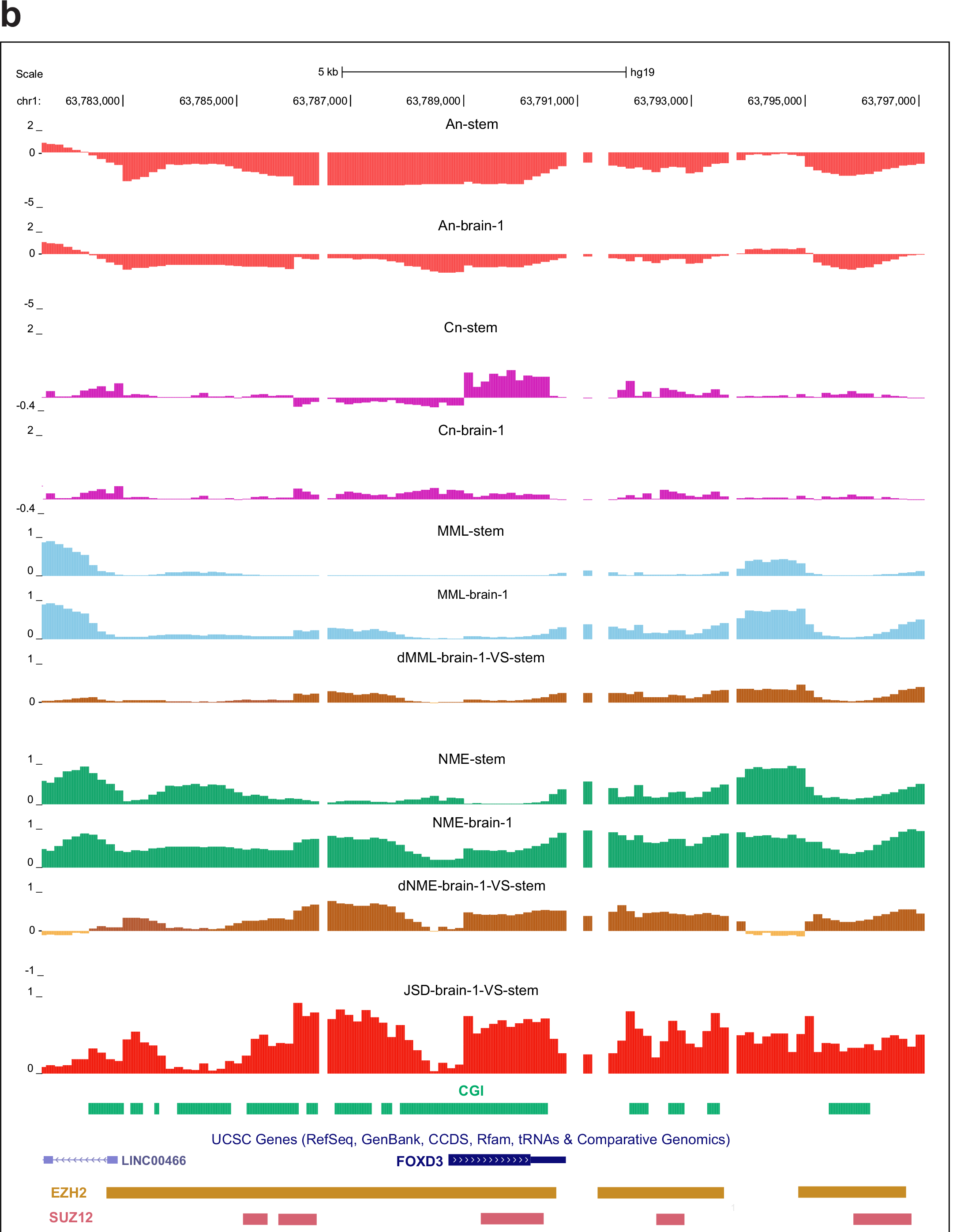

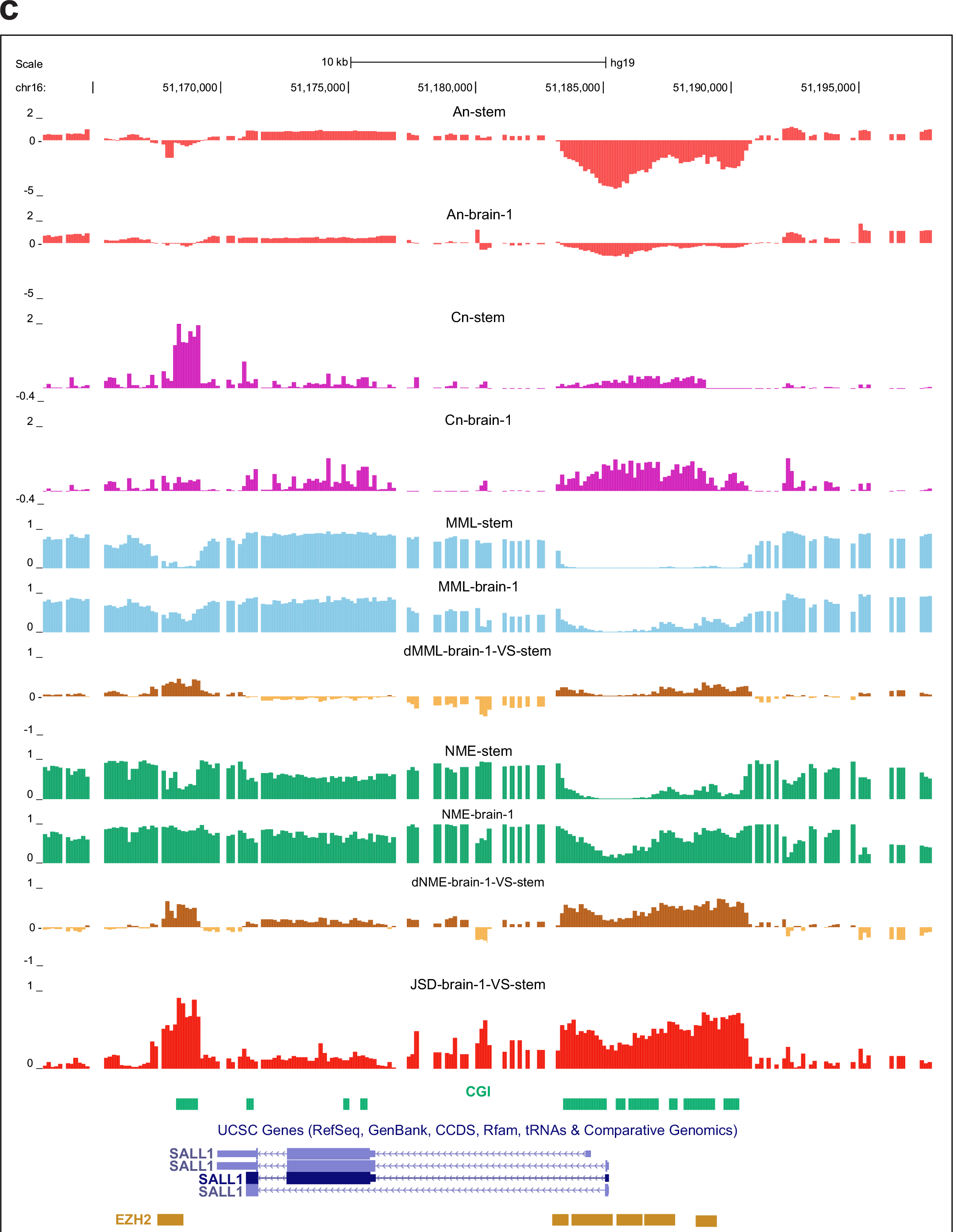

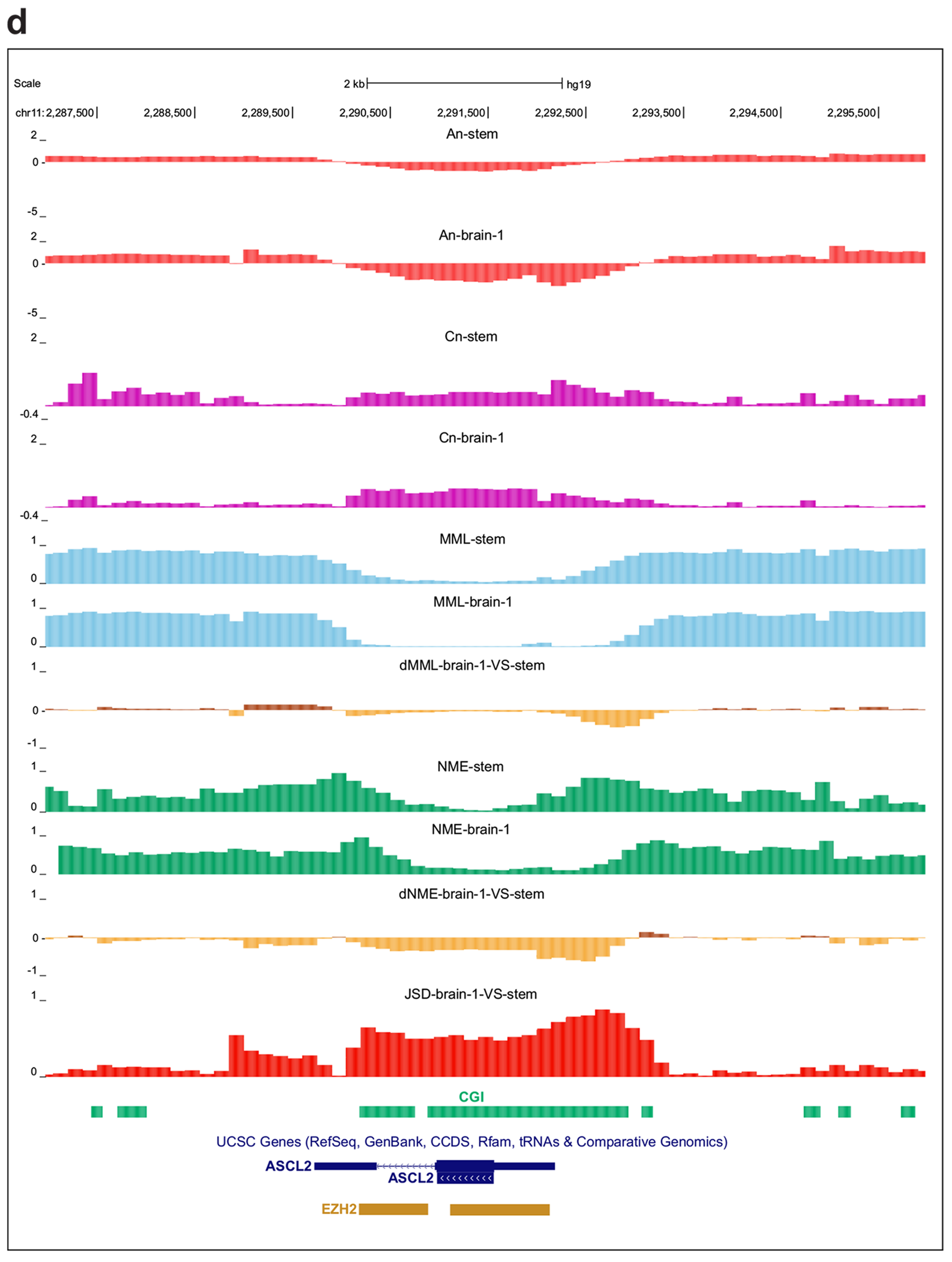

**Extended Data Figure 5.**
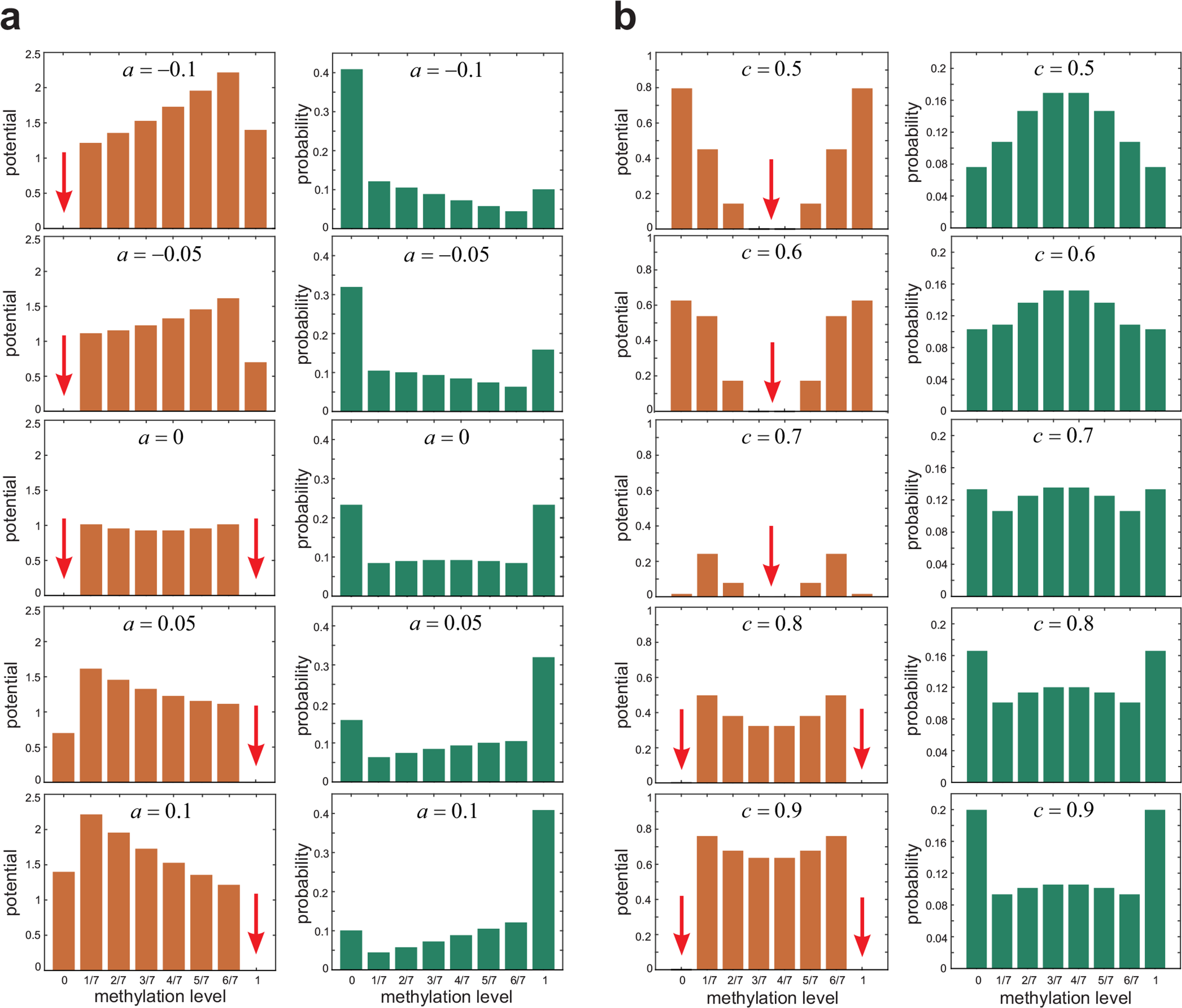
Reallocation of ground states due to PEL deformation explains (non-critical) phase transition and bistability in methylation level. **a**, Potential energy landscapes (left column) and probability distributions (right column) of methylation level within a genomic unit (GU) that contains 7 CpG sites, obtained by simulation when the PEL parameter *c* equals 1 and for five values of parameter *a*. The red arrows indicate ground states. For values of parameter *a* that are sufficiently below 0 the PEL has only one potential well, which is located at methylation level 0 (fully unmethylated state). However, as *a* approaches 0, and for sufficiently large *c* > 0, a new potential well forms at methylation level 1 (fully methylated state), which eventually achieves the same depth as the potential well at 0 and results in a bimodal probability distribution, with modes located at 0 and 1, demonstrating bistable behavior. As *a* increases away from zero, the potential well at 1 becomes deeper, whereas the potential well at 0 becomes shallower and eventually disappears. **b**, Potential energy landscapes (left column) and probability distributions (right column) of methylation level when *a* = 0 and for five values of parameter *c*. When *a* = *c* = 0, the PEL has only one ground state, since methylation of the CpG sites will be statistically independent and equiprobable in this case and the methylation level will follow a binomial distribution. However, as *c* increases away from zero, this ground state eventually disappears and two new ground states form at 0 and 1 resulting in a bimodal probability distribution, with modes located at 0 and 1, demonstrating bistable behavior.

**Extended Data Figure 6.**
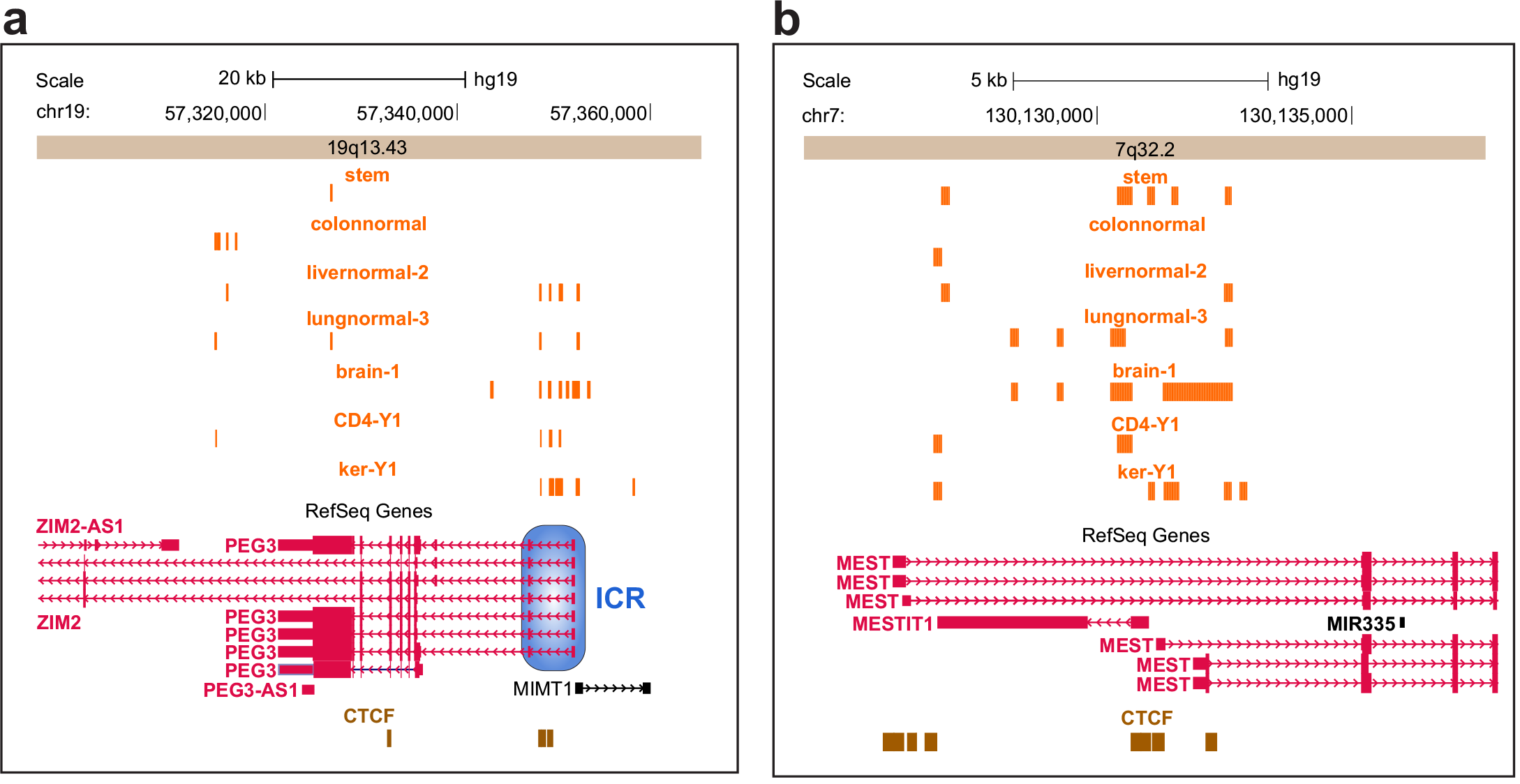
Additional examples of methylation bistability. **a**, UCSC genome browser image displaying part of the 19q13.43 chromosomal region around the *PEG3/ZIM2* promoter. Bistable methylation marks, shown for a number of normal tissues, coincide with the location of the *PEG3/ZIM2* ICR that exhibits CTCF binding. Note that the ICR also includes the transcriptional start site of the imprinted gene *MIMT1*. **b**, UCSC genome browser image displaying part of the 7q32.2 chromosomal region around the *MEST/MESTIT1* promoter. Bistable methylation marks, shown for a number of normal tissues, coincide with areas rich in CTCF binding sites.

**Extended Data Figure 7.**
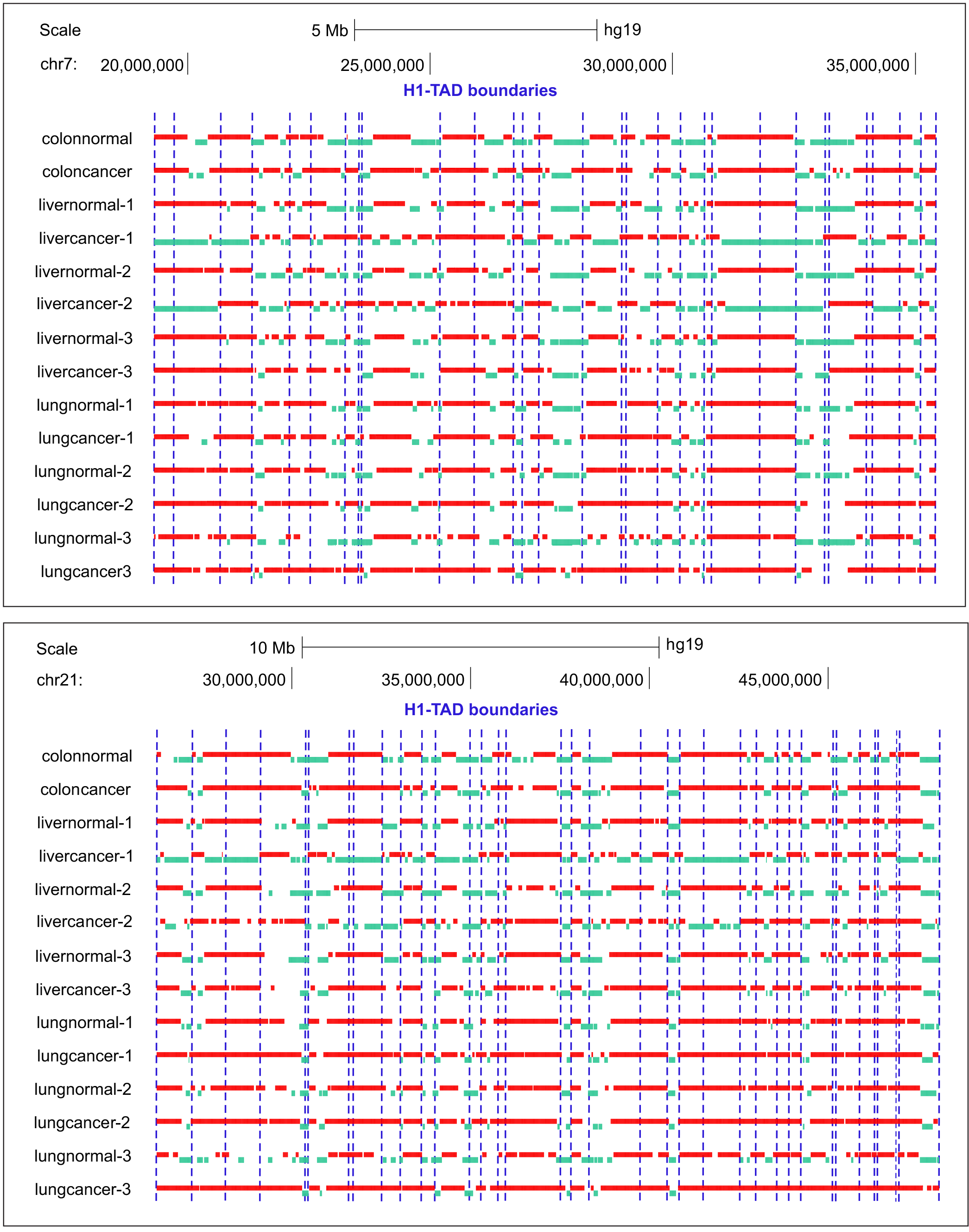
Entropy blocks and TAD boundaries. Examples of two genomic regions in chr7 and chr21 showing that the location of TAD boundaries may correlate with boundaries of ordered (green) or disordered (red) blocks.

**Extended Data Figure 8.**
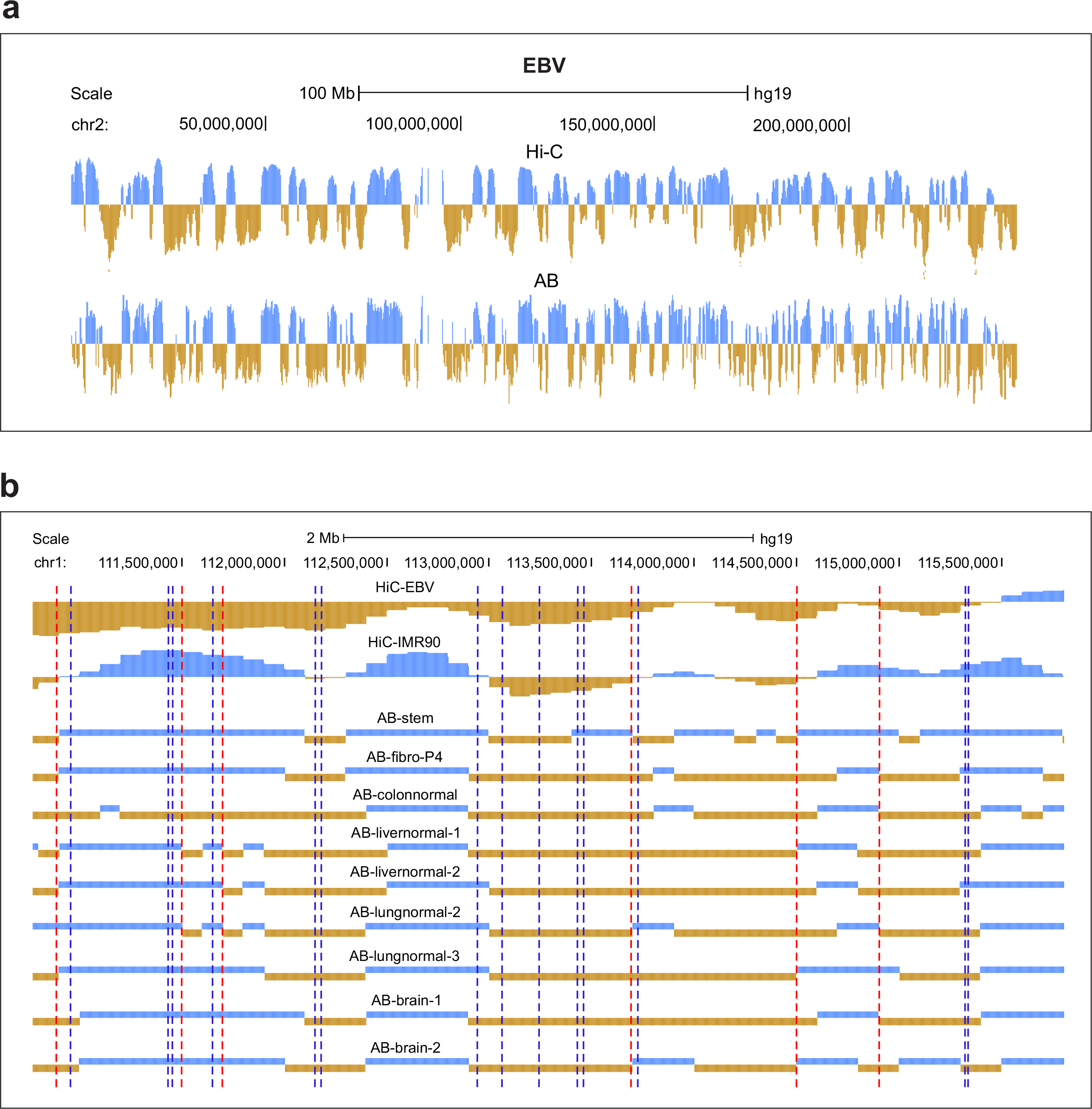
A/B compartment switching. **a**, Random forest based prediction of A/B compartments (AB) in EBV cells using information-theoretic properties of methylation maintenance. **b**, Differences in compartments A and B are observed when comparing EBV and IMR90 Hi-C data. Compartments A (brown) and B (blue), obtained from random forest based predictions of the Hi-C signal using a zero calling margin, demonstrate a similar behavior across samples. Dashed lines indicate predicted TAD boundaries with the red dashed lines corresponding to compartment transitions. **c**, Switching between predicted compartments A (brown) and B (blue) is observed in cancer, with B to A switching being more frequent than A to B switching. **d**, Although *ESR1* is located within compartment A (brown) in normal colon, liver and lung, it is relocated to compartment B (blue) in colon cancer but not in liver and lung cancer. This reorganization is accompanied by appreciable hypermethylation and entropy gain within the CGI near the gene’s promoter. *ESR1* has been implicated in colon cancer. **e**, *CYP2E1* is within compartment B (blue) in normal colon, liver and lung but it is relocated to compartment A (brown) in liver cancer. This reorganization is accompanied by hypomethylation and loss of entropy within the shores of the CGI near the gene’s promoter. *CYP2E1* has been associated with liver cancer susceptibility. **f**, Boxplots of genome-wide JSD distributions within compartments A (brown) and B (blue) in normal colon, liver and lung demonstrate gain in JSD within compartment B in cancer. The boxes show the 25% quantile, the median, and the 75% quantile, whereas each whisker has a length of 1.5x the interquartile range. **g**, The *HOXA* cluster of developmental genes is within compartment B in normal colon, liver and lung. It is however relocated to compartment A in colon and liver cancer but not in lung cancer. Compartmental reorganization of the *HOXA* genes is accompanied by marked hypomethylation and entropy loss within selected loci, implicating a role of chromatin reorganization in altered *HOXA* gene expression within tumors. **h**, The *HOXD* genes are within compartment B in normal colon, liver and lung and are relocated to compartment A in all three cancers. **i**, *SOX9* is within compartment B in colon and lung normal and is relocated to compartment B only in colon cancer. This is accompanied by marked hypomethylation and entropy loss. *SYK* is within compartment B in colon and lung normal and it is relocated to compartment B both in colon and lung cancer. **j**, *MGMT* and *MSH4* are within compartment A in colon and lung normal and they are relocated to compartment B only in colon cancer. Compartmental reorganization is accompanied mostly by hypomethylation and a marked gain in entropy.

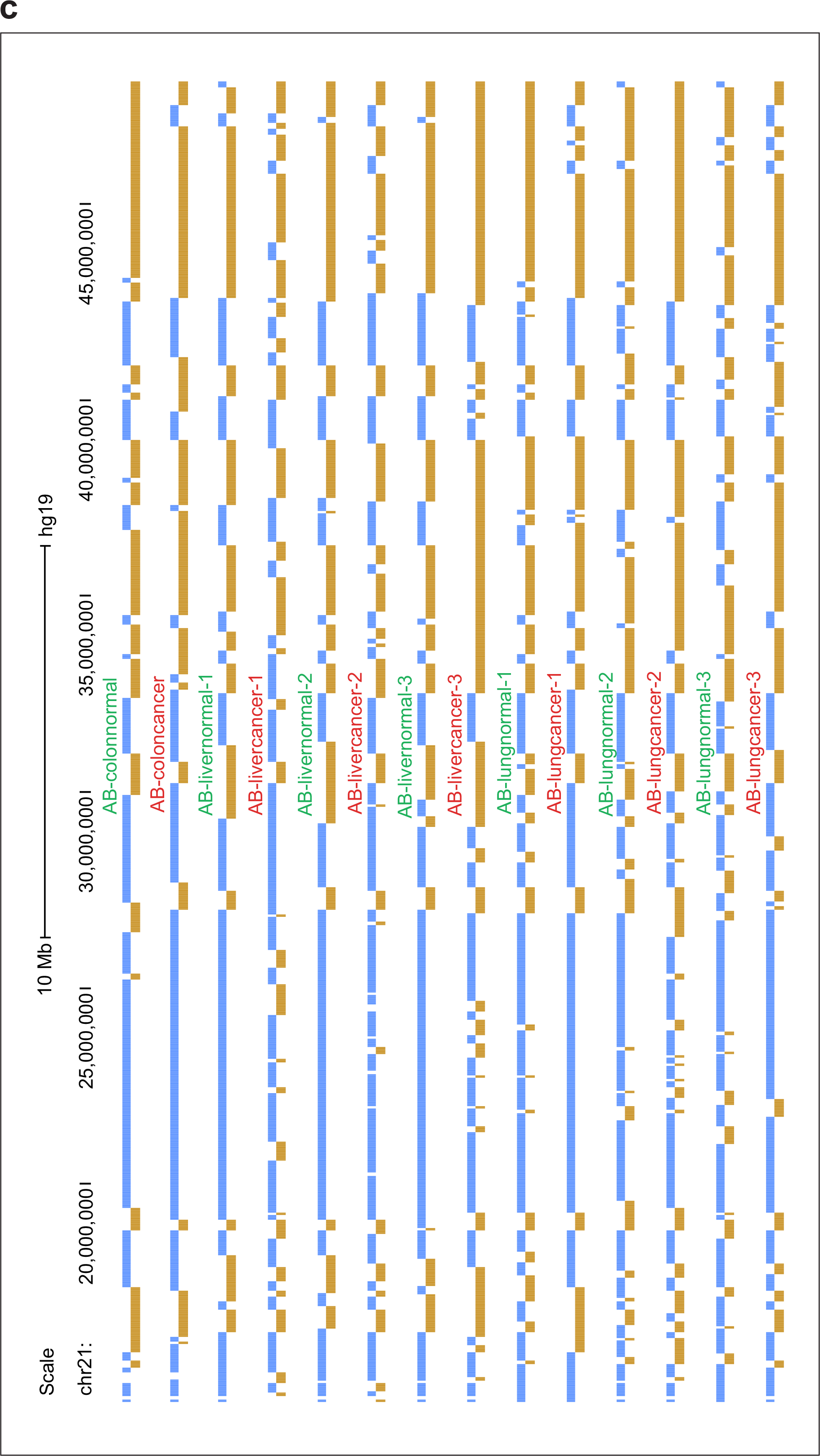

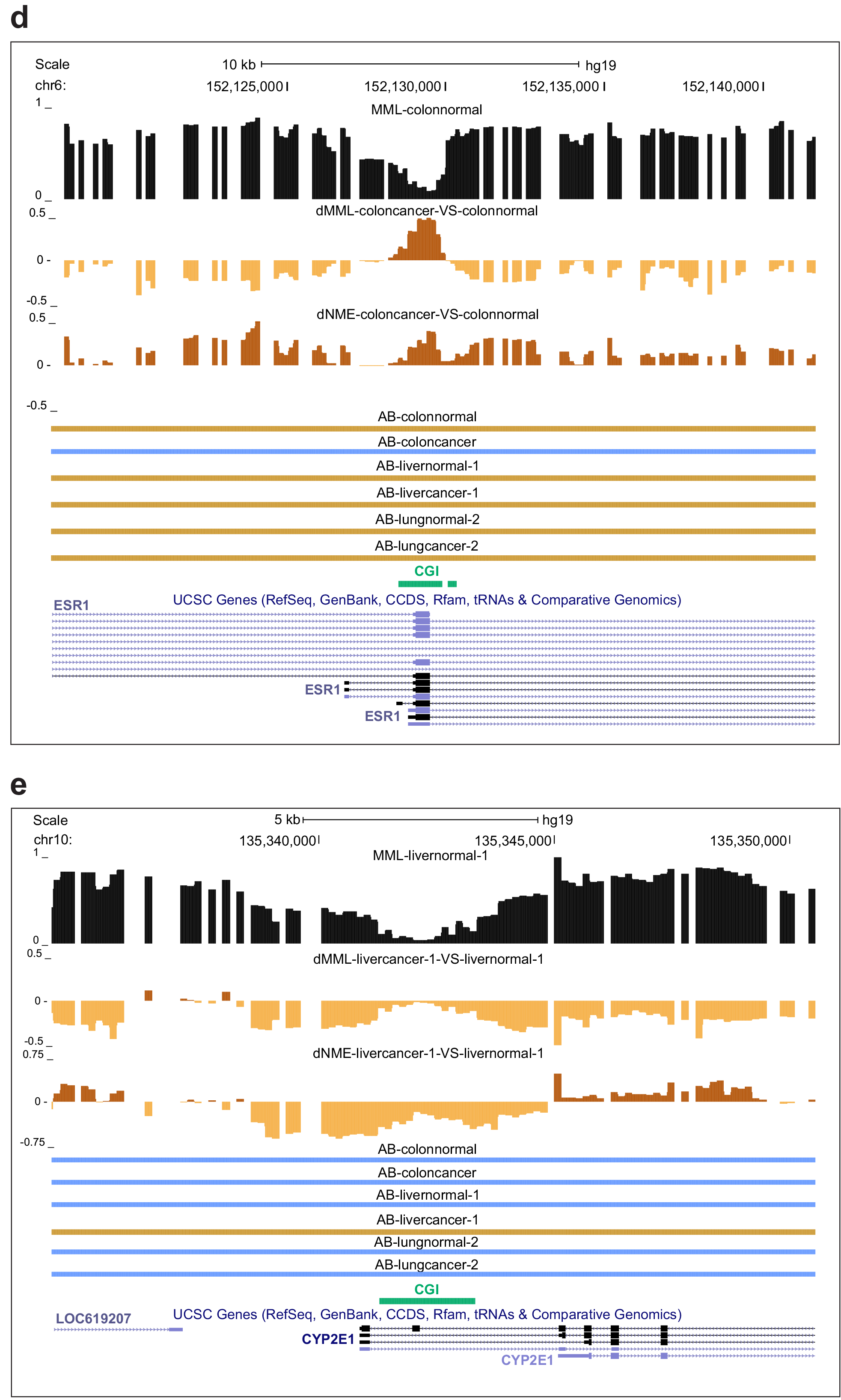

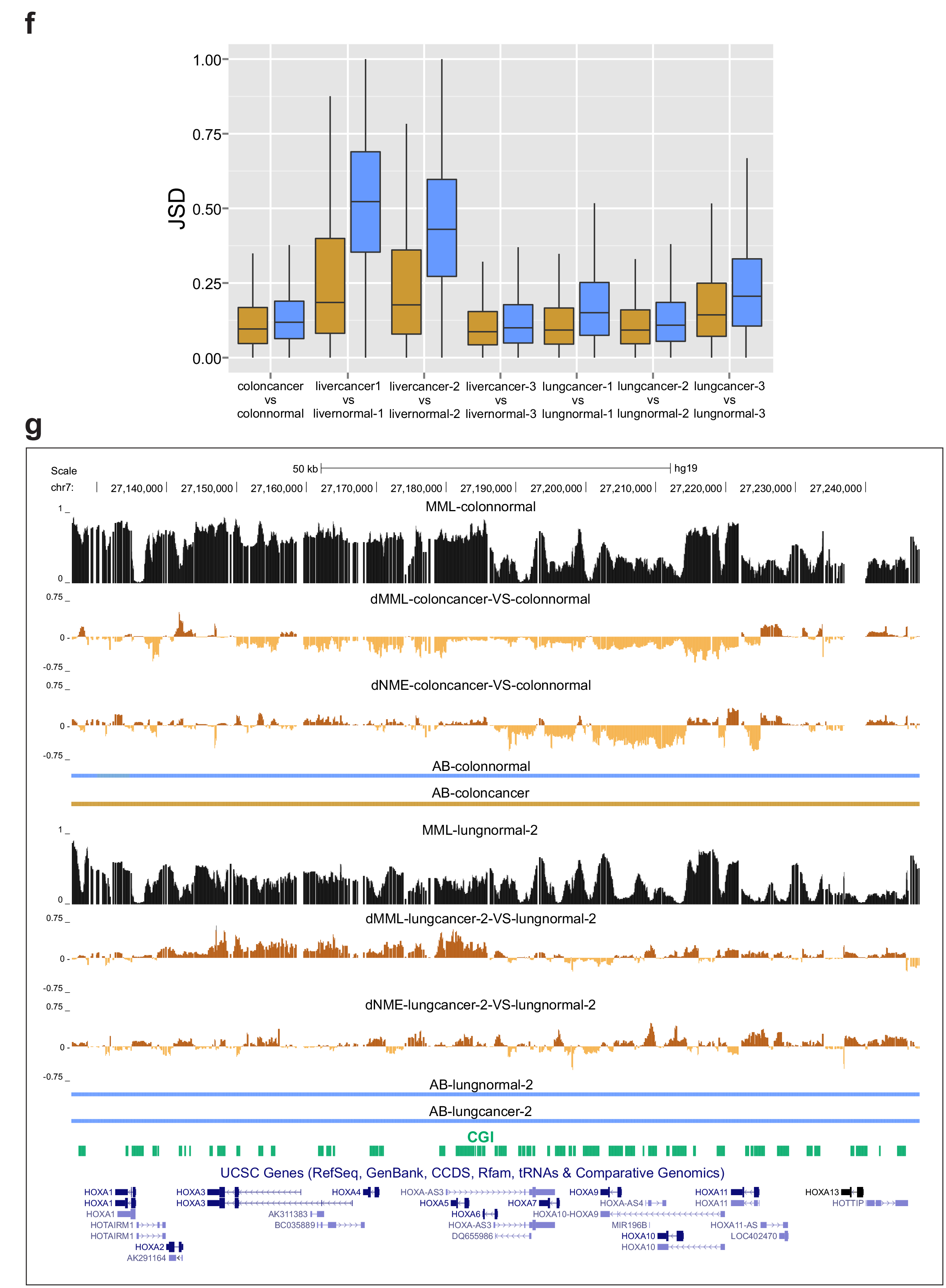

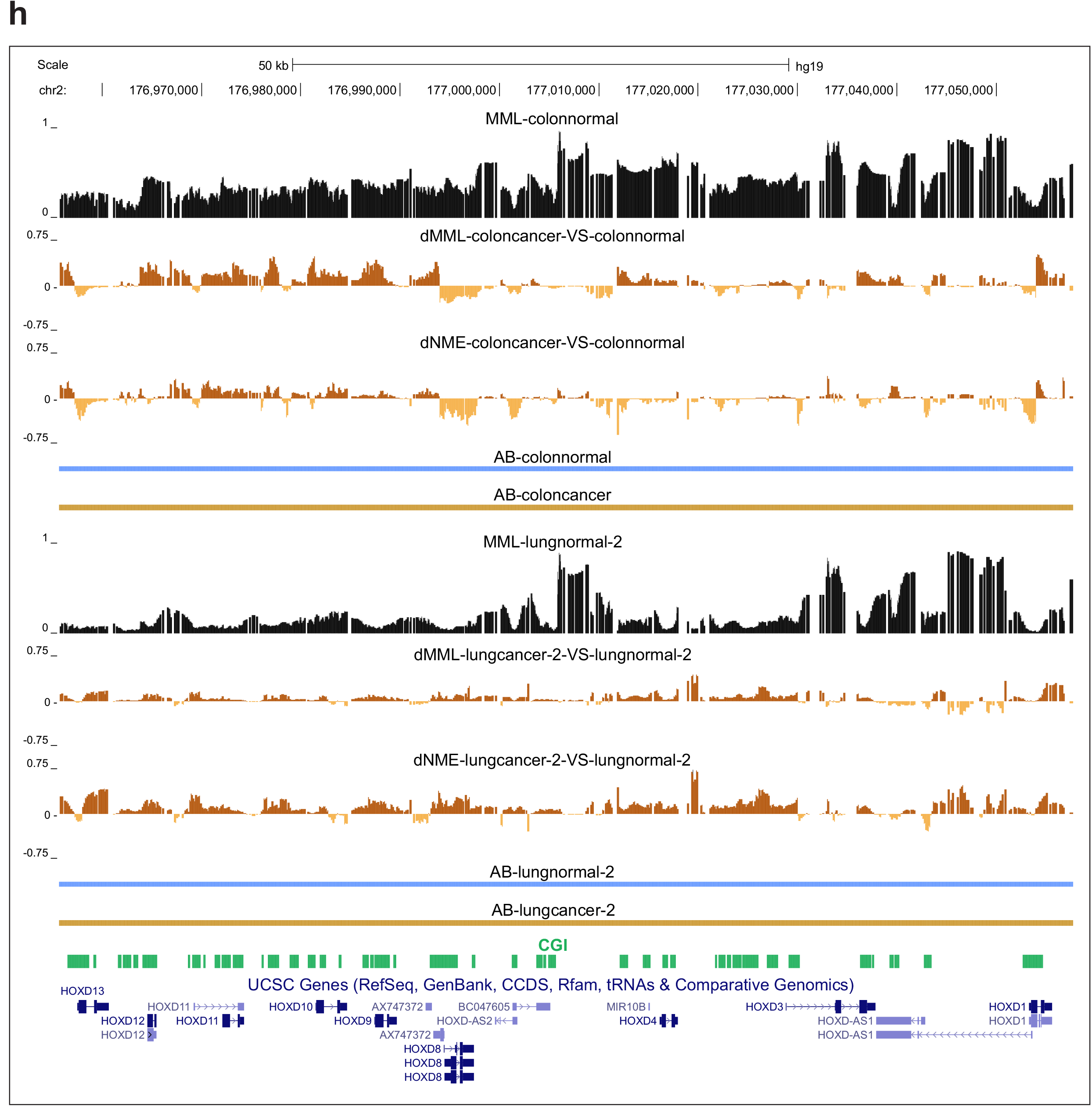

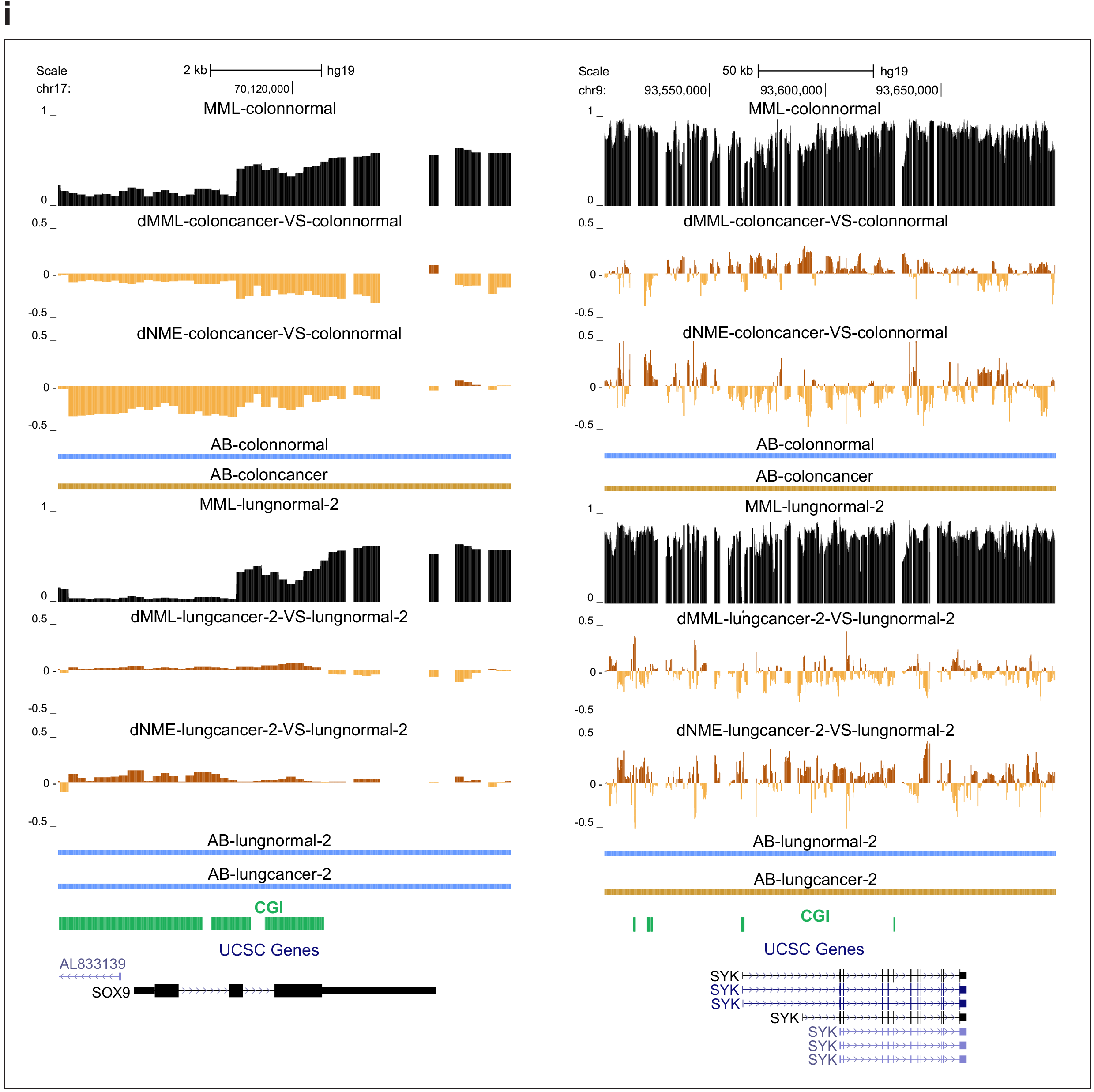

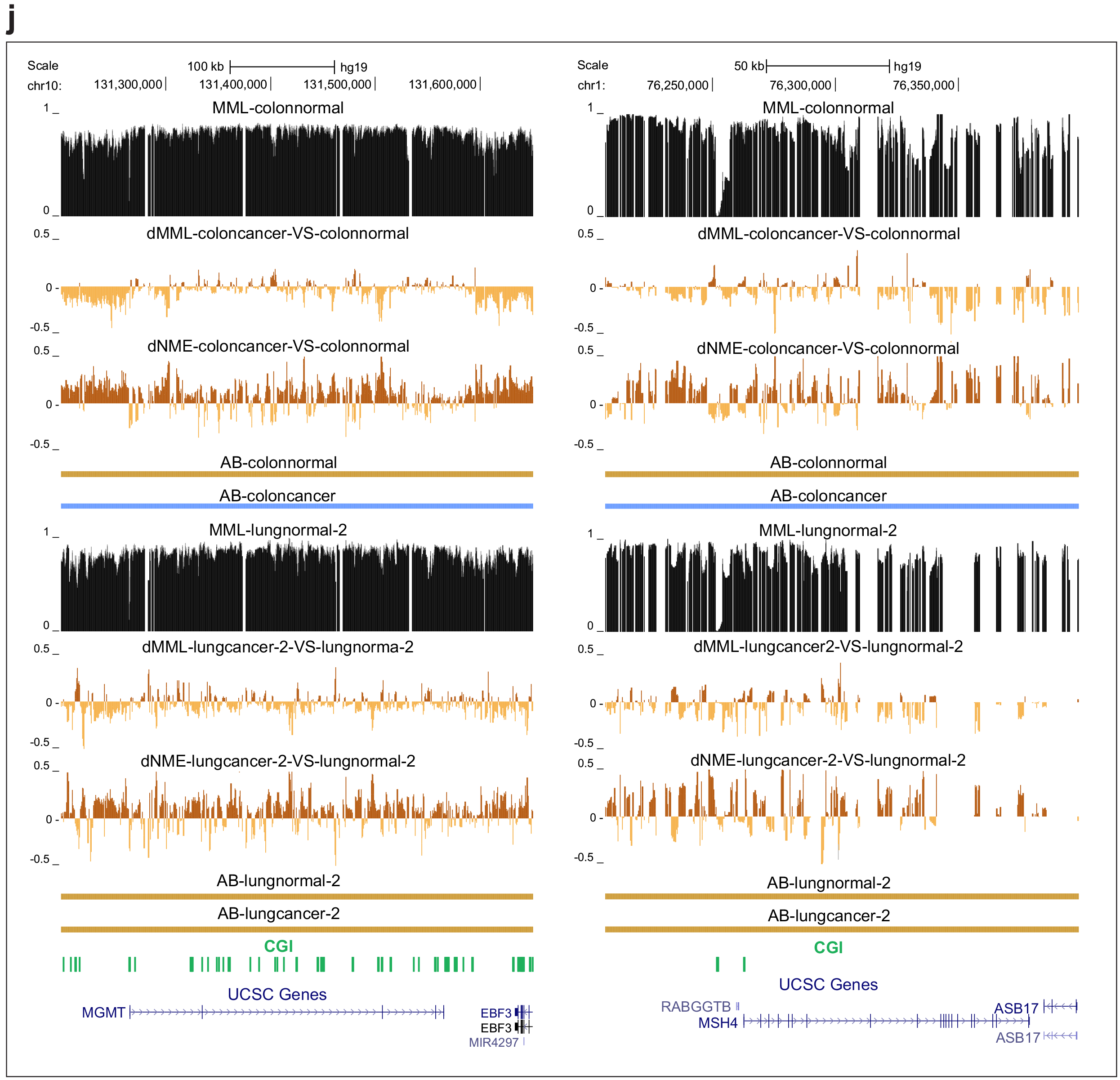

**Extended Data Figure 8.**
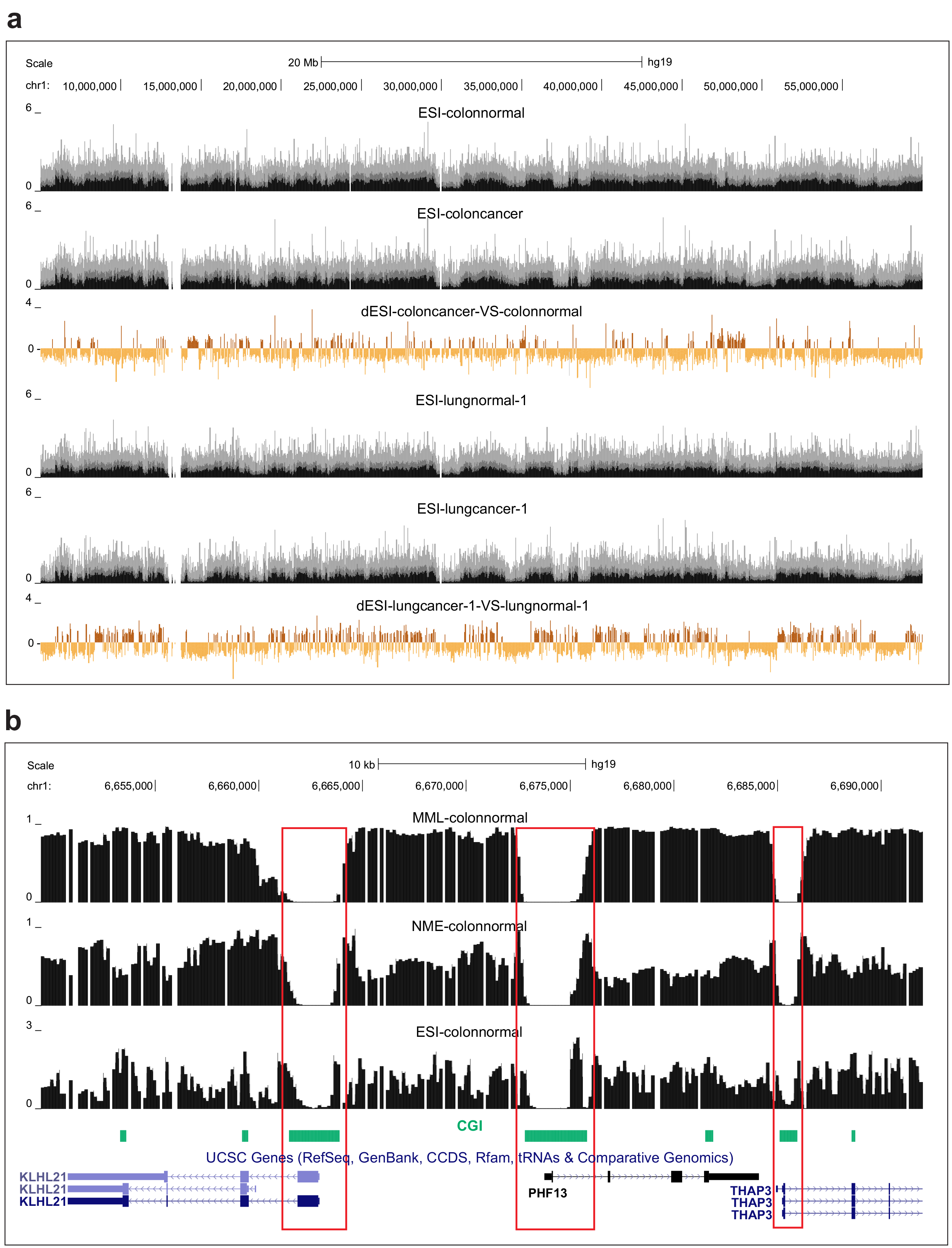
ESI distributions reveal wide behavior in entropic sensitivity. **a**, Large differences in entropic sensitivity (dESI) may be observed genome-wide between normal and cancer tissues, exhibiting alternate bands of hyposensitivity and hypersensitivity. **b**, An example of ESI values in colon normal tissue shows wide-spread entropic sensitivity along the genome. However, unmethylated CGIs may exhibit low entropic sensitivity. *KLHL21* is a substrate-specific adapter of a BCR (BTB-CUL3-RBX1) E3 ubiquitin-protein ligase complex required for efficient chromosome alignment and cytokinesis. *PHF13* regulates chromatin structure. *THAP3* is required for regulation of *RRM1* that may play a role in malignancies and disease. **c**, In liver normal cells, substantial entropic sensitivity is observed within the CGI near the promoter of the polycomb target gene *ENSA*, which is significantly reduced in liver cancer. *ENSA* is known to be hypomethylated in liver cancer. **d**, In lung normal cells, the CGI near the promoter of *NEU1* exhibits low entropic sensitivity, which is significantly increased in lung cancer. *NEU1* sialidase is required for normal lung development and function, whereas its expression has been implicated in tumorigenesis and metastatic potential. **e**, In young CD4^+^ lymphocytes, substantial entropic sensitivity is observed within the CGI near the promoter of *CYP2E1*, which is lost in old individuals. *CYP2E1* is known to be downregulated with age. **f**, The CGI near the promoter of *FLNB* exhibits gain in entropic sensitivity in old CD4^+^ lymphocytes. *FLNB* is known to be downregulated with age.

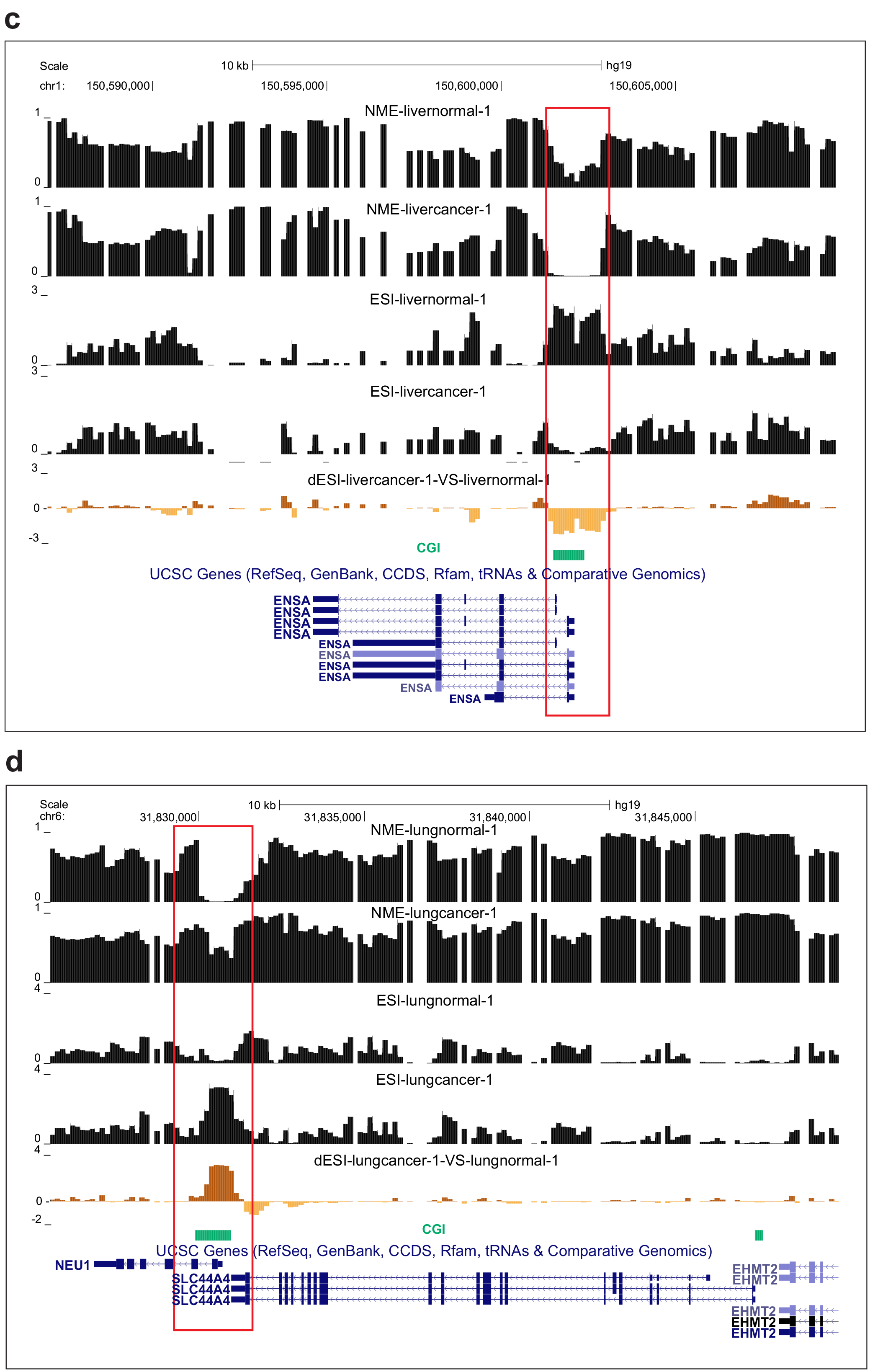

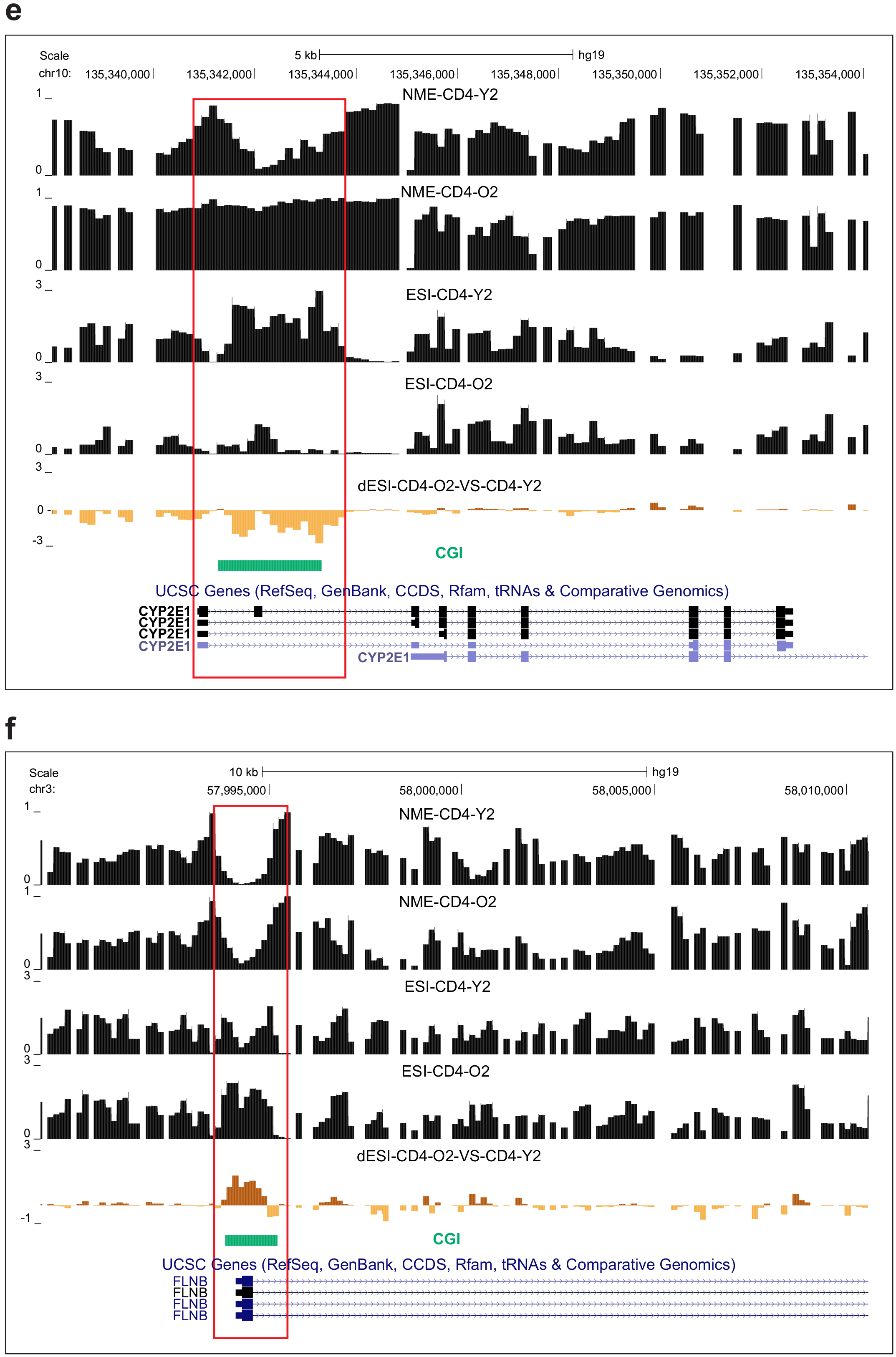

**Supplementary Table 1.** WGBS data samples.

| NICKNAME | MATCHED | SAMPLE TYPE | SOURCE<sup>1</sup> | COVERAGE |
| --- | --- | --- | --- | --- |
| **Stem Cells** |  |  |  |  |
| stem |  | H1 human embryonic stem cell line | [1], SRP072141<sup>2</sup> | 24 |
| **Normal / Cancer** |  |  |  |  |
| colonnormal | 1 | colon normal | [2] | 30 |
| coloncancer | 1 | colon cancer | [2] | 30 |
| livernormal-1 | 2 | liver normal | SRP072078 | 9 |
| livercancer-1 | 2 | liver cancer | SRP072078 | 8 |
| livernormal-2 | 3 | liver normal | SRP072078 | 7 |
| livercancer-2 | 3 | liver cancer | SRP072078 | 8 |
| livernormal-3 | 4 | liver normal | SRP072078 | 18 |
| livercancer-3 | 4 | liver cancer | SRP072078 | 18 |
| livernormal-4 |  | liver normal | [2] | 60 |
| livernormal-5 |  | liver normal | [2] | 41 |
| lungnormal-1 | 5 | lung normal | SRP072078 | 14 |
| lungcancer-1 | 5 | lung cancer | SRP072078 | 15 |
| lungnormal-2 | 6 | lung normal | SRP072078 | 10 |
| lungcancer-2 | 6 | lung cancer | SRP072078 | 10 |
| lungnormal-3 | 7 | lung normal | SRP072078 | 19 |
| lungcancer-3 | 7 | lung cancer | SRP072078 | 18 |
| brain-1 |  | post-mortem brain, pre-frontal cortex, normal | SRP072071 | 11 |
| brain-2 |  | post-mortem brain, pre-frontal cortex, normal | SRP072071 | 12 |
| **HNF Fibroblasts** |  |  |  |  |
| fibro-P4 |  | human neonatal fibroblasts, passage 4 | SRP072075 | 12 |
| fibro-P7 |  | human neonatal fibroblasts, passage 7 | SRP072075 | 11 |
| fibro-P10 |  | human neonatal fibroblasts, passage 10 | SRP072075 | 11 |
| fibro-P31 |  | human neonatal fibroblasts, passage 31 | SRP072075 | 11 |
| fibro-P33 |  | human neonatal fibroblasts, passage 33. senescent | SRP072075 | 11 |
| **CD4 T-Cells** |  |  |  |  |
| CD4-Y1 |  | flow-sorted peripheral CD4 T-cells from an 18 year old female | SRP072075 | 8 |
| CD4-Y2 |  | flow-sorted peripheral CD4 T-cells from a 25 year old female | SRP072075 | 8 |
| CD4-Y3 |  | flow-sorted peripheral CD4 T-cells from a 25 year old female | SRP072075 | 7 |
| CD4-O1 |  | flow-sorted peripheral CD4 T-cells from an 82 year old female | SRP072075 | 7 |
| CD4-O2 |  | flow-sorted peripheral CD4 T-cells from an 82 year old female | SRP072075 | 8 |
| CD4-O3 |  | flow-sorted peripheral CD4 T-cells from an 86 year old female | SRP072075 | 7 |
| **Keratinocytes** |  |  |  |  |
| ker-Y1 |  | keratinocytes from a skin biopsy of a sun-protected site on a young individual | [3] | 8 |
| ker-Y2 |  | keratinocytes from a skin biopsy of a sun-protected site on a young individual | [3] | 8 |
| ker-O1 |  | keratinocytes from a skin biopsy of a sun-exposed site on an older individual | [3] | 7 |
| ker-O2 |  | keratinocytes from a skin biopsy of a sun-exposed site on an older individual | [3] | 7 |
| **EBV** |  |  |  |  |
| EBV |  | EBV-immortalized lymphoblasts | [4] | 9 |

<sup>1</sup>SRP accessions correspond to NCBI Sequencing Read Archive (SRA).
<sup>2</sup>Original sequence along with additional coverage have been deposited in the reference SRP accession.

**Supplementary Table 2.** Statistical analysis results for EZH2/SUZ12 binding association with promoters and enhancers at high JSD genomic loci.

| FISHER'S EXACT TEST FOR COUNT DATA |  |  |  |  |  |  |  |  |  |  |  |
| --- | --- | --- | --- | --- | --- | --- | --- | --- | --- | --- | --- |
| PROMOTERS |  |  |  |  |  |  |  |  |  |  |  |
|  |  | EZH2 |  |  |  |  | SUZ12 |  |  |  |  |
| criterion | # genes | present | absent | frequency | P value | odds ratio | present | absent | frequency | P value | odds ratio |
| dMML | top 1000 | 305 | 695 | 31% | < 2.2E-16 | 2.69 | 94 | 906 | 9% | 2.05E-05 | 2.20 |
|  | bottom 1000 | 140 | 860 | 14% |  |  | 45 | 955 | 5% |  |  |
| JSD | top 1000 | 457 | 543 | 46% | < 2.2E-16 | 7.57 | 191 | 809 | 19% | < 2.2E-16 | 8.84 |
|  | bottom 1000 | 100 | 900 | 10% |  |  | 26 | 974 | 3% |  |  |
| ENHANCERS |  |  |  |  |  |  |  |  |  |  |  |
|  |  | EZH2 |  |  |  |  | SUZ12 |  |  |  |  |
| criterion | # genes | present | absent | frequency | P value | odds ratio | present | absent | frequency | P value | odds ratio |
| dMML | top 100 | 42 | 58 | 42% | 7.24E-13 | 34.95 | 29 | 71 | 29% | 6.20E-09 | 39.92 |
|  | bottom 100 | 2 | 98 | 2% |  |  | 1 | 99 | 1% |  |  |
| JSD | top 100 | 53 | 47 | 53% | < 2.2E-16 | 109.49 | 40 | 60 | 40% | 1.34E-14 | infinite |
|  | bottom 100 | 1 | 99 | 1% |  |  | 0 | 100 | 0% |  |  |

| BINOMIAL LOGISTIC REGRESSION |  |  |  |  |  |  |  |  |  |
| --- | --- | --- | --- | --- | --- | --- | --- | --- | --- |
| PROMOTERS |  |  |  |  |  |  |  |  |  |
|  |  | EZH2 |  |  |  | SUZ12 |  |  |  |
|  |  | coefficient | std error | P value | holdout accuracy* | coefficient | std error | P value | holdout accuracy* |
| JSD | intercept | -2.4030 | 0.0395 | < 2.2E-16 | 82% | -3.9217 | 0.0638 | < 2.2E-16 | 95% |
|  | score | 5.5511 | 0.1991 | < 2.2E-16 |  | 6.1825 | 0.2760 | < 2.2E-16 |  |
| ENHANCERS |  |  |  |  |  |  |  |  |  |
|  |  | EZH2 |  |  |  | SUZ12 |  |  |  |
|  |  | coefficient | std error | P value | holdout accuracy* | coefficient | std error | P value | holdout accuracy* |
| JSD | intercept | -4.3962 | 0.2914 | < 2.2E-16 | 88% | -6.4587 | 0.5133 | < 2.2E-16 | 93% |
|  | score | 18.1070 | 1.7861 | < 2.2E-16 |  | 23.0143 | 2.4591 | < 2.2E-16 |  |

\*90% of data was randomly selected for training, while the remaining was used for estimating performance.

**Supplementary Table 3.** Odds ratio analysis results of bistability enrichment in CGIs, shores, promoters, and gene bodies.

| SAMPLE | CGIs |  | SHORES |  | PROMOTERS |  | GENE BODIES |  |
| --- | --- | --- | --- | --- | --- | --- | --- | --- |
|  | OR | P value | OR | P value | OR | P value | OR | P value |
| stem | 1.03 | 5.19E-01 | 4.34 | 0.00E+00 | 4.22 | 0.00E+00 | 0.90 | 3.06E-14 |
| colonnormal | 0.41 | 4.26E-190 | 1.54 | 0.00E+00 | 1.69 | 0.00E+00 | 0.72 | 0.00E+00 |
| coloncancer | 0.26 | 0.00E+00 | 0.94 | 1.21E-21 | 0.90 | 9.45E-42 | 0.63 | 0.00E+00 |
| livernormal-1 | 0.25 | 0.00E+00 | 1.19 | 1.22E-78 | 1.17 | 3.74E-51 | 0.67 | 0.00E+00 |
| livercancer-1 | 0.23 | 0.00E+00 | 1.30 | 4.20E-166 | 1.21 | 1.34E-62 | 0.84 | 1.43E-158 |
| livernormal-2 | 0.24 | 0.00E+00 | 1.17 | 2.12E-58 | 1.11 | 5.08E-21 | 0.68 | 0.00E+00 |
| livercancer-2 | 0.30 | 0.00E+00 | 1.28 | 1.01E-214 | 1.06 | 1.68E-09 | 0.74 | 0.00E+00 |
| livernormal-3 | 0.26 | 0.00E+00 | 1.28 | 1.73E-143 | 1.24 | 1.66E-83 | 0.71 | 0.00E+00 |
| livercancer-3 | 0.38 | 1.03E-249 | 1.42 | 1.58E-306 | 1.43 | 1.57E-253 | 0.76 | 0.00E+00 |
| livernormal-4 | 0.44 | 1.25E-145 | 1.64 | 0.00E+00 | 1.92 | 0.00E+00 | 0.81 | 9.69E-172 |
| livernormal-5 | 0.49 | 3.51E-120 | 2.01 | 0.00E+00 | 2.24 | 0.00E+00 | 0.89 | 1.46E-59 |
| lungnormal-1 | 0.35 | 9.42E-219 | 1.77 | 0.00E+00 | 1.70 | 0.00E+00 | 0.83 | 3.26E-153 |
| lungcancer-1 | 0.25 | 0.00E+00 | 1.10 | 5.33E-50 | 0.78 | 2.70E-189 | 0.60 | 0.00E+00 |
| lungnormal-2 | 0.34 | 1.47E-219 | 1.68 | 0.00E+00 | 1.64 | 0.00E+00 | 0.84 | 2.50E-125 |
| lungcancer-2 | 0.21 | 0.00E+00 | 1.15 | 3.64E-57 | 1.10 | 2.17E-19 | 0.70 | 0.00E+00 |
| lungnormal-3 | 0.39 | 2.38E-176 | 1.80 | 0.00E+00 | 1.73 | 0.00E+00 | 0.89 | 2.47E-54 |
| lungcancer-3 | 0.23 | 0.00E+00 | 0.97 | 9.14E-07 | 0.70 | 0.00E+00 | 0.62 | 0.00E+00 |
| brain-1 | 1.06 | 7.62E-02 | 3.46 | 0.00E+00 | 3.27 | 0.00E+00 | 1.45 | 6.95E-293 |
| brain-1 | 1.07 | 3.36E-02 | 3.48 | 0.00E+00 | 3.39 | 0.00E+00 | 1.38 | 7.61E-217 |
| fibro-P4 | 0.20 | 0.00E+00 | 0.89 | 3.23E-41 | 0.84 | 6.04E-67 | 0.59 | 0.00E+00 |
| fibro-P7 | 0.19 | 0.00E+00 | 0.81 | 1.15E-147 | 0.76 | 2.39E-184 | 0.57 | 0.00E+00 |
| fibro-P10 | 0.18 | 0.00E+00 | 0.81 | 2.02E-151 | 0.74 | 9.99E-218 | 0.57 | 0.00E+00 |
| fibro-P31 | 0.27 | 0.00E+00 | 1.15 | 3.15E-93 | 0.89 | 1.19E-39 | 0.68 | 0.00E+00 |
| fibro-P33 | 0.27 | 0.00E+00 | 1.18 | 1.46E-114 | 0.91 | 3.21E-24 | 0.68 | 0.00E+00 |
| CD4-Y1 | 1.26 | 6.01E-10 | 2.84 | 0.00E+00 | 2.93 | 0.00E+00 | 1.04 | 1.43E-03 |
| CD4-Y2 | 1.17 | 2.62E-05 | 2.71 | 0.00E+00 | 2.74 | 0.00E+00 | 1.00 | 9.26E-01 |
| CD4-Y3 | 0.89 | 1.50E-03 | 2.50 | 0.00E+00 | 2.52 | 0.00E+00 | 1.11 | 2.82E-27 |
| CD4-O1 | 0.68 | 1.46E-25 | 1.68 | 0.00E+00 | 1.83 | 0.00E+00 | 0.77 | 4.72E-200 |
| CD4-O2 | 0.94 | 1.41E-01 | 2.18 | 0.00E+00 | 2.25 | 0.00E+00 | 0.85 | 4.23E-61 |
| CD4-O3 | 0.93 | 8.54E-02 | 2.01 | 0.00E+00 | 2.11 | 0.00E+00 | 0.84 | 1.76E-76 |
| ker-Y1 | 0.63 | 3.54E-48 | 2.04 | 0.00E+00 | 1.93 | 0.00E+00 | 0.94 | 1.90E-15 |
| ker-Y2 | 0.66 | 4.17E-36 | 2.05 | 0.00E+00 | 1.90 | 0.00E+00 | 0.94 | 3.53E-16 |
| ker-O1 | 0.61 | 6.39E-53 | 1.82 | 0.00E+00 | 1.65 | 0.00E+00 | 0.86 | 2.62E-112 |
| ker-O2 | 0.40 | 1.92E-212 | 1.39 | 0.00E+00 | 1.22 | 5.98E-84 | 0.72 | 0.00E+00 |

depletion: OR < 1
enrichment: OR > 1

## Supplementary Methods

### 1 The parameters of the methylation potential energy landscape

Determining the methylation PEL, given by Eq. 4 in the Main Text, requires that we compute appropriate values for the 2*N −* 1 Ising parameters *a_n_* and *c_n_* from WGBS data, which can be a prohibitively large number of parameters for reliable estimation. To address this issue, we set

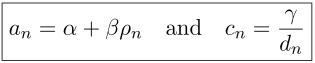

where *ρ_n_* is the CpG density (CGD) within a symmetric neighborhood of 1,000 nucleotides centered at a CpG site *n*, given by

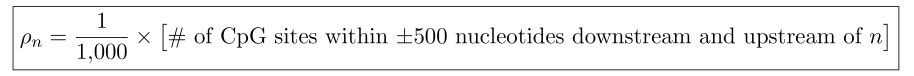

and *d_n_* is the distance of CpG site *n* from its “nearest-neighbor” CpG site *n* − 1, defined by [^1^ For biochemical interactions with the methylation machinery, the most relevant distance between two successive CpG sites is the physical distance among their cytosines. For example, if the DNA sequence under consideration is TCGACCG, then *d_2_* = 4, whereas *d_2_* = 2 for the sequence TCGCG.]

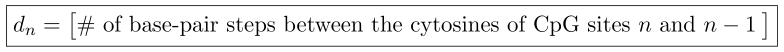

Parameter *α* accounts for intrinsic factors that uniformly affect CpG methylation over a given genomic region, whereas parameter *β* modulates the influence of CGD on methylation [^2^ It is known that DNA methylation depends on the CpG density For example, DNA regions with high CpG content are often protected from methylation, whereas DNA regions with low CpG content are often highly methylated [4, 10, 16].] The expression for *c_n_* accounts for our expectation that the correlation between the methylation states of two consecutive CpG sites will decay as the distance between these two sites increases, since the longer a DNMT enzyme must move along the DNA the higher is the probability of disassociating from the DNA before reaching the next CpG site.

### 2 Phase transition and methylation bistability

The ground state ***x**^*^* of the PEL *V_x_(**x**)* within a genomic unit (GU) is the state at which the potential energy attains its minimum value and represents the most likely methylation state within the GU. Interestingly, *V_x_(**x**)* can be characterized by co-existing ground states, depending on the values of the parameters *a_n_* and *c_n_* of the Ising model.

To see why this true, let us assume for simplicity that the Ising model within a GU will not be significantly affected if we replace the parameters *a_n_* and *c_n_* with their average values; i.e., if we set *a_n_* = *a* and *c_n_* = *c*, for all *n*, where

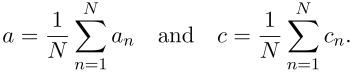

In this case, the PEL is approximately given by

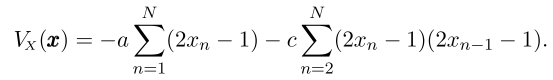

We can verify that, when *a* < 0 and *c* ≥ 0, the PEL’s minimum value will be *N(a − c)* and this value will be reached only at state ***x^*^*** = **0** (fully unmethylated state). On the other hand, when *a* > 0 and *c* ≥ 0, the PEL’s minimum value will be −*N*(*a* + *c*) and this value will be reached only at state ***x^*^*** = **1** (fully methylated state). However, when *a* = 0 and *c* > 0, the PEL’s minimum value will be −*N_c_*, and this value will be reached at *two* different states, 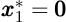 and 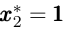. This means that the GU will most likely be in the fully unmethylated or the fully methylated state with equal probability. Taken together, these observations lead to an important result: when *c* > 0, the most likely state of the Ising model will experience (non-critical) phase transition as parameter *a* increases from negative to positive values with the transition occurring at *a* = 0. [^3^ Note, however, that this phase transition will be critical only in the limit as *c* → ∞ and *N* → ∞, since the ID Ising model does not experience criticality for finite values of *c* and *N*[1].].

In addition to the above, and as a direct consequence of well-known formulas of statistical mechanics that relate the magnetization of the ID Ising model with its underlying parameters (e.g., see [1, pp. 33-36]), we can show that, in the limit as *N* → +∞, the mean methylation level (MML) E *[L]* is given by

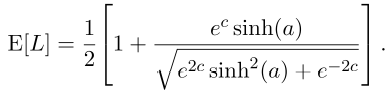

Although this is an asymptotic formula, it helps us gain a valuable insight into the behavior of methylation within a GU, even if the number of CpG sites is small. In particular, methylation within a GU can be subject to a (non-critical) phase transition that is reminiscent to the one that has been extensively studied in statistical mechanics; see Fig. Ml. It turns-out that this behavior is directly related to the potential wells of the PEL associated with the methylation level *L*, given by

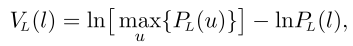

and leads to the observation that, for sufficiently large *c* > 0, the methylation level can be subject to bistability when the parameter *a* is close to the “critical” value 0; see Extended Data Fig. **6**. We attribute this property of the methylation system to a reallocation of the ground states of the PEL *V_L_*(*l*), caused by a biochemically-induced deformation of its topographic surface.

GUs exhibiting methylation bistability may be very important from a biological perspective. Under normal circumstances, bistability may be viewed as an efficient way to move from a fully unmethylated to a fully methylated state or vice versa. In other circumstances, this may be viewed as a biological aberration that may lead to disease. This fundamental epigenetic characteristic can be dynamically controlled by the methylation machinery, as well as by environmental factors, which may influence the particular values of parameters *α*, *β*, and *γ* Detecting bistable regions is possible due to the Ising model, which considers correlation in methylation in terms of parameter *c*.

**Figure Ml.**
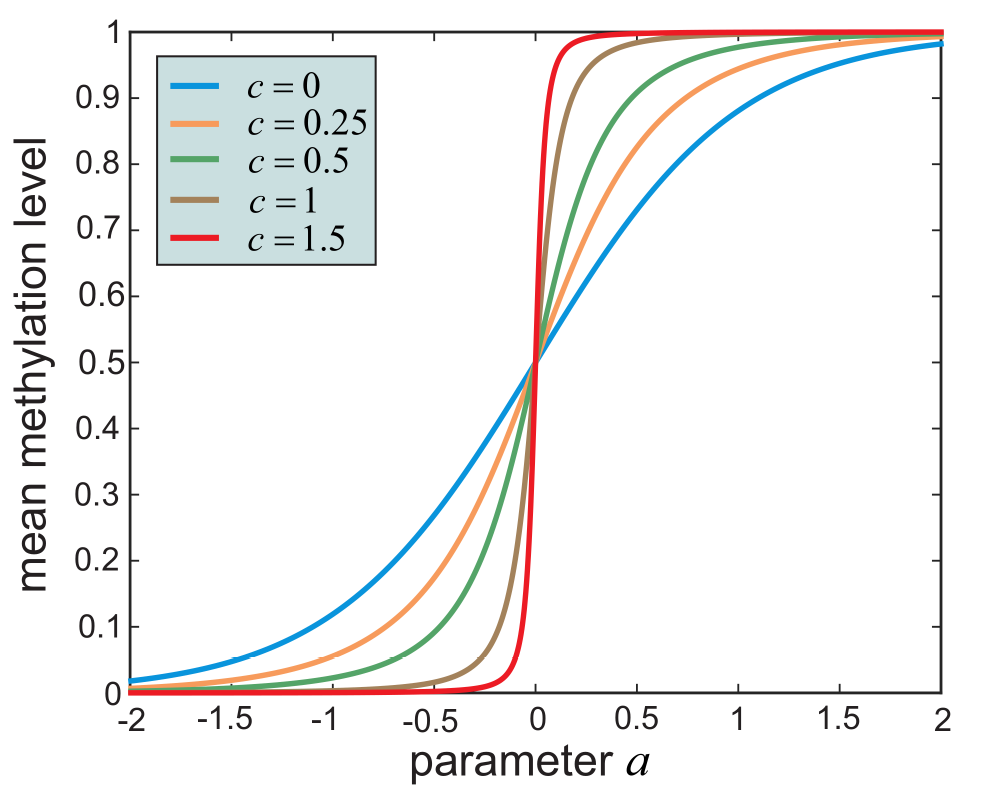
Mean methylation level (MML) of the Ising model within a genomic unit (GU) as a function of the PEL parameter *a* and for various values of the PEL parameter *c*. Note that a 50% MML is achieved if and only if *a* = 0. In addition, positive values of *a* arc associated with MMLs above 50%, whereas negative values of *a* are associated with MMLs below 50%. For small values of *c*, low MML can be achieved only with a value of *a* that is appreciably below zero, whereas high MML requires a value of *a* that is appreciably above zero. This however is not necessarily true for large values of *c*. In this ease, low and high MMLs can be achieved even with values of *a* elose to zero. Although the transition from MMLs below 50% to levels above 50%) is smooth for small values of *c*, this transition becomes sharper for larger values. In the latter cases, a small deviation in the value of *a* from zero can result in a sharp decrease of MML to 0 (fully unmethylated state) or a sharp increase to 1 (fully methylated state) demonstrating the presence of bistable behavior, a form of phase transition.

### 3 Maintenance of methylation information

In this paper, we use a noisy binary communication channel to model the transmission of the DNA methylation state during maintenance. We view this process as one that transmits methylation marks in the form of binary (0-1) bits of information at successive time points. We focus here on the fundamental notion of “methylation channels” and present information-theoretic tools that allow us to quantify the behavior of these channels. For related work on the maintenance of methylation marks, see [7–9, 12, 17].

#### 3.1 The methylation channel

We view the dynamic maintenance of the methylation state at the *n*-th CpG site of the genome as a multi-step transmission process modeled by a Markov chain *X_n_(0)* → *X_n_(l)* → … → *X_n_*(*k* − 1) → *X_n_*(*k*) → …, where *X_n_*(0) is the initial methylation state and *X_n_*(*k*) is the methylation state after the *k*-th step. In this case,

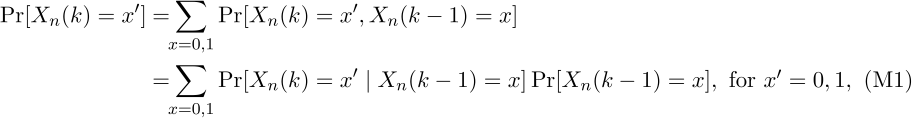

where Pr[*X*(*k*) = *x*′ | *X*(*k* − 1) = *x* is the probability of transmitting the methylation state *x* to the methylation state *x′* during the *k*-th step.

The probabilities *P_n_*(*x*′ | *x*) = Pr[*X_n_*(*k*) = *x′ | *X_n_*(*k* − 1) = *x*]* model the influence of *intrinsic* and *extrinsic* fluctuations to the process of methylation maintenance. These probabilities define an information transmission system, which we refer to as a methylation channel (MC) [^4^ In information theory, this system is known as noisy binary channel [5].]. Clearly, we can specify the MC associated with the *n*-th CpG site by using two parameters, *μ_n_* and *ν_n_*, such that

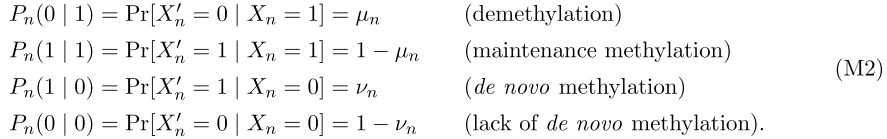

Parameter *μ_n_* quantifies the *net* rate of demethylation, which includes *passive* demethylation (due to lack of DNMT1 activity) as well as *active* demethylation (e.g., due to TET-mediated oxidation of methylated CpGs), whereas parameter *ν_n_* quantifies the *net* rate of methylation, which includes non-specific DNMT1 activity (usually very low), as well as *de novo* DNMT3-driven methylation – see [3, 11] for details.

Two special cases of the previous channel have been extensively studied in information theory: the binary symmetric channel, for which *μ_n_* = *ν_n_*, and the *Z*-channel, for which *ν_n_* = 0. Our treatment here is associated with the general case, which is known in information theory as the binary asymmetric channel. Note that we must limit the range of possible values for the demethylation and *de novo* methylation probabilities to 0 < *μ_n_* ≤ 1/2 and 0 < *ν_n_* ≤ 1/2. These inequalities are imposed by two factors: (a) demethylation and *de novo* methylation always occur in real cells and, therefore, *μ_n_,ν_n_* > 0, and (b) we expect demethylation to occur less frequently than maintenance methylation, in which case *μ_n_* ≤ 1 − *μ_n_*, and the same is expected to be true for *de novo* methylation.

#### 3.2 Estimation of methylation channels

To estimate a MC from WGBS data, we need to determine appropriate values for its parameters *μ_n_* and *ν_n_*. In general, the probabilities of demethylation and *de novo* methylation may vary during regulation of the methylation state. As a result, estimating a MC from data is a very difficult problem. However, we expect that stable conservation of the methylation state requires that the probabilities *μ_n_, ν_n_* of demethylation and *de novo* methylation, as well as the probabilities *P_n_*(1) of CG methylation, do not appreciably change with time [7, 9, 18], in which case the MC will operate near equilibrium. As a consequence, and by virtue of Eq. Ml, we expect to approximately have

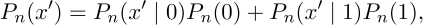

where *P_n_*(*x*) = Pr[*X_n_(k)* = *x* is the probability distribution of methylation at the *n*-th CpG site. This, together with Eq. M2, leads to

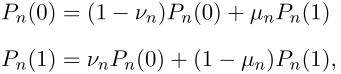

where *P_n_*(0) + *P_n_*(l) = 1. Therefore,

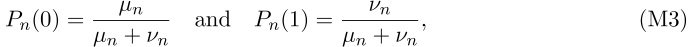

which implies that *Pn(* 1)

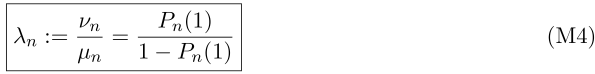

In this case, knowledge of the probability of methylation *P_n_*(1) allows us to compute the ratio *λ_n_* between the *de novo* and demethylation probabilities, which we refer to as the turnover ratio. It turns out that we can use WGBS data to estimate the value of the turnover ratio *λ_n_* at a CpG site *n*. To do so, we can use our maximum likelihood approach to estimate the Ising probability *P_x_(**x**)* and then marginalize this probability to obtain an estimate of *P_n_*(l).

#### 3.3 The average input/output information

The average information required to characterize the methylation state *X* at the input of a MC is given by the GpG entropy (CGE), defined by

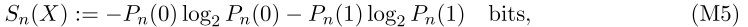

and likewise for the average information associated with the output *X*′. For the MCs considered here, we have *S_n_*(*X*) = *S_n_*(*X*′), since the input and output methylation probabilities are the same. It turns out that

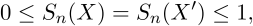

where the lower bound is achieved when *X* and *X*′ take value 0 or 1 with probability 1, and the upper bound is achieved when *X* and *X*′ take value 0 or 1 with equal probability. From Eqs. M3 – M5, the input/output CGEs are given by

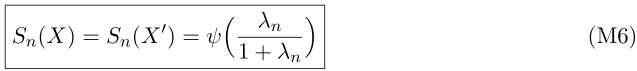

where

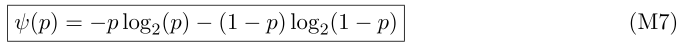

We can therefore compute these entropies from the turnover ratio *λ_n_* without specific knowledge of the probabilities *μ_n_, ν_n_* of demethylation and *de novo* methylation. Note that the CGE tends to zero as *λ_n_* → 0 or *λ_n_* → ∞ and monotonically increases to its maximum value of 1 as *λ_n_* → 1.

#### 3.4 The information capacity of a methylation channel

The information capacity (IC) of a MC expresses the maximum average information that can be conveyed during transmission. Mathematically speaking, the IC of a MC associated with the *n*-th CpG site is defined by

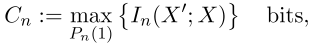

where *I_n_*(*X*′ ; *X*) is the mutual information between the output *X′* and the input *X* of the MC. The mutual information quantifies (in bits) the reduction in the average information required to characterize the output *X′* of the MC due to knowledge of its input *X*. It turns out that

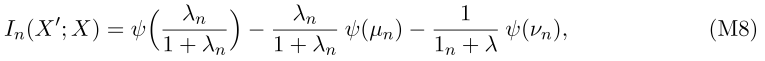

where *Ψ* is given by Eq. M7. Moreover, the IC is given by

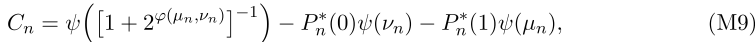

which is achieved by the input probability distribution

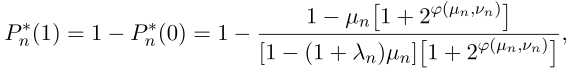

where

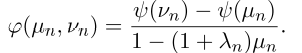

Finally,

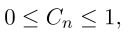

where the lower bound is achieved when *μ_n_* = *ν_n_* = 1/2, whereas the upper bound is achieved in the limit as *μ_n_* → 0 and *ν_n_* → 0.

Unfortunately, we cannot compute the IC from the turnover ratio *λ_n_*, since we also need to know the probability *μ_n_* of demethylation or the probability of *de novo* methylation *ν_n_*. However, we will shortly show that we can compute a reasonable approximation to this value.

#### 3.5 The probability of transmission error

A way to quantify the accuracy of methylation transmission through a MC is to calculate the probability of erroneously transmitting the input methylation state to the output. If the input *X* of a MC takes a value *x*, whereas its output *XℲ* takes a value *x′* ≠ *x*, then a transmission error occurs. In this case, the probability of error is given by

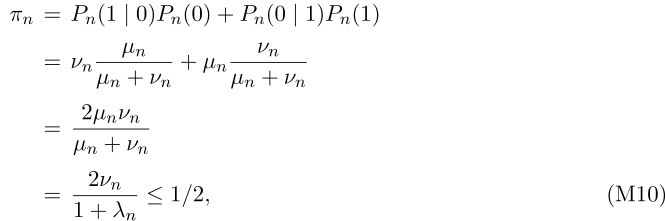

by virtue of Eqs. M2, Eq. M3 & Eq. M4. Note that the inequality is due to the fact that *μ_n_ ν_n_* ≤ 1/2, where equality holds if and only if *μ_n_* = *ν_n_* = 1/2.

#### 3.6 Energy dissipation

Information processing by a MC and, as a matter of fact, by any biological system, requires consumption of free energy. An amount of work is needed to correctly transmit the methylation state at a CpG site, and this consumes energy that is dissipated to the surroundings in the form of heat [2, 13]. Due to stochastic fluctuations in the underlying biochemistry, the methylation system always drifts towards imperfect transmission, characterized by a non-negligible probability of error. Under certain assumptions, it has been shown for some systems [6, 14] that the minimum energy *E* dissipated to the surroundings is related to the probability of error *π* by the following simple formula:

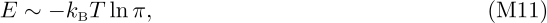

where *k_B_* is Boltzmann’s constant (*k_B_* = 1.3806488 × 10^−23^ JK^−1^) and *T* is the absolute temperature. We postulate that this relationship is also valid in the case of methylation transmission at a given CpG site, at least approximately.

Since the proportionality factor in Eq. Ml is not known, we define the relative dissipated energy (RDE) *ɛ_n_* by

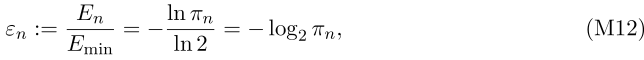

where *E_min_* ~ *k_B_T*ln 2 is the least possible energy dissipation (recall that *π_n_* ≤ 1/2). Note that *ɛ_n_* is a unitless quantity such that 1 ≤ *ɛ_n_* < ∞. Moreover, Eqs. M10 & Eq. M12 imply that

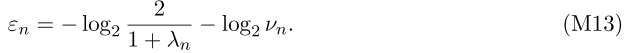

As a consequence, the RDE becomes infinitely large as *λ_n_* → 0 or *λ_n_* → ∞, whereas it takes value −log_2_*ν_n_*, when λ = 1. Notably, we cannot compute the RDE value from the turnover ratio *λ_n_*− for this calculation, we also need to know the probability of *de novo* methylation.

#### 3.7 Approximate computation of ICs and RDEs

We now discuss a method that allows us to approximately compute the ICs and RDEs genome-wide from the turnover ratios *λ_n_*. Despite its approximative nature, the method leads to a number of fundamental insights into the information-theoretic properties of methylation maintenance. Our approach can become more precise once methods for estimating the probabilities of demethylation and *de novo* methylation become available.

The probabilities of demethylation and *de novo* methylation are expected to be small [79, 15]. For this reason, we assume that 0 ≤ *μ_n_* ≤ 0.1 and 0 ≤ *ν_n_* ≤ 0.1 at each CpG site. In addition, since it is not possible to determine the exact values of the demethylation and *de novo* methylation probabilities within a cell population, we may consider via the principle of maximum entropy that these probabilities are random variables that follow a *uniform* distribution within the closed interval [0, 0.1]. In this case, the mean values of *μ_n_* and *ν_n_* within the cell population will be given by 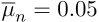 and 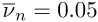.

We can now derive a formula for approximately calculating the IC values within the cell population, which we can use instead of the true values. To do so, we approximate the function *Υ*(*p*) given by Eq. M7, using the following *linear* approximation: [^5^ If we assume that 0 ≤ *p* ≤ 0.1, then we can set *Ψ(p) ≃ Kp*. It turns out that *K* = 5.19 minimizes the square error between *Ψ(p)* and *kp*.]

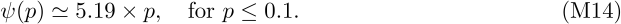

Let us consider a CpG site for which *λ_n_* ≤ 1. Figure M2 shows that the capacity *C_n_* is almost identical to the ratio *I_n_*(*X*′ ; *X*)/*S_n_*(*X*), known as channel transmissibility. This leads to

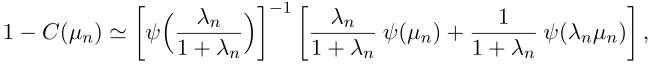

by virtue of Eqs. M6 – Eq. M9, where we now explicitly denote the dependence of the IC on *μ_n_*. As a consequence of Eq. M14, we obtain

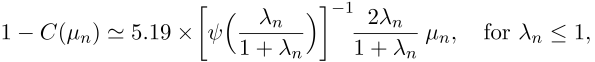

since *μ_n_* ≤ 0.1 and λ*_n_μ_n_* ≤ 0.1 in this case. On the other hand, when *λ_n_* ≥ 1, we have that

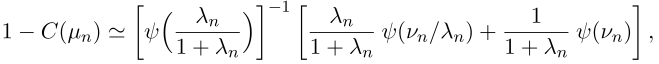

which implies

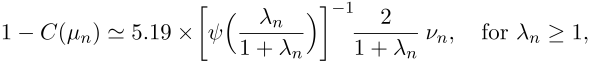

>
since *ν_n_* ≤ 0.1 and *ν_n_*/*λ*_n_, ≤ 0.1 in this case. By taking expectations, and since 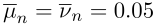, the previous results lead to

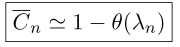

with

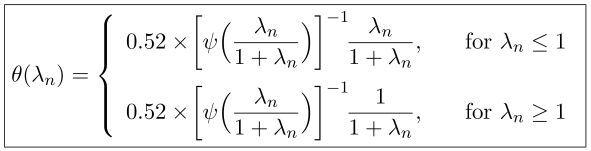

where is 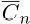 the mean value of the IC within the cell population. Since these formulas depend only on the turnover ratio λ_n_, they allow us to approximately compute the IC values genome-wide from WGBS data.

We can also derive a formula for approximately calculating the RDE values within the cell population. From Eq. M13, we can show that the mean value 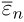 of the RDE at a CpG site with turnover ratio *λ_n_* is given by

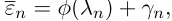

where

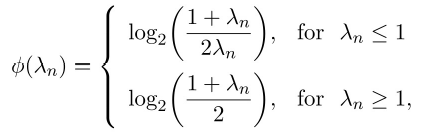

and

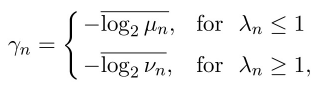

with 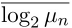 and 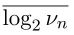 being the mean values of the logarithms of the probabilities of demethylation and *de novo* methylation. It turns out that, since we assume that *μ_n_* follows a uniform distribution within the closed interval [0,0.1], we have −log_2_*μ_n_* = −log_2_ 0.1 + 1.44 × ɛ = 3.32 + 1.44 × ɛ, where ɛ follows an exponential distribution with unit rate. As a consequence, 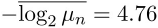 and, the same is true for 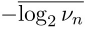 We therefore obtain

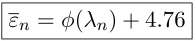

where

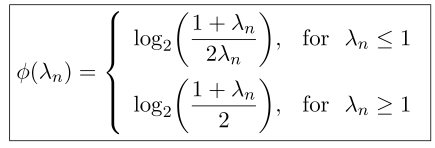

**Figure M2.**
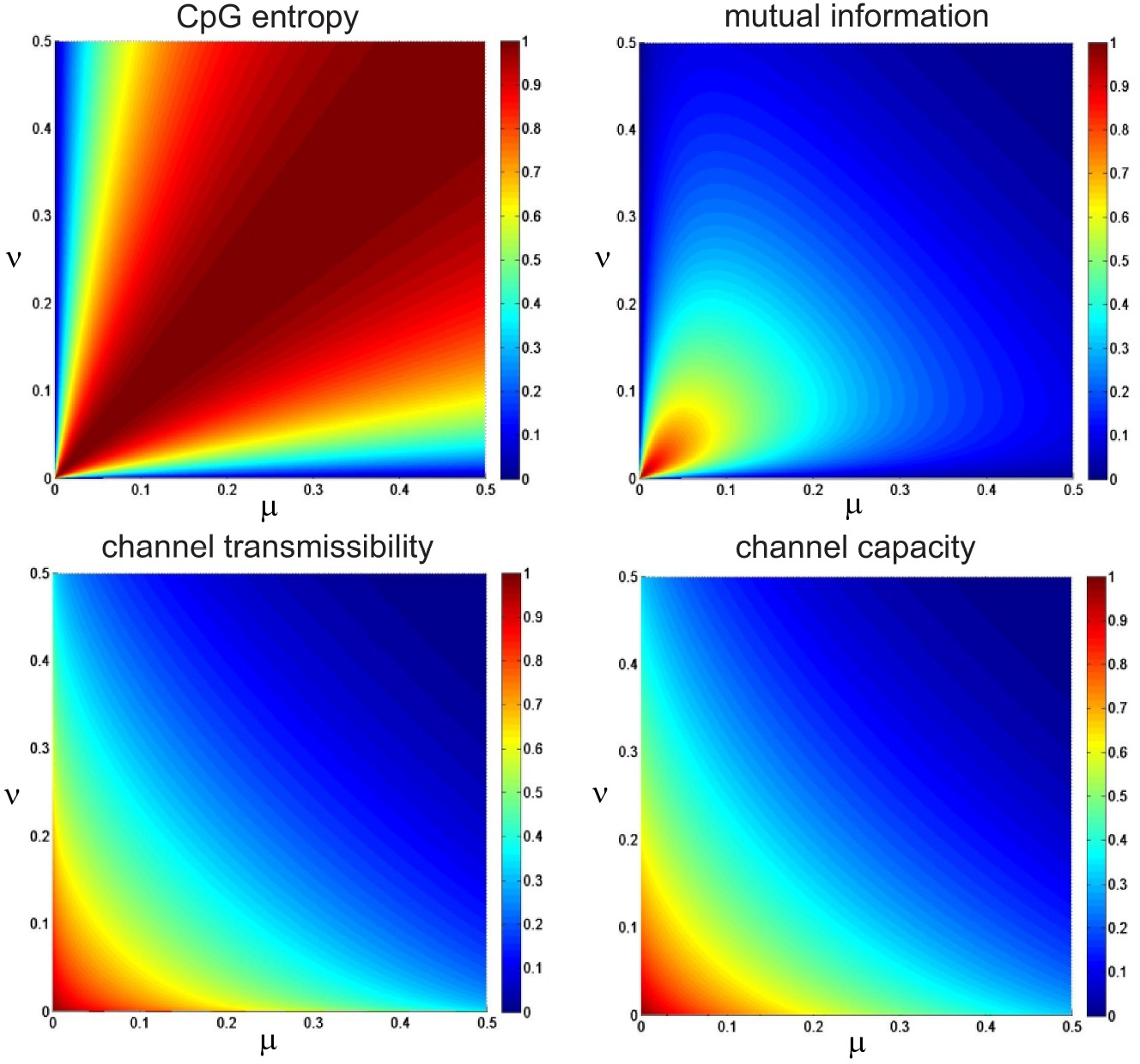
CpG entropy *S*(*X*), mutual information *I*(*X*′ *X*), channel transmissibility *I*(*X*′ *X*)/*S*(*X*), and channel capacity *C* of a MC as a function of the demethylation and *de novo* methylation probabilities *μ* and *ν* Note that the channel capacity is almost identical to the transmissibility.

Since these formulas depend only on the turnover ratio *λ_n_*, they allow us to to approximately compute the RDE values genome-wide from WGBS data.

### 4 The Entropic Sensitivity Index

To investigate the influence of environmental conditions on methylation uncertainty, we postulate that changes in environmental conditions affect the parameters *α, β*, and *γ* of the Ising model and derive a measure that allows us to quantify the effect of parameter variation on the NME. We consider the case for which the Ising parameters *α, β*, and *γ* fluctuate around their true values by random amounts *αG, βG*, and *γG*, respectively. We model *G* as a random variable that follows a zero-mean normal distribution with small standard deviation *σ* Moreover, we hypothesize that the NME *h(g)*, produced by the perturbed Ising model with parameters (1 + *g*)*α*, (1 + *g*)*β* and (1 + *g*)*γ*, is a sufficiently smooth function of *g* so that it can be approximated by the following second-order Taylor series expansion:

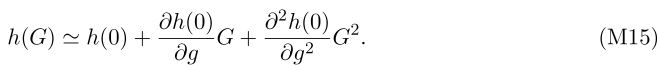

Since *σ* is small and *h* is thought to be a sufficiently smooth function of *g*, we expect that

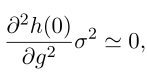

whereas, since *G* follows a normal distribution, we have that E[*G*^3^] =0. As a consequence, we can show from Eq. M15 that the standard deviation *σ_h_* of the entropy is approximately given by

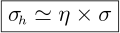

where

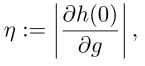

which we refer to as the entropic sensitivity index (ESI).

Evaluating the ESI requires approximating the derivative. Using a finite-difference approximation, we set

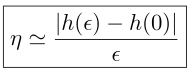

for some small *ɛ*. In turn, this requires that we compute the NME *h*(0) from the “true” Ising model with parameters 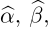, and 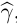, which we estimate from given WGBS data, as well as the NME *h*(*ɛ*) from the “perturbed” Ising model with parameters 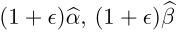, and 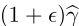. In our calculations, we set *ɛ* = 0.01.

## Supplementary Notes

### 1 Critique of empirically estimating epiallelic probabilities

A number of methylation analysis methods have been recently proposed in [24] using the notions of epipolymorphism as well as combinatorial and informational entropies. These statistics are defined by the probability distribution of *2^K^* possible methylation patterns, known as epialleles, observed within a window of *K* ≥ 4 contiguous CpG sites. However, we question the legitimacy of these methods on the ground that they rely on *empirically* estimating the epiallelic probabilities by using the ratio *N_i_/N*, where *N_i_* is the number of times the *i*-th pattern is observed (epiallelic occurrence) in *N* methylation reads (coverage). This practice is highly questionable from a statistical point of view as we show next.

Let us consider the occurrence number *N_i_* of the *i*-the epiallele in *N* methylation reads to be the number of positive outcomes of *N* independent and identically distributed Bernoulli trials with probability of success *p_i_*, where *p_i_* is the true probability of occurrence of the *i*-th epiallele. In this case, the distribution of *N_i_* when 
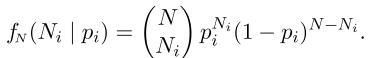

By assuming a uniform prior for the true epiallelic probability *p_i_*, we obtain

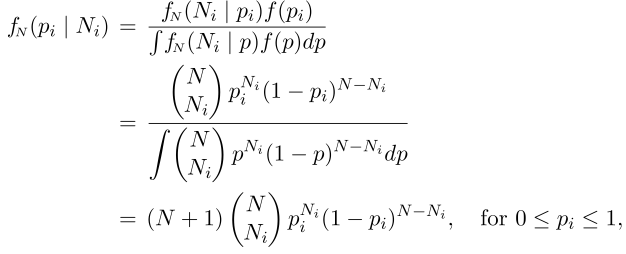

for the posterior distribution *f_N_*(*p_i_* | *N_i_*), which is a beta distribution with parameters *N_i_* + 1 and *N* − *N_i_* + 1.

To evaluate the uncertainty associated with the true value *p_i_* of the *i*-th epiallelic probability when it is empirically estimated by the ratio *N_i_*/*N*, we can compute the 95% (highest posterior density) credible interval associated with the conditional distribution of *p_i_* given *N_i_*, which contains the true value of the epiallelic probability with 95% likelihood. We can calculate this interval by setting

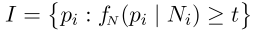

where *t* satisfies the following equation:

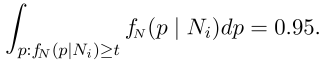

We can then use the width of the credible interval to measure uncertainty about the true epiallelic probability, with a wider interval indicating higher uncertainty.

In [4], the authors combined unique molecular identifiers with reduced representation bisulphite sequencing and constructed methylation data within selected regions of the genome exhibiting 20x coverage, which they included for downstream analysis. Moreover, by using WGBS data from human somatic tissues, they identified genomic regions with at least 14x coverage and used these regions to demonstrate heterogeneity in methylation. In Fig. Nl(a,b), we depict the credible intervals that correspond to these two cases. The results clearly demonstrate the flawed nature of empirically estimating epiallelic probabilities from a small number of methylation reads. For example, an epiallelic occurrence of 2 in data with 14x coverage implies an empirically estimated probability value of 0.14. However, Fig. Nl(a) shows that, with 95% likelihood, the value of the true epiallelic probability can be anywhere between 0.03 and 0.38, which demonstrates the large uncertainty and low accuracy of the empirically estimated value. This can be a major problem, since an epiallelic pattern is defined in [4] as being “noise” if the probability of occurrence is less than 0.2. It turns out that, by using this method, it may not be possible to statistically distinguish “noise” from “signal.” Likewise, an epiallelic occurrence of 7 in data with 14x coverage implies an empirically estimated probability value of 0.5 and a credible interval that would span a wide range of true probability values from 0.26 to 0.73.

We expect that the geometric growth of the number of epiallelic patterns containing an increasing number of CpG sites will worsen the previous problem. A good example is the genomic locus chrll: 2,021,946-2,022,065 depicted in Fig. 4b of [4], which comprises 10 CpG sites. In this case, 1,024 epiallelic probabilities are empirically estimated from 20 observations. Due to insufficient data, however, the value of at least 1,004 of these probabilities must be set equal to zero, even if the true values of these probabilities are not zero.

With such uncertain and inaccurate estimates, the previous problem extends well beyond the issue of defining epiallelic patterns that constitute “noise,” since the error incurred may significantly affect downstream analysis when using epipolymorphisms or entropies. To demonstrate the significance of this issue, we used the method in [4] and estimated these quantities in a clonal population under varying levels of pattern heterogeneity. By following [4], we considered a clonal population that included two main epiallelic patterns, each with probability of occurrence (1 − *π*)/2, where *π* is the net probability of occurrence of the remaining “noise” patterns, which we assume to occur with equal probability. We then used *π* to quantify the level of pattern heterogeneity in the population.

We considered a genomic region with 10 CpG sites, such as the one depicted in Fig. 4b of [4], within a clonal population that includes two main epiallelic patterns, each with probability of occurrence (1 − *π*)/2, as well as 1,022 “noisy” patterns, each with probability of occurrence *π*/l,022. We then analytically calculated, for a given level of pattern heterogeneity, its epipolymorphism *Q* and informational entropy *H* by

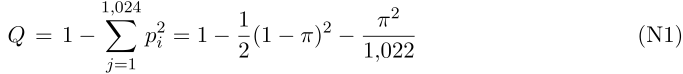

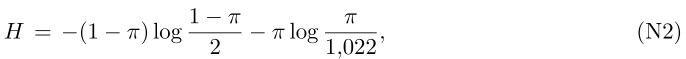

which imply that 0.5 ≤ *Q* ≤ 0.9990 and 0.8115 ≤ *H* ≤ 6.9295. For a given value of *i*r, we subsequently employed Monte Carlo sampling and independently generated 2,000,000 epiallelic patterns, which we grouped into sets of 20. We then empirically estimated the epiallelic probabilities within each set and calculated the epipolymorphism and entropy values using Eqs. NI & Eq. N2. We finally used these values to compute the 95% confidence intervals associated with the underlying empirical estimators by finding the 2.5-th and 97.5-th percentiles of the resulting 100.0 samples of the epipolymorphism and of the 100,000 samples of the entropy. The locations and widths of these confidence intervals can serve as measures of estimation accuracy. The results depicted in Fig. Nl(c,d) clearly demonstrate that empirical estimation of epipolymorphisms and entropies is prone to large errors and strongly support our view that this practice is highly questionable and not appropriate for reliable downstream analysis of methylation data.

**Figure N1.**
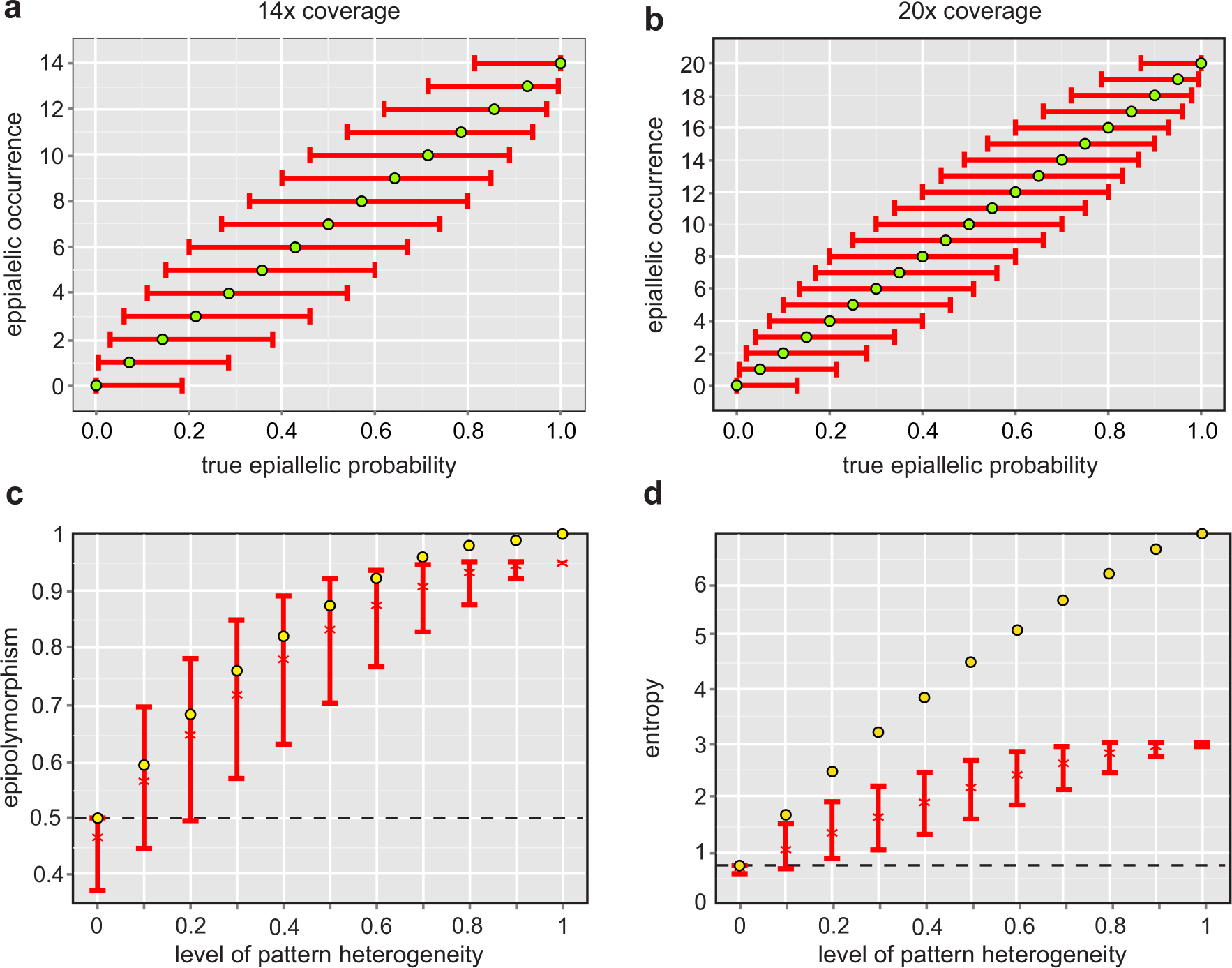
**a**, Empirically estimated probabilities (14x coverage) of an epiallelic pattern for different epiallelic occurrences (green circles) and the corresponding 95% high posterior density credible intervals (red error bars), **b**, Empirically estimated probabilities (20x coverage) of an epiallelic pattern for different epiallelic occurrences (green circles) and the corresponding 95% high posterior density credible intervals (red error bars). The widths of the credible intervals in **a** & **b** indicate appreciable uncertainty in the empirically estimated probability values, with maximum uncertainty occurring when the epiallelic pattern is observed in half of the available methylation reads, **c**, True values of the epipolymorphism (yellow circles) of a genomic region within a clonal population that contains 10 CpG sites with different levels of pattern heterogeneity and the corresponding 95% confidence intervals (red error bars) associated with empirically estimating the epipolymorphism. Although empirical estimation of the epipolymorphism leads, on the average, to values (red crosses) that arc relatively close to the true values, the estimator is associated with confidence intervals that may extend below this example’s allowable minimum value of 0.5 (dotted line), or with confidence intervals that do not include the true value at high levels of pattern heterogeneity, **d**, True values of the entropy (yellow circles) of the same genomic region as in **c** and the corresponding 95%) confidence intervals (red error bars). The empirical entropy estimator leads to completely inaccurate results since, in the absence of pattern heterogeneity, the resulting confidence interval extends below this example’s allowable minimum value of 0.8115 (dotted line), whereas, in the presence of pattern heterogeneity, the confidence intervals do not contain the true entropy values.

Empirical estimation of epiallelic probabilities was originally proposed in [2] together with an experimental method producing methylation data with an extraordinary coverage of about 10.0 x, which alleviates the previous problem. Unfortunately, this method can only be used to probe a very small number of genomic regions and is associated with a separate host of issues, such as the inability to distinguish PCR duplicates from truly replicated data [5]. To address the PCR duplication problem, UMI filtering was used in [4] but did not consider sufficient depth for meaningful epiallelic analysis.

In conclusion, extensive use of empirically estimating epiallelic probabilities and of other quantities defined from these probabilities, such as epipolymorphisms and entropies, call into question many analysis results and biological conclusions presented in [2, 4]. In particular, a number of predictions in [4] based on epiallelic patterns classified as “noise” should be retracted until statistically significant results can be presented in support of these conclusions, since we have convincingly demonstrated in this section that “noise” cannot be statistically distinguished from “signal” when using an empirical approach on small sample sizes. This turns out to be a much lesser problem when using the Ising model proposed in this paper, since the underlying parametric assumptions reduce the burden of statistical estimation from *2^K^* parameters [i.e., the *2^K^* probabilities *P*(*x_n_,x_n_*+1, … *x_n+K-1_*)] in the case of epipolymorphism, to the three *α, β, γ* parameters in the case of our model. In addition, our method can estimate joint probability distributions over regions that contain many more than 4 contiguous CpG sites, using methylation data of lower coverage than the one considered in [2, 4].

### 2 Estimation of TAD boundaries (additional results)

We now provide additional results regarding our method for estimating TAD boundaries. Understanding these results requires familiarity with the GenometriCorr package [1].

#### 2.1 HI stem predictive regions vs. HI stem TAD boundaries

GeometriCorr employs a number of output measures for evaluating the correlation of genome-wide data with given genomic features. Our analysis produced the following values for these
measures:

| GenometriCorr Output |  |
| --- | --- |
| reference.population | 5,862 |
| query.population | 404 |
| relative.distances.ks.p.value | $4.83 \times 10^{-8}$ |
| relative.distances.ecdf.area.correlation | $1.82 \times 10^{-1}$ |
| projection.test.p.value | 0 |
| projection.test.lower.tail | FALSE |
| relative.distances.ecdf.deviation.area.p.value | $< 2 \times 10^{-4}$ |
| scaled.absolute.min.distance.sum.p.value | $< 2 \times 10^{-4}$ |
| jaccard.measure.p.value | $< 2 \times 10^{-4}$ |
| scaled.absolute.min.distance.sum.lower.tail | TRUE |
| jaccard.measure.lower.tail | FALSE |

The value of “reference.population” in the table above indicated that there were 5,862 annotated TAD boundaries in available TAD annotations for HI stem cells. The value of “query.population” indicated that there were 404 predictive regions that may contain TAD boundaries. The very low P value calculated in “relative.distances.ks.p.value,” together with the observation that “relative. distances.ecdf.area.correlation” was positive, showed that, in general, the TAD boundaries and the predictive regions were closer to each other than expected by chance. The “projection.test.p.value” was zero, indicating either significant overlap or significant lack of overlap. The “projection.test.lower.tail” was FALSE, meaning that there was significantly more overlap between the predictive regions and the TAD boundaries than by chance. The P values given by “relative.distances.ecdf.deviation.area.p.value,” “scaled.absolute.min.distance.sum.p.value,” and “jaccard.measure.p.value’ indicated that the observed spatial relationships (absolute or relative distance apart) were significantly different than what was expected by chance. From these P values we could tell whether the predictive regions and the TAD boundaries were significantly close or significantly far apart. Since the value of the “scaled.absolute.min.distance.sum.lower.tail” was TRUE, we knew that the absolute distances between the predictive regions and the TAD boundaries were consistent and small. On the other hand, the “jaccard.measure.lower.tail” was FALSE, indicating an unexpectedly high overlap, as defined by the Jaccard measure. We should finally mention that the relative distance test of GenometriCorr revealed that true TAD boundaries that fell outside predictive regions were significantly more likely to be closer to these regions than expected by chance. We therefore concluded that all tests in this analysis indicated that TAD boundaries were located within predictive regions or were close to these regions in a statistically significant manner.

#### 2.2 Combined predictive regions vs. combined TAD boundaries

In this case, our analysis produced the following values for GeometriCorr measures discussed above:

| GenometriCorr Output |  |
| --- | --- |
| reference.population | 10,276 |
| query.population | 6,632 |
| relative.distances.ks.p.value | 0 |
| relative.distances.ecdf.area.correlation | $1.28 \times 10^{-1}$ |
| projection.test.p.value | 0 |
| projection.test.lower.tail | FALSE |
| relative.distances.ecdf.deviation.area.p.value | $< 2 \times 10^{-4}$ |
| scaled.absolute.min.distance.sum.p.value | $< 2 \times 10^{-4}$ |
| jaccard.measure.p.value | $< 2 \times 10^{-4}$ |
| scaled.absolute.min.distance.sum.lower.tail | TRUE |
| jaccard.measure.lower.tail | FALSE |

These values were similar to the ones obtained when using only HI stem cells, thus leading to the same conclusion as before. However, 6,632 predictive regions were correlated with 10,276 annotated TAD boundaries, as compared to correlating 404 predictive regions with 5,862 annotated TAD boundaries in the case of HI stem cells.

